# Lamellar Normative Modelling of the Hippocampus Across the Human Lifespan

**DOI:** 10.64898/2026.05.02.722431

**Authors:** Yanwu Yang, Na Gao, Nitin Sharma, Sicheng Dai, Guinan Su, Richard Dinga, Pierluigi Selvaggi, Jens Krüger, Yue Cai, Li Liang, Zhiyuan Liu, Saige Rutherford, Tengfei Guo, Thomas Wolfers, Alzheimer’s Disease Neuroimaging Initiative

## Abstract

The hippocampus is a central hub of human memory and cognition and is closely associated with brain disorders. Studies have shown that it exhibits complex structural variation across the lifespan, yet the details of hippocampal morphology changes remain poorly understood. Here, we establish norms over the hippocampal geometry that resolve lamellar morphology and map lifespan trajectories across more than 27,000 individuals from 158 scanning sites. Hippocampal geometry shows spatially non-uniform developmental and ageing patterns across the lifespan, with lamellar thickness, width and length following dissociable trajectories. Across multiple brain disorders, this representation reveals localized and heterogeneous alterations beyond conventional subfield-level summaries, and uncovers a dichotomy in disease-associated patterns, with neurodegenerative conditions and schizophrenia showing predominant atrophy, whereas some other disorders exhibit focal or regionally selective hypertrophy. Transfer to a longitudinal Alzheimer’s Disease Neuroimaging Initiative cohort further supports out-of-sample generalization of our approach and enables individual-level tracking and conversion risk stratification. Overall, this work establishes a population-scale geometric reference for the hippocampus, extends normative brain mapping from coarse regional phenotypes to anatomically organized subcortical structure, and enables anatomically grounded characterization of disease-related alterations and individual-level deviation mapping, providing a principled basis for understanding and stratifying brain disorders across the lifespan in health and disease.

## Introduction

Brain charts provide population-level reference trajectories of brain structure across the lifespan^1–3^, enabling deviation mapping at the individual level that links typical ageing to neurodevelopmental and neurodegenerative processes. Normative modelling extends this framework by explicitly capturing inter-individual variability and quantifying disease-related departures from typical trajectories^3–5^. However, most existing approaches rely on global or regionally aggregated phenotypes, which obscure spatially localized variation within anatomically complex structures and limit sensitivity to fine-grained morphological heterogeneity^6–8^.

This limitation is particularly critical for brain systems with intrinsic internal organization^9–11^. The hippocampus represents a prominent example, given its central role in memory and its involvement in a broad range of cognitive functions, including learning and spatial representation^9,12,13^. Rather than forming a homogeneous structure, the hippocampus exhibits continuous structural variation along multiple anatomical axes, paralleling well-established gradients of functional organization, particularly along the anterior-posterior (long) axis and the skeletal-lateral dimension defined by its intrinsic geometry^14–16^. Converging evidence from anatomical, electrophysiological, genetic and high-resolution imaging studies supports heterogeneous organization along the longitudinal axis^17–19^. Consistent with this view, in vivo neuroimaging studies have revealed region-specific vulnerability of the hippocampus to ageing and disease^19–21^, reinforcing the intrinsically non-uniform nature of hippocampal change across the lifespan^22–24^.

Recent advances in normative modelling have enabled robust lifespan charting of brain structure and provided a powerful reference for quantifying individual-level deviations^25^. However, extending these approaches to the fine-grained organization of the hippocampus remains challenging in practice. A major challenge lies in establishing anatomically consistent representations that allow direct comparison across individuals while preserving continuous variation along the long axis. Existing approaches to hippocampal morphology typically rely on subfield volumetry or surface-based analyses^26–29^, which impose discrete boundaries or lack intrinsic correspondence along the longitudinal dimension, thereby limiting sensitivity to axis-aligned variation. At the lamellar level, lamellae are defined as thin slices perpendicular to the anatomical long axis and uniformly distributed along it. This representation, grounded in the intrinsic longitudinal organization of the hippocampus, provides a principled basis for establishing structural correspondence across individuals^21,30^. Building on this concept, we employed axis-referenced morphometric modelling (ARMM) ^29,31^, a skeletal-based framework that defines an intrinsic coordinate system along the curved long axis from the tail apex to the skeletal aspect of the head. This framework discretizes the hippocampus into anatomically aligned lamellae and enables quantitative measurement of lamellar thickness, width and long-axis length.

Here, we establish a unified geometric normative charting framework for the hippocampus across the human lifespan. By leveraging ARMM-based lamellar representations, we characterize hippocampal geometry in a large population dataset of more than 27,000 individuals and derive population-scale reference trajectories of lamellar thickness, width and long-axis length. Across a broad spectrum of brain disorders, including autism spectrum disorder (ASD), attention-deficit/hyperactivity disorder (ADHD), anxiety disorders (ANX), major depressive disorder (MDD), schizophrenia (SCZ), mild cognitive impairment (MCI) and dementia (DM), we show that hippocampal geometry exhibits temporally dissociated and spatially organized lifespan trajectories, with distinct geometric components capturing separable processes of maturation and degeneration. Across disorders, deviations form disease-specific, spatially organized patterns that are captured with high anatomical precision within the lamella-based framework, enabling improved sensitivity to disease-related alterations and enhanced discriminative performance. Generalization to an independent longitudinal ADNI cohort further supports the utility of hippocampal geometry for tracking disease progression and stratifying individual-level conversion risk.

## Results

### Lifespan trajectories of lamella-based hippocampus morphology

We quantified hippocampal geometry using lamella-resolved morphometric features derived from an axis-referenced modelling framework (Fig. 1a), capturing lamellar thickness, width and long-axis length within an anatomically consistent coordinate system^31^. The ARMM representation demonstrated improved surface reconstruction accuracy, point-wise correspondence and measurement reproducibility relative to alternative shape models^32^ (Supplementary Figs. S1-S3 and Supplementary Table S1-S3). A large-scale lifespan cohort comprising 27,069 individuals was assembled (Table 1; Fig. 1b), of whom 20,746 participants without clinical diagnoses were assembled to construct normative models spanning childhood to late adulthood. Normative trajectories were estimated using generalized additive models for location, scale and shape (GAMLSS) ^33^, modelling age-dependent variation while accounting for sex and site effects (see Methods). Model validation and evaluation of alternative GAMLSS specifications are detailed in the Supplementary Table S7, Figs. S4-S9.

**Fig. 1.**
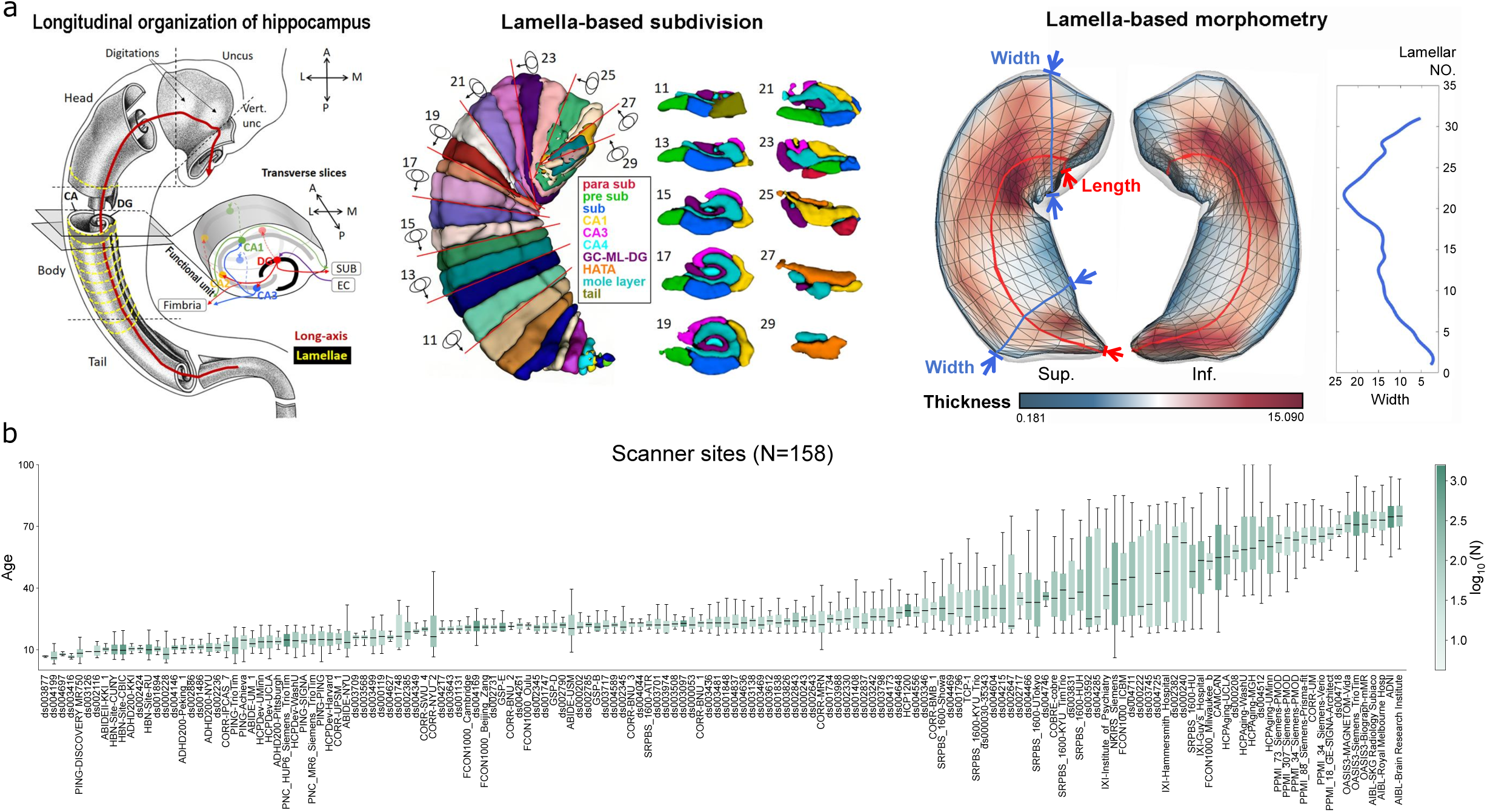
Geometry-aware framework for lamella-based representation of hippocampal morphology across the lifespan. a, Schematic overview of the hippocampal geometric mapping. The organization of the hippocampus is defined relative to its intrinsic lamellar architecture, enabling anatomically consistent characterization of morphology along the long axis. Axis-referenced morphometric modelling further subdivides the hippocampal surface into lamellar units and quantifies complementary geometric features, including lamellar thickness, width, and long-axis length, thereby capturing fine-grained internal structural variation beyond conventional regional summaries. b, Distribution of the curated cohort across scanner sites used in this study. Boxplots show the age distribution at each site, ordered across the lifespan, with color indicating sample size on a log-scaled axis. The reference dataset comprises 158 scanner sites and spans a broad age range, providing the population-scale basis for modelling hippocampal geometry across development, adulthood, and ageing.

**Table 1.**
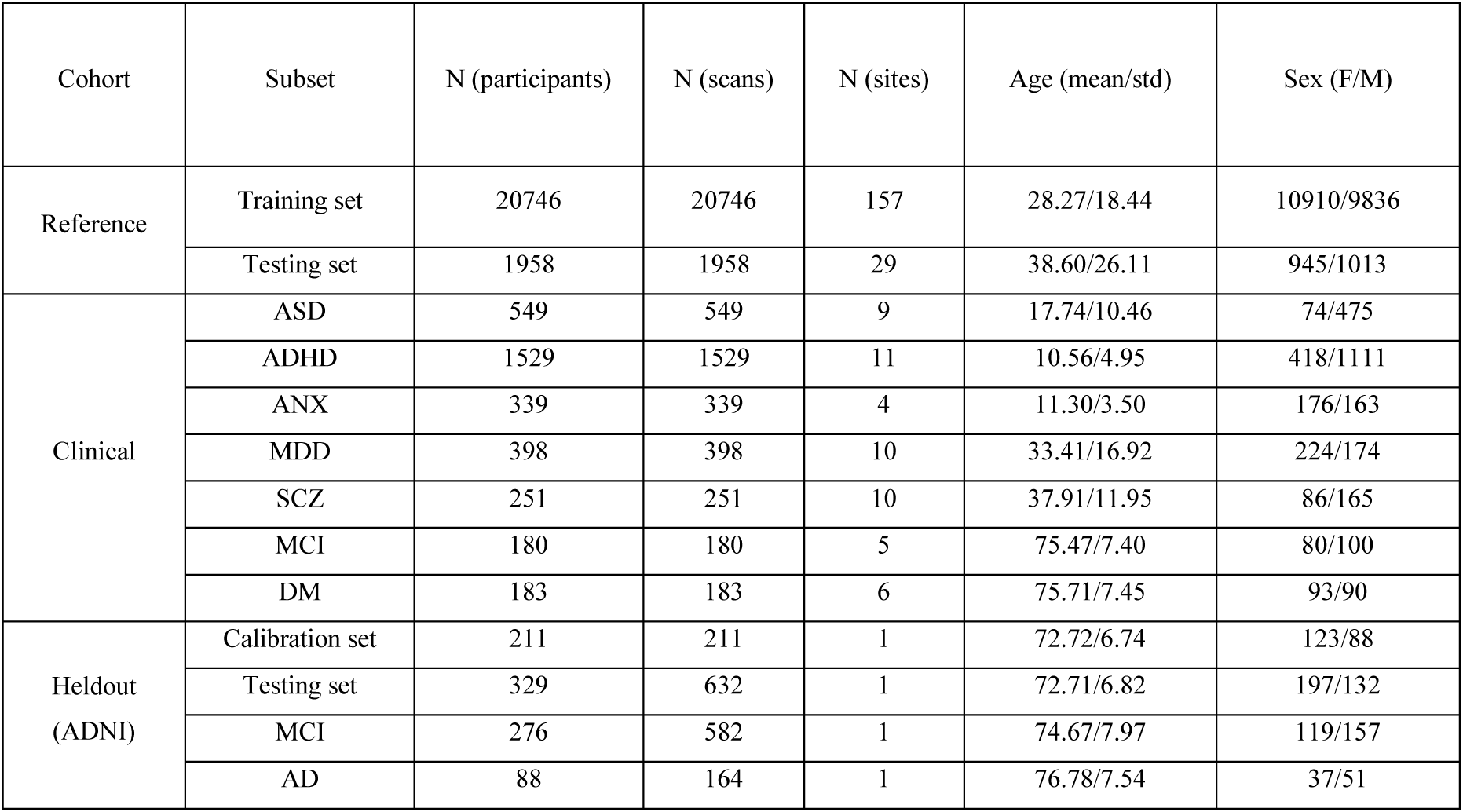
Sample description and demographics.

| Cohort | Subset | N (participants) | N (scans) | N (sites) | Age (mean/std) | Sex (F/M) |
| --- | --- | --- | --- | --- | --- | --- |
| Reference | Training set | 20746 | 20746 | 157 | 28.27/18.44 | 10910/9836 |
|  | Testing set | 1958 | 1958 | 29 | 38.60/26.11 | 945/1013 |
| Clinical | ASD | 549 | 549 | 9 | 17.74/10.46 | 74/475 |
|  | ADHD | 1529 | 1529 | 11 | 10.56/4.95 | 418/1111 |
|  | ANX | 339 | 339 | 4 | 11.30/3.50 | 176/163 |
|  | MDD | 398 | 398 | 10 | 33.41/16.92 | 224/174 |
|  | SCZ | 251 | 251 | 10 | 37.91/11.95 | 86/165 |
|  | MCI | 180 | 180 | 5 | 75.47/7.40 | 80/100 |
|  | DM | 183 | 183 | 6 | 75.71/7.45 | 93/90 |
| Heldout<br>(ADNI) | Calibration set | 211 | 211 | 1 | 72.72/6.74 | 123/88 |
|  | Testing set | 329 | 632 | 1 | 72.71/6.82 | 197/132 |
|  | MCI | 276 | 582 | 1 | 74.67/7.97 | 119/157 |
|  | AD | 88 | 164 | 1 | 76.78/7.54 | 37/51 |

We first summarized hippocampal morphology at a global level by averaging lamellar thickness and width within the head, body and tail, along with long-axis length (Fig. 2a). These features exhibited distinct, regionally heterogeneous trajectories across the lifespan. Long-axis length peaked in early adulthood (30.3 years). Lamellar width reached its maximum earlier, peaking at 27.9, 20.5 and 14.2 years in the head, body and tail, respectively. In contrast, lamellar thickness showed more protracted trajectories, peaking at 40.5 years in the head, 59.1 years in the body, and 29.3 years in the tail. These results reveal a temporal dissociation across geometric features, with width and length peaking earlier than thickness across hippocampal subdivisions. Figures 2b and 2d illustrate the spatial distribution of explained variance for lamellar thickness and width, respectively. Overall, the proportion of variance explained by age is modest (ranging from 0 to 0.2) yet exhibits clear spatial structure. Stronger age-related effects are observed in dorsal regions for lamellar thickness and in the body and tail for lamellar width. Complementary visualization (Supplementary Video) further demonstrates that lifespan-related morphological changes are primarily characterized by curvature alterations along the long axis, particularly at the anterior and posterior ends, and by progressive thinning in posterior regions.

**Fig. 2.**
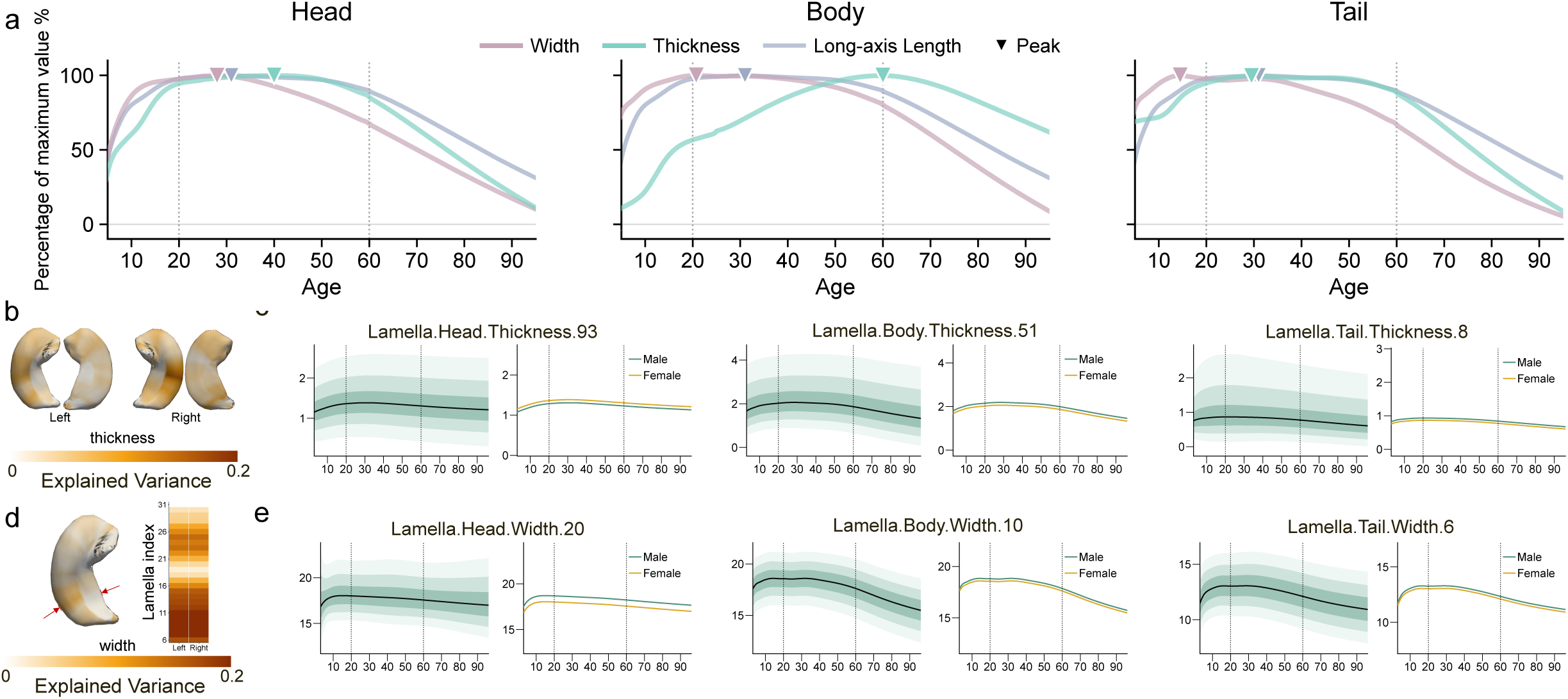
Lifespan trajectories of hippocampal geometric features reveal regionally heterogeneous patterns of maturation and ageing. a, Lifespan trajectories of global hippocampal geometry across the head, body and tail, shown as normalized values relative to the maximum for each feature. Width, thickness and long-axis length exhibited distinct age-related profiles, indicating that different geometric properties of the hippocampus follow non-uniform developmental and ageing trajectories across anatomical subdivisions. Inverted triangles denote the estimated ages at peak values. b, Spatial maps of explained variance for lamella-based thickness in the left and right hippocampus. c, Representative lifespan trajectories of lamellar thickness in the head, body and tail, with shaded regions indicating normative centiles (5th, 25th, 50th, 75th and 95th percentiles) and corresponding sex-specific differences. d, Spatial maps of explained variance for lateral-skeletal width in the left and right hippocampus. e, Representative lifespan trajectories of lamellar width in the head, body and tail, shown separately for males and females. Similar to thickness, width displayed marked regional heterogeneity, supporting a fine-grained view of hippocampal morphological change across the lifespan.

We next investigated normative trajectories of lamella-wise thickness and width, with each lamella annotated according to its assignment to the hippocampal head, body or tail (Fig. 2c,e; shown for the left hippocampus). Across lamellae, lamellar thickness increases progressively from childhood into early adulthood, reaching a peak in mid-adulthood, followed by an accelerated decline in later life. In contrast, lamellar width increases more rapidly during early development, stabilizes earlier, and subsequently undergoes a more gradual age-related reduction. This temporal dissociation is consistently observed across lamellae, indicating that distinct geometric features follow separable lifespan trajectories. Thickness captures a more prolonged pattern of maturation and degeneration, whereas width reflects earlier structural growth and earlier stabilization. Sex differences are evident in a subset of regions, with males generally showing higher values across several features. However, these effects are spatially heterogeneous across lamellae and anatomical subdivisions (Supplementary Figs. S4–S7).

### Heterogeneous lamella-based hippocampal geometric deviations across disorders

The lifespan normative reference enables quantification of individual-level deviations from age-resolved trajectories. Applying this framework across multiple brain disorders, we derived deviation scores by referencing individual measurements to the normative distribution. Representative profiles from a healthy control (HC) and an individual with dementia (DM) illustrate how lamella-wise measurements can be interpreted relative to population-level expectations (Fig. 3a).

**Fig. 3.**
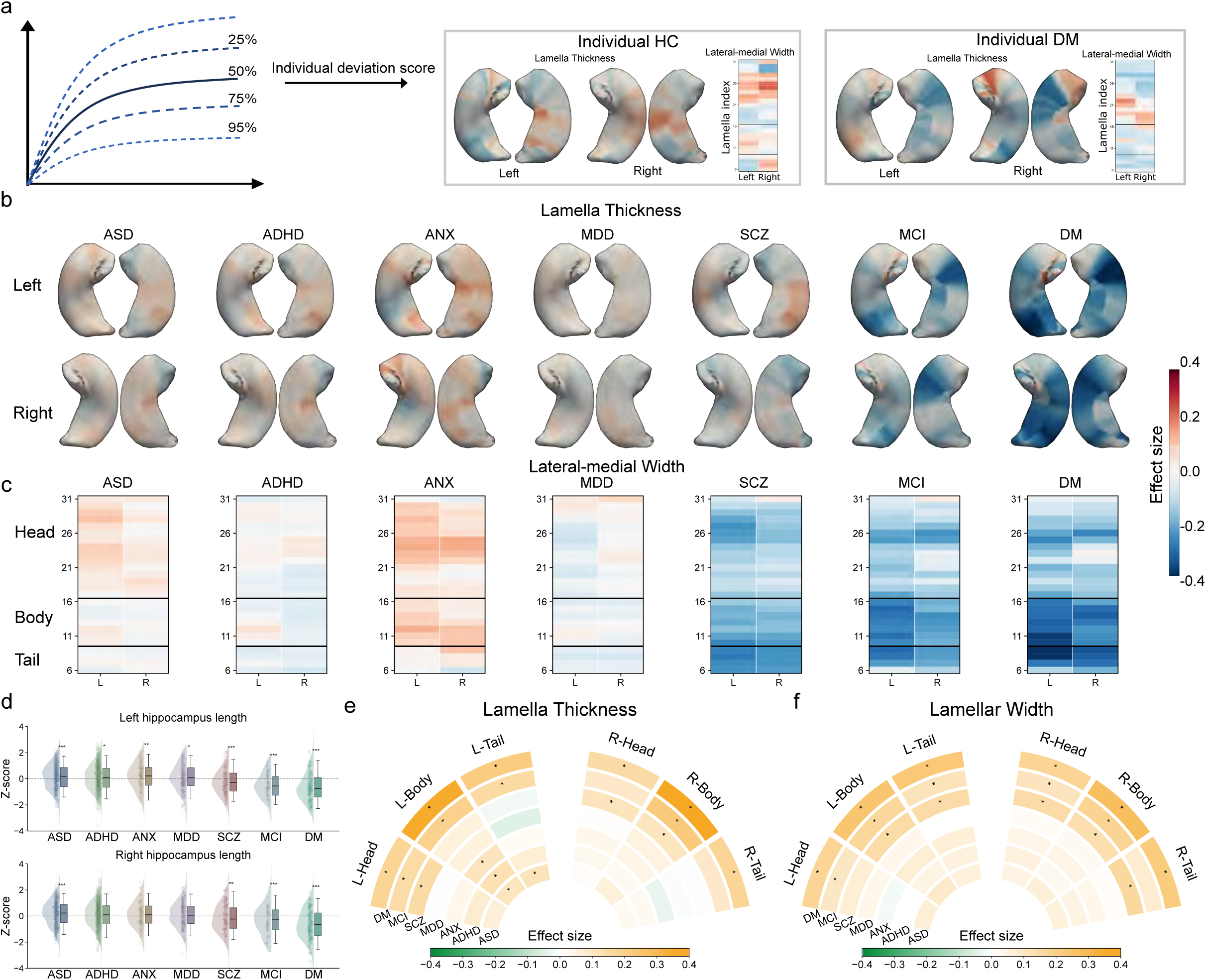
Hippocampal geometric deviations across multiple brain disorders reveal heterogeneous and spatially specific patterns. a, Schematic illustration of individual deviation scoring derived from the normative model. Individual feature values are referenced to the normative distribution and quantified as deviation scores, enabling subject-level assessment of hippocampal abnormality. Representative examples from a healthy control and an individual with dementia are shown. b, Spatial maps of effect sizes for lamellar thickness across multiple brain disorders, shown separately for the left and right hippocampus. c, Spatial maps of effect sizes for lateral-skeletal width across multiple brain disorders, shown separately for the left and right hippocampus. d, Group comparisons of hippocampal long-axis length in the left and right hemispheres across clinical cohorts. e,f, Region-level summaries of lamellar thickness and width effect sizes across hippocampal head, body and tail subdivisions in the left and right hemispheres. (* indicates corrected P < 0.05)

We next compared deviation maps from each disorder with those of healthy controls in the test set (Fig. 3b-d), revealing marked heterogeneity across conditions. Spatial patterns of lamellar thickness (Fig. 3b) indicated that effect sizes were modest in most disorders, with more pronounced alterations observed in SCZ, MCI and DM. Neurodevelopmental disorders, including ASD and ADHD, showed relatively weak effects (within ±0.05), characterized primarily by subtle increases in thickness in the bilateral hippocampal body. Notably, ASD, ADHD and ANX exhibited consistent focal reductions in the hippocampal head. MDD, by contrast, showed minimal deviation across regions. SCZ was associated with focal reductions in thickness, particularly in the hippocampal head, whereas both MCI and DM showed stronger and more spatially extensive deviations with highly similar anatomical patterns, indicative of shared trajectories of structural degeneration. Spatial patterns of lamellar width (Fig. 3c) revealed a broadly similar global trend, yet with distinct regional profiles across disorders. ADHD and MDD exhibited overall weak effects, whereas ANX showed a generalized widening. ASD showed a subtle increase, predominantly in the hippocampal head, whereas SCZ, MCI, and DM were characterized by widespread thinning, consistent with the atrophic pattern in thickness. Alterations in long-axis length (Fig. 3d) further differentiated these conditions. ASD and ANX tended to show relative elongation, with ASD exhibiting bilateral effects and ANX showing left-lateralized changes. In contrast, neurodegenerative conditions were marked by consistent shortening of the long axis. SCZ likewise demonstrated significant bilateral reductions in long-axis length, aligning with its overall pattern of structural contraction.

To summarize the overall burden of abnormality, we quantified composite deviation as the L1 norm of z scores (|z|) across the head, body and tail for both lamellar thickness and width (Fig. 3e,f). Colors denote effect sizes, with statistical significance indicated by * for FDR-corrected P < 0.05. Consistent with the spatial patterns described above, SCZ, MCI and DM exhibited larger effect sizes across both features, reflecting a greater burden of individual-level deviation. In SCZ, significant effects were primarily localized to lamellar thickness in the bilateral head and right body, as well as to width across most regions except the left head. By contrast, other disorders showed subtler and more spatially restricted effects: significant deviations were limited to the left body in ASD, ADHD and ANX, and to the left tail in ASD for thickness, with no significant effects observed for width. These indicate that hippocampal abnormalities are both spatially heterogeneous and disorder-specific.

### Lamella-based representation improves spatial sensitivity and clinical discrimination

To assess how spatial granularity influences sensitivity to disease-related alterations, we compared three partitioning schemes in patients with MCI and DM: the commonly used head-body-tail (HBT) tri-partition, the long-axis-based lateral-skeletal partition and the lamellar partition. Group-level spatial maps (Fig. 4a,b) showed that HBT-based features produced larger global effect sizes but substantially reduced spatial specificity. The localized alteration in the left hippocampal head in MCI was largely obscured under the HBT scheme, whereas our geometry-based representation precisely localized to the posterior portion of the head. In addition, this framework revealed a focal expansion within subregions of the hippocampal head, potentially reflecting localized inflammation-related changes that are blurred under coarser spatial definitions. Complementary geometric features further enhanced interpretability: both lamellar width and long-axis length provided additional sensitivity, with long-axis length showing significant reductions in MCI (left: effect size = −0.28) and more pronounced shortening in DM (left: effect size = −0.34).

**Fig. 4.**
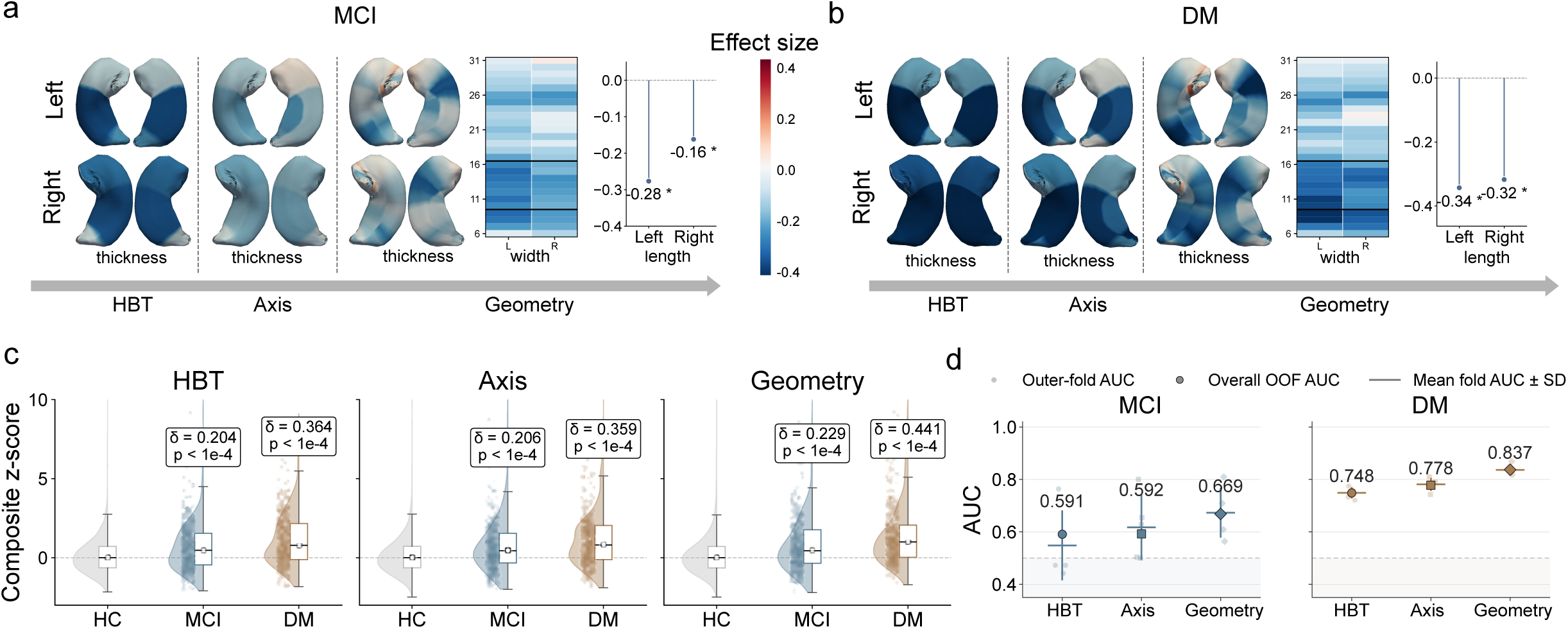
Fine-grained geometric representations improve sensitivity to disease-related hippocampus abnormalities and classification performance. a,b, Spatial patterns of hippocampus deviations in mild cognitive impairment (MCI) and dementia (DM) across three levels of representation, including head-body-tail (HBT), axis-based, and geometry-based models. Thickness, width and long-axis length are shown separately for the left and right hippocampus. Compared with coarser representations, the geometry-based framework revealed more spatially specific and stronger deviations. c, Group comparisons of composite deviation scores across healthy controls (HC), MCI and DM under HBT, axis-based and geometry-based representations. Cliff’s delta and corresponding P values are shown for pairwise comparisons with HC. d, Classification performance for MCI and DM using HBT, axis-based and geometry-based features. Points indicate outer-fold area under the receiver operating characteristic curve (AUC), squares indicate the overall out-of-fold AUC, and error bars indicate mean fold AUC ± s.d. Geometry-based features achieved the highest performance for both MCI and DM, supporting the advantage of fine-grained hippocampal geometry for detecting and classifying disease-related abnormalities.

At the deviation level, we quantified global abnormality using a calibrated composite deviation score. To account for differences in feature dimensionality, deviations were first aggregated using the L1 norm and then calibrated against the healthy control distribution (Fig. 4c). Coarse-grained partition based on HBT and axis features did not show a significant shift in MCI, whereas lamella-based features revealed clear deviations relative to controls (MCI: HBT, effect size = 0.204; axis, effect size = 0.206; lamella, effect size = 0.229; all P < 0.001). Increasing spatial granularity was associated with progressively larger group differences. A similar trend was observed in DM, where all representations reached statistical significance, with effect sizes increasing from HBT to lamellar partition (HBT: 0.364; axis: 0.359; ours: 0.441; all P < 0.001). We next evaluated discriminative performance using out-of-fold classification (Fig. 4e). To control for class imbalance and ensure comparability across feature sets, samples were balanced within each fold, and feature selection was performed within the training data under a stratified cross-validation framework, followed by linear support vector machine classification. Lamella-based features achieved the highest accuracy (AUC = 0.669 for MCI; 0.837 for DM), outperforming both HBT (0.591; 0.748) and axis-based representations (0.592; 0.778). These demonstrate that increasing spatial granularity enhances both sensitivity to disease-related deviations and discriminative performance, enabling more precise localization of hippocampal alterations and improving their utility for clinical characterization. (Additional results of comparison with volume in supplementary Fig. S10).

### Transferability and longitudinal characterization of hippocampal geometry

We next evaluated the transferability of the normative model to unseen data and its ability to capture longitudinal disease progression in an external cohort. The normative reference was directly applied to the Alzheimer’s Disease Neuroimaging Initiative (ADNI) without retraining, with site-specific intercepts re-estimated to account for acquisition heterogeneity (Fig. 5a). Calibration was performed using 50% of baseline cognitively normal (CN) individuals, and the remaining baseline and longitudinal data were used as the test set for downstream analyses. Representative individual trajectories (Fig. 5b) illustrated heterogeneous patterns of progression. In one case, deviations remained subtle during the MCI stage and increased sharply at conversion to AD (∼65 years), whereas in another case, elevated deviations were already evident during MCI (78.1 years), prior to clinical diagnosis. These observations indicate that structural deviations may precede or lag behind clinical diagnosis in a subject-specific manner. At the group level, longitudinal analyses using linear mixed-effects models revealed stage-dependent changes in lamellar thickness and width (Fig. 5c). Minimal changes were observed in CN–CN, followed by progressively increasing slopes with advancing disease. The CN–MCI stage already showed substantial changes, indicating early acceleration prior to clinical conversion, while the MCI–AD transition exhibited the strongest effects. In contrast, MCI–MCI and AD–AD stages were relatively stable in most substructures, consistent with a plateau following transition events. Long-axis length showed consistently small slope changes across all stages (|slope| < 0.05/year), suggesting more gradual longitudinal variation compared to thickness and width.

**Fig. 5.**
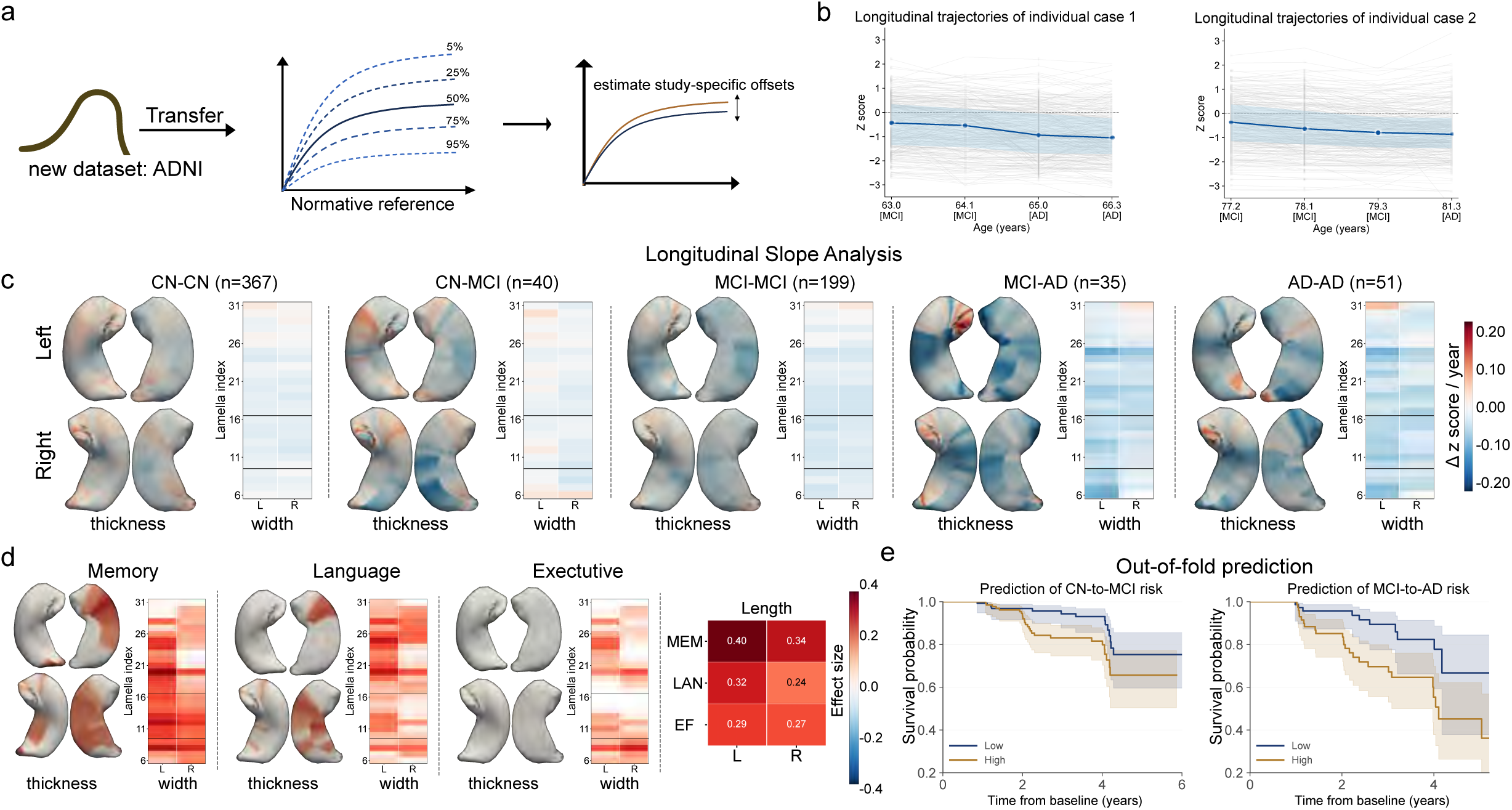
Normative geometric deviations support out-of-sample transfer and longitudinal characterization in an independent clinical cohort. a, Schematic illustration of transfer to the ADNI cohort. The normative reference established in the reference dataset was applied to the new ADNI dataset by estimating study-specific offsets, enabling deviation scoring in an independent longitudinal cohort without refitting the full normative model. b, Representative longitudinal trajectories of individual deviation scores from two participants in ADNI, illustrating subject-level tracking of hippocampal abnormalities across disease progression. c, Longitudinal slope analysis of lamellar thickness and lateral-skeletal width across diagnostic transition groups, shown separately for the left and right hippocampus. d, Associations between hippocampal deviation scores and cognitive performance in ADNI, shown for memory, language and executive function. e, Out-of-fold prediction of conversion risk for MCI and AD progression, shown by Kaplan-Meier survival curves stratified into low- and high-risk groups.

We next examined associations between hippocampal deviations and cognitive function using harmonized neuropsychological measures in ADNI (Fig. 5e). Memory showed the strongest associations, primarily involving inferior lateral of the bilateral hippocampus, with additional contributions from head and tail features. These effects were consistently observed for lamellar width. Language exhibited a similar but attenuated pattern, whereas executive function showed weaker associations overall. Long-axis length demonstrated robust associations with cognition, particularly memory (left: effect size = 0.40; right: 0.34). Finally, we evaluated the clinical utility of predicting disease progression and conversion risk (Fig. 5f). Out-of-fold predictions, combined with Kaplan–Meier and Cox proportional hazards revealed clear stratification between low- and high-risk groups. The geometry-based representation achieved significant discrimination for conversion from CN to MCI (p = 0.019) and from MCI to AD (p = 0.004). These results position hippocampal geometry as a transferable and promising clinically actionable marker that enables individual-level tracking of disease progression, resolves inter-individual heterogeneity beyond diagnostic categories, and provides a robust basis for risk stratification in neurodegenerative disease.

## Discussion

We establish a large-scale normative reference of hippocampal geometry across the human lifespan, leveraging data from over 27,000 individuals. By integrating long-axis length, lamellar thickness and width within a lamella-based representation, we systematically delineate hippocampal geometry in an anatomically consistent manner. This framework enables fine-grained characterization of spatial heterogeneity in hippocampal structure across ageing and disease. Across a range of neuropsychiatric and neurological conditions, hippocampal deviations exhibit distinct and spatially specific patterns that are robustly captured at the lamellar level. Importantly, this representation provides increased anatomical specificity and interpretability, enabling the detection of localized alterations that may be obscured by conventional coarse-grained biomarkers. Through transfer to an independent longitudinal cohort, we further demonstrate generalizability and support the utility of hippocampal geometry for individual-level tracking and disease progression. These results establish a scalable, generalizable framework for characterizing hippocampal structure, providing new insights into its spatial organization and alterations across the lifespan and in disease.

### Geometric mapping of hippocampal morphology

A key methodological advance of this study lies in the use of an intrinsic geometric framework that enables anatomically consistent comparison of hippocampal structure across individuals and over time. Unlike conventional surface-based morphological approaches^32,34^, which anchor correspondence to tissue interfaces that are often unstable under segmentation noise and scan variability, particularly in the hippocampal head, where inter-individual variability is large, our framework is rooted in aligning hippocampal lamella along the long-axis. This representation provides biologically interpretable measurements of long-axis length, lamellar thickness and width, enabling quantification of substructural variation across the lifespan. This distinction is critical, as fine-grained morphometric analysis requires anatomically meaningful correspondence across individuals to ensure interpretability and comparability^34,35^. This is particularly critical for lifespan studies, in which age-related trajectories must be resolved from highly heterogeneous anatomy, and for disease studies, in which subtle and spatially selective alterations may otherwise be obscured by coarse or unstable representations^35,36^. The resulting framework therefore does not merely increase spatial resolution, but enables a qualitatively different form of hippocampal analysis, one that is anatomically comparable, quantitatively stable, and suitable for large-scale lifespan and clinical investigation.

### Lifespan trajectories reveal spatially structured ageing

Our lifespan analysis reveals that hippocampal geometry follows a spatially structured and temporally dissociated pattern of maturation and ageing that is not captured by coarse summaries. Although subcortical grey matter volume is reported to peak around the second decade of life^1^, our results show that hippocampal geometric features do not follow a uniform developmental trajectory. Lamellar width peaks earliest, around the second decade, whereas thickness reaches its maximum predominantly after the third decade, and long-axis length continues to increase into early adulthood. This divergence indicates that hippocampal morphology cannot be adequately described by a single global trajectory, substantively extending the growing literature on differential subfield trajectories^37^ where volumetric measures conflate geometrically distinct properties that mature on distinct timescales.

Consistent with prior studies showing non-uniform age relationships across hippocampal subfields, anterior-posterior subregions and shape measure^38^, our results demonstrate that this heterogeneity extends beyond regions to include geometry. Different dimensions of hippocampal form therefore carry distinct temporal information, suggesting that geometric decomposition provides a more precise account of lifespan variation than region-averaged representations. This dissociation is further structured along the intrinsic longitudinal organization of the hippocampus. At the lamellar level, distinct territories exhibit asynchronous maturation and decline, indicating that hippocampal ageing reflects a coordinated but non-uniform process rather than a single global program^39^.

Importantly, the convergence of decline across geometric features in later life suggests that age-related atrophy arises from multiple, partially independent processes rather than a monotonic continuation of early developmental trajectories^40^. These findings support a view in which hippocampal morphology is best understood as a multi-dimensional system, in which distinct geometric components track separable aspects of structural organisation across the lifespan. This interpretation is consistent with evidence for graded molecular and functional organization along the hippocampal long axis^17^. The earlier maturation of lamellar width and the more protracted trajectory of thickness may reflect differences in the timing of large-scale structural organization versus local laminar refinement, linking macroscale geometric variation to underlying biological processes. This interpretation remains consistent with current models of hippocampal organization^41–43^.

### Disease-specific and heterogeneous deviation patterns

Building on the lifespan normative reference, our framework enables the quantification of hippocampal morphological deviations at both the individual and group levels with fine spatial resolution. At the individual level, this provides a sensitive means of tracking disease-related change. Representative trajectories (Fig. 5b) illustrate that the transition from CN to AD may be accompanied by increased feature-level variability, despite relatively preserved global medians. This suggests that early disease-related alterations may manifest as localized and heterogeneous deviations before converging into a more uniform pattern of atrophy^44–46^. In some cases, detectable geometric changes may precede clinical conversion, indicating that hippocampal morphology may capture preclinical progression not yet reflected in diagnostic criteria. These support the use of substructural markers for early detection and individualized monitoring of disease trajectories.

At the group level, we observe distinct but partially overlapping patterns of deviation across disorders. Neurodevelopmental and psychiatric conditions, including ASD, ADHD and ANX, show subtle and regionally specific increases relative to normative trajectories, most consistently reflected as increased lamellar thickness in the hippocampal body, with additional alterations in width and long-axis length observed in subsets of conditions, suggesting a shift in early structural development rather than uniform expansion. By comparison, neurodegenerative and severe psychiatric conditions, including SCZ, MCI and DM, exhibit a more consistent and spatially widespread pattern of reduction across thickness, width and length, indicative of structural atrophy. Notably, these patterns vary in their spatial distribution and magnitude across disorders, with SCZ showing more focal reductions in lamellar thickness, particularly in the anterior hippocampal head, together with more global reductions in width, whereas MCI, and DM exhibit more convergent shrinkage across geometric dimensions. This divergence in directionality indicates that hippocampal geometry differentiates between developmental deviation and degenerative loss across disorders.

The relative enlargement observed in neurodevelopmental and anxiety-related conditions may reflect deviations in early hippocampal development, consistent with prior evidence of increased hippocampal volume and atypical maturation in these disorders, suggesting a shift in developmental timing and structural refinement rather than uniform growth ^47–50^. By contrast, the reductions observed in schizophrenia and neurodegenerative conditions are consistent with progressive hippocampal atrophy, which has been linked to disruptions in memory and context processing^47,51–53^ and may contribute to impaired integration of internal representations with external inputs, particularly in anterior hippocampal regions, a process often discussed within self–world frameworks^9,54^. Hippocampal geometry thus provides a common reference for interpreting whether disease-related variation reflects altered development or progressive degeneration.

At the MCI stage, we observed a focal thickening in the left hippocampal head, particularly at its anterior tip. This pattern is unlikely to reflect a simple reversal of neurodegeneration but instead suggests that early hippocampal involvement is spatially heterogeneous and temporally non-linear. Prior studies across the Alzheimer’s disease continuum have reported hippocampal hyperactivation and regionally increased cortical thickness or grey-matter volume in early stages, which have been interpreted as compensatory remodelling, pathological hyperexcitability, or neuroinflammatory responses^55–57^. Consistent with these observations, our finding may reflect a transient and regionally selective process preceding overt atrophy. Several non-exclusive processes could underlie this effect, including compensatory plasticity, inflammation-related structural expansion, or the coexistence of local hypertrophy and degeneration due to subregional vulnerability^58–61^. Notably, the anterior hippocampal head, positioned at the interface of memory and affective circuits, may be particularly sensitive to early dysfunction^60,62,63^. Such localized deviations, confined to the anterior hippocampal head, would be largely obscured under coarser representations, underscoring the importance of spatially resolved geometric modelling for capturing early and heterogeneous disease processes.

### Strength and Limitations

A key strength of this study lies in the integration of lifespan normative modelling with fine-grained geometric characterization of the hippocampus, enabling the quantification of individual- and group-level deviations with high spatial specificity. By leveraging multi-level lamella-based representations, our framework captures subtle and spatially heterogeneous patterns that are not accessible to conventional volumetric or coarse parcellation approaches. This fine-grained design, combined with a large multi-site cohort and a distribution-aware normative modelling strategy, allows for robust estimation of deviation profiles and facilitates cross-dataset transfer, thereby providing a principled framework for studying disease-related heterogeneity and progression at the individual level. Besides, several limitations warrant consideration. First, the lamella-based stratification is critical for resolving fine-grained hippocampal geometry; however, not all lamellar subdivisions were equally informative in our analysis. To improve robustness, we regrouped features based on the finest reliable lamellar units and excluded unstable regions, particularly in the hippocampal tail (e.g., width indices 1– 5). While this strategy enhances statistical stability, it reduces the completeness of geometric coverage and may omit subtle but potentially relevant variations. Second, the age distribution of the cohort is uneven, with underrepresentation of infants and individuals in mid-adulthood. This limits our ability to fully capture lifespan trajectories, particularly early developmental dynamics. In particular, sparse infant data constrains the establishment of reliable normative references for early childhood and may obscure critical periods of rapid neurodevelopment. In addition, although multi-site data were incorporated, site-related variability remains a potential source of bias. Differences in scanner hardware, acquisition protocols and preprocessing pipelines can introduce systematic shifts in feature distributions that are not fully eliminated by statistical adjustment. In this work, we adopted a GAMLSS-based normative modelling framework, which models both location and higher-order distributional parameters and has been shown to better account for site effects and improve cross-cohort transfer compared with traditional harmonization approaches such as ComBat^1^. Nevertheless, residual site-dependent variability may still influence absolute deviation estimates and warrants further investigation. Future work should also extend beyond cross-sectional modelling to incorporate longitudinal data, enabling direct characterization of within-subject trajectories and reducing reliance on population-level inference of disease progression. Finally, the geometric measures used here (e.g., thickness, width and lamella-based indices) represent macroscopic structural proxies and do not map one-to-one onto underlying biological processes such as synaptic density, gliosis or neuroinflammation. Therefore, mechanistic interpretations should be made with caution, and future studies integrating multimodal imaging or histological data will be essential for linking these geometric signatures to their biological substrates.

### Conclusion

In this study, we establish a lamella-based geometric normative framework that captures the continuous internal organisation of the hippocampus and anchors individual variation to population-derived reference trajectories across the lifespan. Moving beyond conventional region-level summaries, this approach enables anatomically grounded, spatially resolved quantification of hippocampal morphology, yielding individualized deviation profiles that preserve fine-grained heterogeneity across development and disease. Hippocampal structure follows temporally dissociated and spatially organized trajectories, and disease-related alterations form distinct patterns that differentiate altered developmental processes from progressive degeneration. By enabling precise characterization of localized and heterogeneous deviations, including early and spatially confined alterations, this framework provides a principled basis for detecting subtle structural changes that may precede overt clinical manifestation. Generalization to independent longitudinal data further supports its robustness in large, multi-site settings and its utility for individual-level tracking and risk stratification. These findings highlight that hippocampal morphology is not adequately described by a single aggregate measure, but instead reflects a multi-dimensional geometric system with dissociable components that follow distinct lifespan trajectories. By explicitly modelling these geometric dimensions within an anatomically consistent framework, this work provides a principled basis for understanding how structural organisation, development and degeneration are differentially expressed across the hippocampus.

## Online Methods

### Ethics of the study

Description of informed consent and other ethical procedures is extensively provided in each referenced study, with additional information summarized in Supplementary Table S1. All data processing was conducted within the secure computing environment of the German Network for Bioinformatics Infrastructure (de.NBI), where data storage, and analysis comply with EU General Data Protection Regulation (GDPR) and relevant German data privacy laws was guaranteed. Further this infrastructure is ISO certified, which ensures security and the highest standards.

### Reference cohort

The cohort of individuals was aggregated from 158 scanning sites across multiple studies, including CAMCAN, SALD, SRPBS-OPEN, NKIRS, CORR, AIBL, ABIDE-I, ABIDE-II, SLIM, SRPBS-1600, GSP, ADHD200, COBRE, and FCON1000, HCP1200, HCP-Aging, HCP-Dev, IXI, OASIS, PING, PNC, PPMI, TCP and datasets from OpenNeuro (ds000030, ds000053, ds000119, ds000202, ds000208, ds000222, ds000228, ds000240, ds000243, ds001131, ds001408, ds001486, ds001734, ds001747, ds001748, ds001796, ds001838, ds001848, ds001894, ds002116, ds002236, ds002330, ds002345, ds002382, ds002385, ds002424, ds002643, ds002647, ds002717, ds002731, ds002785, ds002790, ds002837, ds002843, ds002886, ds003037, ds003097, ds003126, ds003138, ds003242, ds003346, ds003416, ds003436, ds003469, ds003481, ds003499, ds003508, ds003568, ds003592, ds003612, ds003643, ds003701, ds003709, ds003717, ds003745, ds003798, ds003826, ds003831, ds003877, ds003974, ds003988, ds004044, ds004146, ds004169, ds004173, ds004199, ds004215, ds004217, ds004261, ds004285, ds004302, ds004349, ds004466, ds004469, ds004512, ds004556, ds004589, ds004604, ds004636, ds004648, ds004697, ds004711, ds004718, ds004725, ds004746, ds004837). Further details on each study are provided in the corresponding publications (Supplementary Table 1). In total, the reference cohort included 20,907 healthy or non-diagnosed individuals (52.24% females). The participants’ ages ranged from 5 to 95 years (Fig. 1b). Comprehensive information on each scanning site, including sample size, mean age, standard deviation, and sex ratio, is available in Supplementary Table 2. Participants with missing demographic information, missing T1-weighted MRI data, or who had withdrawn from the respective studies were excluded from further analyses.

### Clinical cohort

As for the clinical datasets, we integrated data from ABIDE I, ABIDE II, ADHD200, AIBL, COBRE, HBN, OASIS3, SRPBS-1600, SRPBS-OPEN, TCP, ds002424, ds000030 and NKIRS. Clinical groups with available diagnostic information and more than 100 participants were included wherever possible. Based on these criteria, a total of 3,651 individuals with clinical conditions were included and grouped into seven diagnostic cohorts: autism spectrum disorder (ASD; n = 637), attention-deficit/hyperactivity disorder (ADHD; n = 1,610), generalized anxiety disorder (ANX; n = 339), major depressive disorder (MDD; n = 398), schizophrenia (SCZ; n = 251), mild cognitive impairment (MCI; n = 180), and dementia (DM; n = 183).

### Held-out data

In this study, we leveraged the ADNI cohort for external evaluation. Notably, in contrast to conventional evaluations that rely on splits between a reference cohort for training and clinically matched healthy individuals for testing, the held-out data in our setting were acquired from previously unseen scanner sites, thereby providing a stringent assessment of out-of-distribution generalization. However, the ADNI cohort comprises a large number of distributed acquisition sites, with longitudinal samples unevenly distributed across sites, resulting in limited per-site data that precludes robust site-specific transfer or recalibration. To address this, we instead adopted a study-specific transfer strategy, treating ADNI as a single external target domain while preserving the integrity of its longitudinal structure. Given the longitudinal design of ADNI, half of the baseline cognitively normal (CN) participants were used for model calibration, while the remaining baseline CN individuals formed the test set and were evaluated alongside MCI and AD participants from independent scans. In total, 211 CN participants were used for calibration, analogous to a training set, whereas the test set comprised 329 CN individuals (632 scans). The corresponding clinical cohorts included 276 MCI participants (582 scans) and 88 AD participants (164 scans).

### Preprocessing

For preprocessing, T1-weighted structural MRI scans were processed using FreeSurfer to obtain hippocampal segmentations and surface reconstructions. Automated quality control was performed based on the Euler number derived from the reconstructed surfaces, which serves as a proxy for topological defects and reconstruction reliability, and subjects not meeting the predefined criterion were excluded. The resulting hippocampal surfaces were then passed to the Axis-Referenced Morphometric Model (ARMM) pipeline, where cases that could not be successfully processed or mapped into a common anatomical representation were automatically removed through the regroup procedure. Lamellar thickness, lamellar width and long-axis length were subsequently derived within the ARMM framework, enabling spatially consistent quantification of hippocampal geometry across individuals.

### Geometry mapping

The axis-referenced morphometric model (ARMM) establishes a consistent interior coordinate system based on the longitudinal organization of the hippocampus. The model comprises two key components: an inscribed skeletal surface (IMS) lying in the superior-inferior hippocampal symmetry plane and a set of radial vectors (“spokes”) on the IMS.

ARMM employs a hippocampal atlas that captures the average anatomy derived from a large cohort of ex vivo 7T MRI and histological datasets^19^. This 3D probabilistic atlas, which encodes both the mean anatomy and the anatomic variability of hippocampal subfields, was constructed by applying groupwise diffeomorphic registration to high-resolution (0.2 mm isotropic) ex vivo MRI scans from 31 human hippocampal specimens.

Based on this template, manual landmarks were placed to adjust the position of the long-axis along the CA-DG border^32,64^, with the long axis fixed onto the IMS of the overall hippocampal shape. Through conformal mapping, this IMS was reparametrized to obtain layers-oriented perpendicular to the long-axis. The geometric constraints of skeletal representation were then applied to compute spokes on the skeletal surface, thereby constructing the lamellar architecture of the hippocampal volume.

The automated pipeline involves three main steps: (1) deformation of the ARMM template to individual subjects (cross-sectional); (2) refinement of IMS and spokes to fit the boundary surface of individual hippocampi (both cross-sectional and longitudinal); and (3) extraction of morphometric features (the local thickness, width and long-axis length).

First, the template ARMM was deformed to each target baseline hippocampus through a large deformation diffeomorphic metric mapping (LDDMM) framework that adjusted the velocity and momentum fields to match the target shape while preserving the skeletal axis constraints. For longitudinal data, hippocampal surface boundaries of all time points were rigidly aligned to the baseline. Second, spoke direction and length were adjusted following the skeletal geometry constraints^65,66^: (1) the implied surface of skeletal representation should agree with the original boundary surface; (2) spokes should be vertical to the boundary surface; (3) the spokes should not self-intersect, which ensures i) smooth interior and ii) a one-to-one correspondence between surface vertices and their associated thickness. The representation of longitudinal hippocampal morphology shares the same IMS position but refines its spokes to model biological atrophy or hypertrophy progression during ageing and disease. Finally, local thickness was defined as the Euclidean distance from each vertex on the superior or inferior surface to the IMS. Local width was computed along the parameterized lines on the IMS, which are perpendicular to the long axis. The long axis length is calculated as the length of the curved long axis.

The ARMM framework establishes good correspondence between spokes across individual hippocampus^67^. In this study, morphological measurements were defined at the lamellar level. Accordingly, local features (e.g., thickness and width) were regrouped into their respective lamellae, and the median value within each lamella was taken as the representative measurement. This resulted in 112 lamellar thickness features, 31 lamellar width features and a global long-axis length measure. Owing to instability in the tail region, five width features were excluded, yielding a final set of 26 lamellar width features (lamellae index 6–31).

### Normative modelling

Based on these anatomically referenced lamellar features, normative modelling was employed to characterize population-level variation and establish individualized reference distributions. To minimize potential bias arising from site- and demographic-related heterogeneity, the reference sample of individuals without clinical diagnoses was divided into training and test subsets using a stratified procedure based on scanning site, sex, and age.

Participants were first grouped by site so that each subset retained a representative contribution from every location, requiring at least 40 individuals per site. Within each site, the split was further balanced to preserve the distributions of sex and age. To facilitate fair comparison with the clinical cohorts, the test subset was additionally aligned to the patient groups with respect to age and sex, and, when feasible, site composition. This design allowed the healthy test sample to provide a suitable reference for evaluating disorder-related deviations. Participants from the diagnostic cohorts were introduced only at the testing stage, enabling an independent assessment of cross-disorder deviation profiles.

Normative modelling was performed within the Generalized Additive Models for Location, Scale and Shape framework using R version 4.1.2, called from Python through a custom rpy2-based interface. For each geometric feature, including lamellar thickness, width, and long-axis length, the model incorporated smooth effects of age and sex, while scanner site was modelled as a random intercept to account for site-specific variation. We used the SHASH (Sinh-Arcsinh) distribution family to accommodate non-Gaussian response distributions and improve sensitivity to distributional asymmetry and extreme centiles. By parameterizing not only the mean and variance, but also skewness and kurtosis as functions of covariates, this framework provides a flexible description of lifespan-related variation in hippocampal geometry. Specially, the z-score is obtained as:

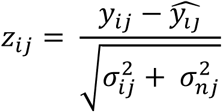

The computation of the z-score includes predicted mean *ŷ_ij_*, true response *y*_*ij*_, predicted variance *σ*_*ij*_ and normative variance *σ*_*nj*_. For model evaluation, the GAMLSS-based normative framework was applied to the held-out test set comprising only participants without clinical diagnoses, yielding estimates of the four distributional parameters, μ, σ, ν, and τ. Model performance was assessed using several complementary metrics, including explained variance, mean squared log-loss, skewness, and kurtosis. These measures characterized both overall goodness of fit and the extent to which the predicted distributions reproduced the empirical spread and higher-order properties of the observed data.

### Group comparisons

We compared z-scores between each clinical cohort and individuals of the test set using a nonparametric group-level analysis. Statistical significance was assessed with two-sided Mann–Whitney U tests, which are appropriate for comparing independent samples without assuming normality. To control for multiple comparisons, the resulting p-values were adjusted using false discovery rate correction. In addition to statistical significance, effect sizes were quantified using Cliff’s delta to capture the magnitude and direction of group differences.

### Slope analysis

To characterize stage-specific longitudinal changes in hippocampal geometry, we performed linear mixed-effects (LME) modelling using the ADNI cohort. Diagnostic stages were defined based on transitions between consecutive visits, including CN-CN, CN-MCI, MCI-MCI, MCI-AD and AD-AD. For each stage, non-baseline visits were paired with the corresponding subject-specific baseline measurement, and time was defined as years from baseline. For each geometric feature and hemisphere, LME models of the form

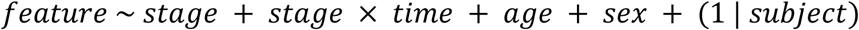

were fitted, where stage-specific slopes were derived from the interaction between stage and time. Stage-specific intercepts and slopes were estimated by including stage-specific indicator variables and their interaction with time. Age at baseline and sex were included as covariates. Subject-specific variability was accounted for by modelling random intercepts. The resulting stage-specific slopes represent the rate of change in feature z score per year from baseline, enabling direct comparison of progression rates across disease stages.

## Data availability

In this study, we used brain imaging data from multiple large-scale open-access neuroimaging initiatives, including CAMCAN, SALD, ADNI, SRPBS-OPEN, NKIRS, CORR, AIBL, ABIDE-I, ABIDE-II, SLIM, SRPBS-1600, GSP, ADHD200, COBRE, FCON1000, HCP1200, HCP-Aging, HCP-Development, IXI, OASIS, PING, PNC, PPMI, TCP, and several datasets from OpenNeuro. All datasets used in this study are publicly available and can be accessed upon reasonable request or through application procedures defined by the respective data repositories.

## Code availability

The code used in this study is publicly available GitHub at https://github.com/podismine/HippoLifespan. Actively maintained forks are hosted at https://github.com/MHM-lab, which also provide the GAMLSS-based normative modelling toolkit employed in this work. All trained models are distributed through these repositories and can be freely accessed for non-commercial use. The analyses were conducted using the following software packages: Deformetrica v4.3.0 (https://gitlab.com/icm-institute/aramislab/deformetrica), R v4.1.2 (https://www.r-project.org/), Python v3.10.12 (https://www.python.org/), pyvista v0.47.1 (https://pyvista.org/), vtk v9.6.0 (https://vtk.org/), GAMLSS v5.4.22 (https://www.gamlss.com/), scikit-learn v1.5.2 (https://scikit-learn.org), Nilearn v0.11.0 (https://nilearn.github.io/stable/index.html), and Matplotlib v3.9.2 (https://matplotlib.org/).

## Acknowledgements

We thank all participants and the clinicians and researchers involved in recruitment and assessment. YY and TW gratefully acknowledge support from the Alexander von Humboldt Foundation for YY’s research fellowship. We thank Stephen M. Pizer for insightful discussions and valuable guidance that contributed to the refinement of the overall framework of this study. TW acknowledges funding from the German Research Foundation (DFG; Emmy Noether Programme, grant 513851350), the BMBF/DLR project FEDORA (01EQ2403G) and funding from the Carl-Zeiss-Stiftung (P2022-00-087). TF received funding from the National Natural Science Foundation of China (82422027and U24A20340). This work was supported by the BMBF-funded de.NBI Cloud within the German Network for Bioinformatics Infrastructure (de.NBI; grants 031A532B, 031A533A, 031A533B, 031A534A, 031A535A, 031A537A–D, 031A538A), a secure processing environment for sensitive data.

Data used in the preparation of this article were obtained from the Alzheimer’s Disease Neuroimaging Initiative (ADNI) database (https://adni.loni.usc.edu). The ADNI was launched in 2003 as a public–private partnership, led by Principal Investigator Michael W. Weiner, MD. The primary goal of ADNI has been to test whether serial magnetic resonance imaging (MRI), positron emission tomography (PET), other biological markers, and clinical and neuropsychological assessments can be combined to measure the progression of mild cognitive impairment (MCI) and early Alzheimer’s disease (AD). For up-to-date information, see https://www.adni-info.org. As such, the investigators within ADNI contributed to the design and implementation of ADNI and/or provided data but did not participate in analysis or writing of this report. A complete listing of ADNI investigators can be found at: http://adni.loni.usc.edu/wp-content/uploads/how_to_apply/ADNI_Acknowledgement_List.pdf. Data collection and sharing for the Alzheimer’s Disease Neuroimaging Initiative (ADNI) is funded by the National Institute on Aging (National Institutes of Health Grant U19AG024904). The grantee organization is the Northern California Institute for Research and Education, and the study is coordinated by the Alzheimer’s Therapeutic Research Institute at the University of Southern California. ADNI data are disseminated by the Laboratory for Neuro Imaging at the University of Southern California. In the past, ADNI has also received funding from the National Institute of Biomedical Imaging and Bioengineering, the Canadian Institutes of Health Research, and private sector contributions through the Foundation for the National Institutes of Health (FNIH) (https://www.fnih.org), including generous contributions from AbbVie, Alzheimer’s Association, Alzheimer’s Drug Discovery Foundation, Araclon Biotech, BioClinica, Inc., Biogen, Bristol-Myers Squibb Company, CereSpir, Inc., Cogstate, Eisai Inc., Elan Pharmaceuticals, Inc., Eli Lilly and Company, EuroImmun, F. Hoffmann-La Roche Ltd and its affiliated company Genentech, Inc., Fujirebio, GE Healthcare, IXICO Ltd., Janssen Alzheimer Immunotherapy Research & Development, LLC., Johnson & Johnson Pharmaceutical Research & Development LLC., Lumosity, Lundbeck, Merck & Co., Inc., Meso Scale Diagnostics, LLC., NeuroRx Research, Neurotrack Technologies, Novartis Pharmaceuticals Corporation, Pfizer Inc., Piramal Imaging, Servier, Takeda Pharmaceutical Company, and Transition Therapeutics.

Data used in this study were provided in part by the Human Connectome Project (HCP), WU-Minn Consortium (Principal Investigators: David Van Essen and Kamil Ugurbil; 1U54MH091657), funded by the 16 NIH Institutes and Centers that support the NIH Blueprint for Neuroscience Research, and by the McDonnell Center for Systems Neuroscience at Washington University in St. Louis. Additional data were obtained from the Human Connectome Project in Aging (HCP-A; U01AG052564) and the Aging Adult Vulnerability and Resiliency in the Aging Adult Brain Connectome (AABC; U19AG073585), funded by the National Institute on Aging, as well as the HCP-Development 2.0 Release (DOI: 10.15154/1520708), supported by the National Institute of Mental Health (U01MH109589).

Data were also obtained from the Australian Imaging, Biomarkers and Lifestyle Study of Ageing, funded by the Commonwealth Scientific and Industrial Research Organisation (CSIRO), and made available via the ADNI database (http://www.loni.usc.edu/ADNI). AIBL investigators contributed data but did not participate in the analysis or writing of this report. A full list of investigators is available at http://www.aibl.csiro.au. Data used in the preparation of this article were obtained from the Parkinson’s Progression Markers Initiative database (https://www.ppmi-info.org). PPMI is a public–private partnership funded by the Michael J. Fox Foundation for Parkinson’s Research and its funding partners.

Data collection and sharing were also provided by the Cambridge Centre for Ageing and Neuroscience. CamCAN is funded by the UK Biotechnology and Biological Sciences Research Council (BB/H008217/1), with additional support from the UK Medical Research Council and the University of Cambridge.

Data were additionally obtained from the Pediatric Imaging, Neurocognition and Genetics (PING) study, now shared through the National Institute of Mental Health Data Archive (NDA). This publication reflects the views of the authors and does not necessarily represent those of the NIH or PING investigators.

## Author Contributions

Y.Y. and T.W. conceived the study and designed the overall analytical framework. N.G. implemented the ARMM framework and performed data preprocessing and feature analysis. Y.Y. conducted the large-scale modelling and statistical analyses. T.W. and T.G. guided the methodological development and supervised the project. S.D. and T.W. contributed to data curation and dataset development. N.S. contributed to the analytical development and supported the statistical analyses. All authors contributed to the interpretation of the results, critically revised the manuscript, and approved the final version for submission.

## Roles of funding

The funders had no influence on the study design, analyses or interpretations.

## References

1. Bethlehem, R. A. I. et al. Brain charts for the human lifespan. Nature 604, 525–533 (2022).

2. Rutherford, S. et al. Charting brain growth and aging at high spatial precision. eLife 11, e72904 (2022).

3. Rutherford, S. et al. The normative modeling framework for computational psychiatry. Nat. Protoc. 17, 1711–1734 (2022).

4. Wolfers, T. et al. Mapping the Heterogeneous Phenotype of Schizophrenia and Bipolar Disorder Using Normative Models. JAMA Psychiatry 75, 1146 (2018).

5. Marquand, A. F., Rezek, I., Buitelaar, J. & Beckmann, C. F. Understanding Heterogeneity in Clinical Cohorts Using Normative Models: Beyond Case-Control Studies. Biol. Psychiatry 80, 552–561 (2016).

6. Yang, X., Goh, A., Chen, S. A. & Qiu, A. Evolution of hippocampal shapes across the human lifespan. Hum. Brain Mapp. 34, 3075–3085 (2013).

7. Shafiei, G. et al. Reproducible Brain Charts: An open data resource for mapping brain development and its associations with mental health. Neuron 113, 3758–3779.e6 (2025).

8. Sun, L. et al. Human lifespan changes in the brain’s functional connectome. Nat. Neurosci. 28, 891–901 (2025).

9. Genon, S., Bernhardt, B. C., La Joie, R., Amunts, K. & Eickhoff, S. B. The many dimensions of human hippocampal organization and (dys)function. Trends Neurosci. 44, 977–989 (2021).

10. Borne, L. et al. Functional re-organization of hippocampal-cortical gradients during naturalistic memory processes. NeuroImage 271, 119996 (2023).

11. Ma, Y. et al. Age-Related Alterations in Hippocampal Microstructure Quantified Using High-Gradient Diffusion MRI (DMRI) in an Unfolded Hippocampal Space. Aging Cell 25, e70274 (2026).

12. Poppenk, J., Evensmoen, H. R., Moscovitch, M. & Nadel, L. Long-axis specialization of the human hippocampus. Trends Cogn. Sci. 17, 230–240 (2013).

13. Strange, B. A., Witter, M. P., Lein, E. S. & Moser, E. I. Functional organization of the hippocampal longitudinal axis. Nat. Rev. Neurosci. 15, 655–669 (2014).

14. Vos De Wael, R., et al. Anatomical and microstructural determinants of hippocampal subfield functional connectome embedding. Proc. Natl. Acad. Sci. 115, 10154–10159 (2018).

15. Xie, H., et al. Longitudinal hippocampal axis in large-scale cortical systems underlying development and episodic memory. Proc. Natl. Acad. Sci. 121, e2403015121 (2024).

16. Nordin, K., et al. Two long-axis dimensions of hippocampal-cortical integration support memory function across the adult lifespan. Preprint at 10.7554/eLife.97658.2 (2025).

17. Vogel, J. W. et al. A molecular gradient along the longitudinal axis of the human hippocampus informs large-scale behavioral systems. Nat. Commun. 11, 960 (2020).

18. Cembrowski, M. S. & Spruston, N. Heterogeneity within classical cell types is the rule: lessons from hippocampal pyramidal neurons. Nat. Rev. Neurosci. 20, 193–204 (2019).

19. Adler, D. H. et al. Characterizing the human hippocampus in aging and Alzheimer’s disease using a computational atlas derived from ex vivo MRI and histology. Proc. Natl. Acad. Sci. 115, 4252–4257 (2018).

20. Ayhan, F. et al. Resolving cellular and molecular diversity along the hippocampal anterior-to-posterior axis in humans. Neuron 109, 2091–2105.e6 (2021).

21. Andersen, P., Soleng, A. F. & Raastad, M. The hippocampal lamella hypothesis revisited11Published on the World Wide Web on 12 October 2000. Brain Res. 886, 165– 171 (2000).

22. Fraser, M. A. et al. Longitudinal trajectories of hippocampal volume in middle to older age community dwelling individuals. Neurobiol. Aging 97, 97–105 (2021).

23. Nordin, K. et al. Two long-axis dimensions of hippocampal-cortical integration support memory function across the adult lifespan. eLife 13, RP97658 (2025).

24. Homayouni, R. et al. Age-related differences in hippocampal subfield volumes across the human lifespan: A meta-analysis. Hippocampus 33, 1292–1315 (2023).

25. Rutherford, S. et al. The normative modeling framework for computational psychiatry. Nat. Protoc. 17, 1711–1734 (2022).

26. Haast, R. A. M. et al. Insights into hippocampal perfusion using high-resolution, multi-modal 7T MRI. Proc. Natl. Acad. Sci. 121, e2310044121 (2024).

27. Diers, K. et al. An automated, geometry-based method for hippocampal shape and thickness analysis. NeuroImage 276, 120182 (2023).

28. DeKraker, J. et al. Automated hippocampal unfolding for morphometry and subfield segmentation with HippUnfold. eLife 11, e77945 (2022).

29. Gao, N. et al. MR-based spatiotemporal anisotropic atrophy evaluation of hippocampus in ALZHEIMER’S DISEASE progression by multiscale skeletal representation. Hum. Brain Mapp. 44, 5180–5197 (2023).

30. Sloviter, R. S. & Lømo, T. Updating the Lamellar Hypothesis of Hippocampal Organization. Front. Neural Circuits 6, (2012).

31. Gao, N., et al. HippMetric: A skeletal-representation-based framework for cross-sectional and longitudinal hippocampal substructural morphometry. Preprint at 10.48550/ARXIV.2512.19214 (2025).

32. Gao, N., Ye, C., Chen, H., Hao, X. & Ma, T. MRI-based axis-referenced morphometric model corresponding to lamellar organization for assessing hippocampal atrophy in dementia. Hum. Brain Mapp. 45, e26715 (2024).

33. Devlin, J., Chang, M.-W., Lee, K. & Toutanova, K. BERT: Pre-training of Deep Bidirectional Transformers for Language Understanding. in Proceedings of the 2019 Conference of the North American Chapter of the Association for Computational Linguistics: Human Language Technologies, Volume 1 (Long and Short Papers) (eds. Burstein, J., Doran, C. & Solorio, T.) 4171–4186 (Association for Computational Linguistics, Minneapolis, Minnesota, 2019). doi:10.18653/v1/N19-1423.

34. DeKraker, J., Köhler, S. & Khan, A. R. Surface-based hippocampal subfield segmentation. Trends Neurosci. 44, 856–863 (2021).

35. DeKraker, J. et al. Evaluation of surface-based hippocampal registration using ground-truth subfield definitions. eLife 12, RP88404 (2023).

36. DeKraker, J. et al. HippoMaps: multiscale cartography of human hippocampal organization. Nat. Methods 22, 2211–2222 (2025).

37. Homayouni, R. et al. Two-year changes in hippocampal subfield volumes are age-dependent across the lifespan. Neurobiol. Aging 154, 62–70 (2025).

38. Tamnes, C. K., Bos, M. G. N., Van De Kamp, F. C., Peters, S. & Crone, E. A. Longitudinal development of hippocampal subregions from childhood to adulthood. Dev. Cogn. Neurosci. 30, 212–222 (2018).

39. Malykhin, N. V., Huang, Y., Hrybouski, S. & Olsen, F. Differential vulnerability of hippocampal subfields and anteroposterior hippocampal subregions in healthy cognitive aging. Neurobiol. Aging 59, 121–134 (2017).

40. Bussy, A. et al. Hippocampal shape across the healthy lifespan and its relationship with cognition. Neurobiol. Aging 106, 153–168 (2021).

41. Soltesz, I. & Losonczy, A. CA1 pyramidal cell diversity enabling parallel information processing in the hippocampus. Nat. Neurosci. 21, 484–493 (2018).

42. Mizuseki, K., Diba, K., Pastalkova, E. & Buzsáki, G. Hippocampal CA1 pyramidal cells form functionally distinct sublayers. Nat. Neurosci. 14, 1174–1181 (2011).

43. Chen, L. et al. Four-dimensional mapping of dynamic longitudinal brain subcortical development and early learning functions in infants. Nat. Commun. 14, 3727 (2023).

44. Planche, V. et al. Structural progression of Alzheimer’s disease over decades: the MRI staging scheme. Brain Commun. 4, fcac109 (2022).

45. Punzi, M. et al. Atrophy of hippocampal subfields and amygdala nuclei in subjects with mild cognitive impairment progressing to Alzheimer’s disease. Heliyon 10, e27429 (2024).

46. Wei, X. et al. Mapping cerebral atrophic trajectory from amnestic mild cognitive impairment to Alzheimer’s disease. Cereb. Cortex 33, 1310–1327 (2023).

47. Selemon, L. D. A role for synaptic plasticity in the adolescent development of executive function. Transl. Psychiatry 3, e238–e238 (2013).

48. Eltokhi, A., Janmaat, I. E., Genedi, M., Haarman, B. C. M. & Sommer, I. E. C. Dysregulation of synaptic pruning as a possible link between intestinal microbiota dysbiosis and neuropsychiatric disorders. J. Neurosci. Res. 98, 1335–1369 (2020).

49. Penzes, P., Cahill, M. E., Jones, K. A., VanLeeuwen, J.-E. & Woolfrey, K. M. Dendritic spine pathology in neuropsychiatric disorders. Nat. Neurosci. 14, 285–293 (2011).

50. Xing, Y., Mo, Y., Chen, Q. & Li, X. Synaptic pruning mechanisms and application of emerging imaging techniques in neurological disorders. Neural Regen. Res. 21, 1698–1714 (2026).

51. Moyer, C. E., Shelton, M. A. & Sweet, R. A. Dendritic spine alterations in schizophrenia. Neurosci. Lett. 601, 46–53 (2015).

52. Glantz, L. A. Reduction of Synaptophysin Immunoreactivity in the Prefrontal Cortex of Subjects With Schizophrenia: Regional and Diagnostic Specificity. Arch. Gen. Psychiatry 54, 943 (1997).

53. Saxena, S. & Caroni, P. Selective Neuronal Vulnerability in Neurodegenerative Diseases: from Stressor Thresholds to Degeneration. Neuron 71, 35–48 (2011).

54. Nour, M. M., Liu, Y., El-Gaby, M., McCutcheon, R. A. & Dolan, R. J. Cognitive maps and schizophrenia. Trends Cogn. Sci. 29, 184–200 (2025).

55. Ray, A., Loghinov, I., Ravindranath, V. & Barth, A. L. Early hippocampal hyperexcitability and synaptic reorganization in mouse models of amyloidosis. iScience 27, 110629 (2024).

56. Williams, M. E. et al. Higher cortical thickness/volume in Alzheimer’s-related regions: protective factor or risk factor? Neurobiol. Aging 129, 185–194 (2023).

57. Corriveau-Lecavalier, N., Adams, J. N., Fischer, L., Molloy, E. N. & Maass, A. Cerebral hyperactivation across the Alzheimer’s disease pathological cascade. Brain Commun. 6, fcae376 (2024).

58. Heneka, M. T. et al. Neuroinflammation in Alzheimer disease. Nat. Rev. Immunol. 25, 321–352 (2025).

59. Grady, C. L. et al. Evidence from Functional Neuroimaging of a Compensatory Prefrontal Network in Alzheimer’s Disease. J. Neurosci. 23, 986–993 (2003).

60. Phan, T. X. et al. Increased Cortical Thickness in Alzheimer’s Disease. Ann. Neurol. 95, 929–940 (2024).

61. Yakushev, I. et al. Increased hippocampal head diffusivity predicts impaired episodic memory performance in early Alzheimer’s disease. Neuropsychologia 48, 1447–1453 (2010).

62. Chen, J. et al. Selective vulnerability of hippocampal sub-regions in patients with subcortical vascular mild cognitive impairment. Brain Imaging Behav. 18, 922–929 (2024).

63. Angeli, P. A., DiNicola, L. M., Saadon-Grosman, N., Eldaief, M. C. & Buckner, R. L. Specialization of the human hippocampal long axis revisited. Proc. Natl. Acad. Sci. 122, e2422083122 (2025).

64. Gross, D. W., Misaghi, E., Steve, T. A., Wilman, A. H. & Beaulieu, C. Curved multiplanar reformatting provides improved visualization of hippocampal anatomy. Hippocampus 30, 156–161 (2020).

65. Damon, J. Smoothness and geometry of boundaries associated to skeletal structures, II: Geometry in the Blum case. Compos. Math. 140, 1657–1674 (2004).

66. Damon, J. Smoothness and geometry of boundaries associated to skeletal structures I: sufficient conditions for smoothness. Ann. Inst. Fourier 53, 1941–1985 (2003).

67. Pizer, S. M. et al. Skeletons, Object Shape, Statistics. Front. Comput. Sci. 4, 842637 (2022).

