## Supplementary Information for "Lamellar Normative Modelling of the Hippocampus Across the Human Lifespan"

### Common nomenclature

AD – Alzheimer's disease

ADHD - Attention deficit hyperactivity disorder

AIC - Akaike Information Criterion

ANX - Anxiety or phobic disorders

ASD - Autism spectrum disorder

BIC - Bayesian information criterion

CN - Control or cognitively normal

DX - Diagnosis

FDR - False discovery rate

GAMLSS - Generalised additive models for location scale and shape

GMV - Total cortical grey matter volume

HC – Healthy control

ICC - Intraclass correlation coefficient

IQR - Interquartile range

MCI – Mild cognitive impairment

MDD - Major depressive disorder

MRI - Magnetic resonance imaging

SCZ – Schizophrenia

### Supplementary Methods

#### 1. Statistics & Reproducibility

All datasets were preprocessed using Freesurfer (version 7.4.1) and deformetrica (4.3.0) within the de.NBI cloud platform to ensure identical preprocessing across cohorts. The preprocessing included hippocampus segmentation, registration, surface refinement, and spoke refinement. The preprocessing code is available here: https://github.com/freesurfer/freesurfer and https://gitlab.com/icm-institute/aramislab/deformetrica. When translating our models into external datasets or unseen clinical cohorts, particularly in a federated setting, this preprocessing pipeline should be followed. The reference cohort included 22,704 individuals (healthy participants across 157 scanning sites), split into training (20,746 individuals) and testing sets (1,958 individuals). No statistical method was used to predetermine sample size. Instead, all available participants meeting the quality criteria were included. All models and analyses were implemented in Python 3.11, using PyTorch 2.3.1, and executed on the de.NBI HPC platform with Singularity containers to ensure computational reproducibility.

#### 2. Axis-referenced morphometric modeling of hippocampal geometry

Our framework is implemented based on our previous developments in axis-referenced morphometric modeling, with specific adaptations and extensions tailored to the current large-scale analysis. Concretely, the implementation consists of the following components.

The axis-referenced morphometric model (ARMM) provides a geometrically consistent internal coordinate system for the hippocampus by explicitly modeling its longitudinal organization. The framework is based on two key components: an inscribed medial surface (IMS), defined within the superior-inferior symmetry plane of the hippocampus, and a set of radial vectors ("spokes") emanating from the IMS to the boundary surface.

The ARMM template is derived from a high-resolution hippocampal atlas constructed from ex vivo 7T MRI and histological data[1]. This probabilistic atlas captures both mean anatomy and inter-individual variability of hippocampal subfields and was generated using groupwise diffeomorphic registration of 31 ex-vivo specimens (0.2 mm isotropic resolution). Based on this atlas, anatomical landmarks were manually defined to constrain the long axis along the CA-dentate gyrus (DG) interface, ensuring anatomically meaningful alignment (following the procedures described in studies[2,3]). The long axis was subsequently embedded onto the IMS of the hippocampal shape.

To establish an intrinsic lamellar organization, the IMS was reparametrized using conformal mapping, yielding a set of coordinate lines approximately orthogonal to the long axis. Under medial representation constraints, spokes were then constructed at each point on the IMS, forming a lamellar decomposition of the hippocampal volume.

The automated ARMM pipeline consists of three main steps. First, the template model is deformed to individual hippocampal shapes using a large deformation diffeomorphic metric mapping (LDDMM) framework, which preserves topology and skeletal-axis consistency while matching subject-specific geometry. For longitudinal data, all follow-up surfaces are rigidly aligned to the baseline. Second, the IMS and spokes are iteratively refined to fit individual hippocampal boundaries, subject to geometric constraints: (i) the reconstructed boundary from the medial representation must agree with the observed surface; (ii) spokes are required to be approximately orthogonal to the boundary; and (iii) spokes must not intersect, ensuring a one-to-one correspondence between surface vertices and medial coordinates. In longitudinal settings, the IMS position is fixed across time points, while spoke geometry is updated to capture structural progression. Third, morphometric features are extracted from the resulting representation.

Specifically, local thickness is defined as the Euclidean distance from each surface vertex to the IMS. Local width is computed along parameterized trajectories on the IMS that are approximately perpendicular to the long axis. The global long-axis length is defined as the arc length of the curved longitudinal axis.

The ARMM framework establishes point-wise correspondence across individuals via the shared medial coordinate system. In this study, morphometric features were analyzed at the lamellar level. To improve robustness and interpretability, vertex-wise measurements were aggregated within each lamella, and the median value was used as the representative metric for that lamellar unit.

### Supplementary Results

#### 3. Evaluation of ARMM representation accuracy against other shape models

Five shape similarity metrics were employed for quantitative validation of the surface agreement between the reconstructed measurable surface of ARMM and the ground truth surface: average surface distance (Q), maximum surface distance (q), Hausdorff distance (HD), surface area error (Ae) and curvedness error (Ce). Let S represents the ground truth surface, T denotes the test surface, p and q are points on surfaces T and S respectively, and n is the number of points on the surface T. The average surface distance is defined as:

$$Q = \frac{1}{n}\sqrt{\sum_{i=1}^{n} \| p_i - q_i \|^2},\ i = 1, \dots, n.$$

The maximum surface distance is defined as:

$$q = \max \| p_i - q_i \|,\ i = 1, \dots, n$$

The Hausdorff distance is defined as:

$$HD(\vartheta, \xi) = \max\left(h(\vartheta, \xi), h(\xi, \vartheta)\right)$$

The curvedness at each vertex is defined by

$$c = \sqrt{(k_{max}^2 + k_{min}^2)/2},$$

where $k_{max}$ and $k_{min}$ represent the main curvature. These metrics provide an indirect assessment of measurement error in morphometric models, as greater reconstruction fidelity to the ground truth yields more accurate morphological measurements. In the distance measurements, $Q$ captures global deviation characteristics, $q$ identifies localized maximum point-wise errors, and HD quantifies topological consistency through bidirectional extreme value analysis.

To evaluate the measurement accuracy of ARMM on 3T MRI data, structural MRI scans from the ADNI dataset were used. In this dataset, each subject underwent three scans with one-year intervals between successive scans. The cohort comprised 271 participants, stratified into three groups: cognitively normal (CN), stable mild cognitive impairment (sMCI), and progressive mild cognitive impairment (pMCI). This included 87 CN individuals (40 males/47 females; age 75.06 ± 5.11 years), 102 sMCI patients (54 males/48 females; age 75.15 ± 7.22 years), and 82 pMCI patients (19 males/63 females; age 75.15 ± 7.22 years), yielding a total of 813 longitudinal scans.

Other morphometric representation methods included in the comparison were cm-rep, ds-rep, and SPHARM-PDM. After hippocampal segmentation and surface extraction, the four different morphometric models (ARMM, cm-rep, ds-rep, and SPHARM-PDM) were fitted to the hippocampal shape data, and the reconstructed boundary surfaces were compared with the ground truth surfaces, which were directly extracted from the binary label images of the hippocampus.

Supplementary Figure S1 shows the reconstructed surfaces of a randomly selected hippocampus using different shape models. Colors indicate the distance from each point on the reconstructed surface to the ground truth surface (top left), with red representing larger distances. All four models capture the global morphology well but differ in local details. m-s-rep and SPHARM-PDM are closest to the ground truth, both capturing the finger-like protrusions in the head region, with SPHARM-PDM providing better detail at the head curvature. cm-rep lacks local details in some regions, such as the medial and lateral head. ds-rep yields the most featureless surface due to reconstruction from fewer skeletal sample points. In terms of local accuracy, m-s-rep and SPHARM-PDM show evenly distributed errors below 1 mm, which is within image resolution. cm-rep exhibits larger errors at the medial head and medial tail, likely due to large curvature that ignores small undulations during deformation. ds-rep has few spoke vertices pointing to the boundary; the absence of red does not imply no error. The largest errors for ds-rep occur at the medial head and tail, similar to the error distribution of cm-rep.

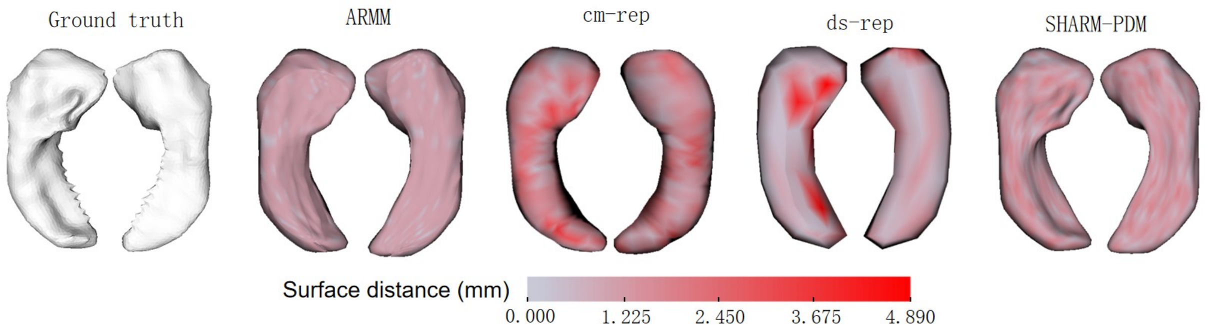

**Supplementary Fig. S1. The reconstructed boundary surface of a random selected hippocampus by different shape models with surface distance compared to the ground truth surface.**

The comparison results are listed in Table S3. At all observation points, the mean maximum surface distance of ARMM was 0.74 mm, which was smaller than that of the other methods: cm-rep (3.04mm), ds-rep (8.83mm), and SPHARM-PDM (1.05mm). In addition, reconstruction errors between the fitted and ground truth surfaces were computed separately for three time points: baseline (t0), the second observation (t1), and the third observation (t2).

Compared with the other methods, ARMM consistently produced the smallest reconstruction errors at all time points. The mean surface distance was approximately 0.003mm, and the mean maximum surface distance did not exceed 0.8mm, which is below the image resolution. The mean Hausdorff distance was substantially smaller than that of the other methods. The error in reconstructed surface area relative to the ground truth area was about 0.8mm$^2$ for ARMM. At t0, the curvature error of ARMM was comparable to that of SPHARM-PDM, both being lower than those of the other methods.

From t0 to t2, the mean reconstruction errors for ARMM, cm-rep, and SPHARM-PDM all decreased, whereas the point-wise surface distance error of ds-rep increased. A non-parametric Kruskal-Wallis test revealed no significant differences in reconstruction errors across the three time points for ARMM, indicating that the measurement accuracy of ARs-rep on hippocampal morphology was consistent across observation points. Furthermore, no significant difference in reconstruction accuracy was found between the two datasets, suggesting that the morphological representation accuracy of ARMM is consistent across different datasets.

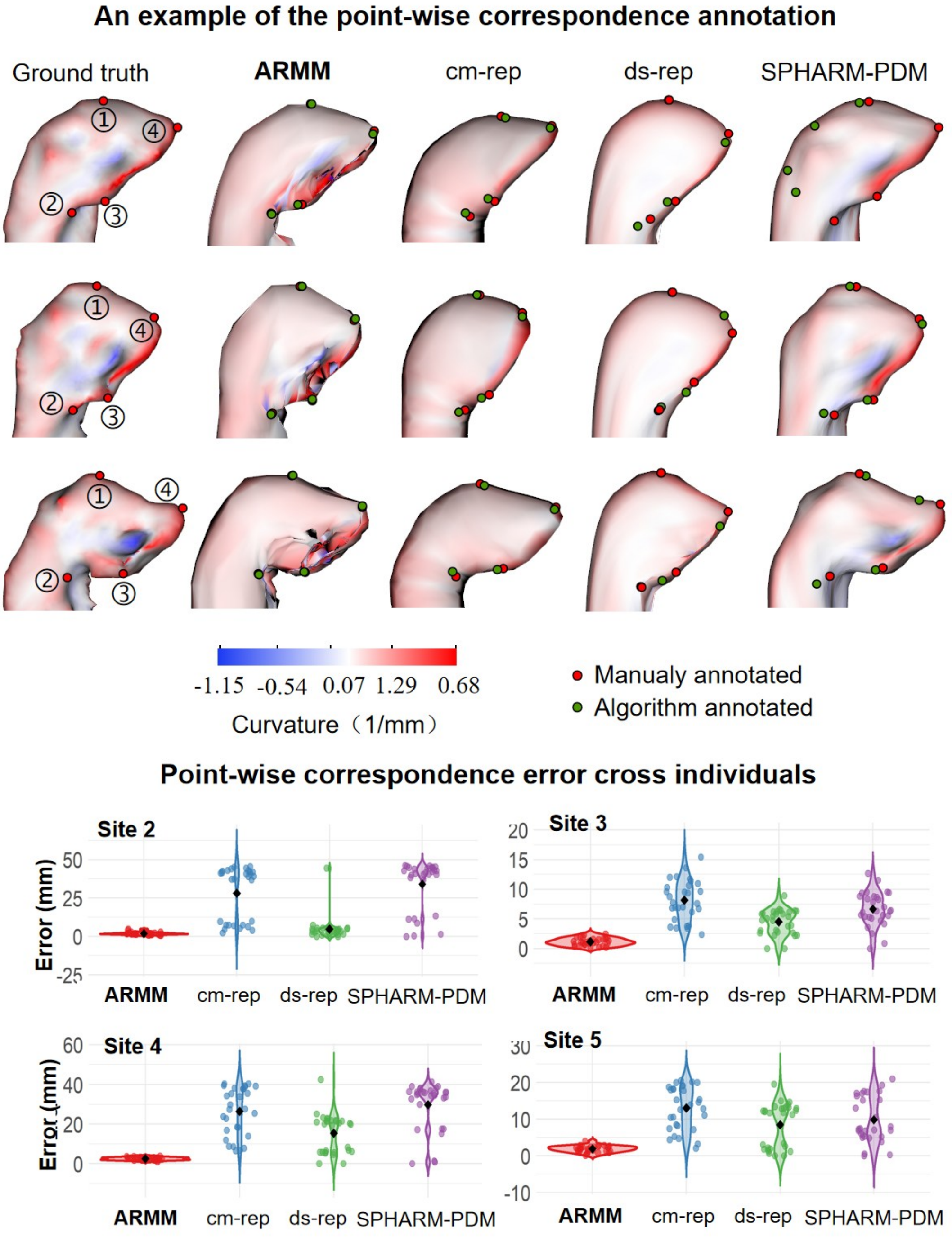

**Supplementary Fig. S2. Evaluation of the point-wise correspondence error across individuals.**

4. Evaluations of ARMM point-wise correspondence accuracy

The cross-subject and longitudinal correspondence accuracy is quantified by computing point-wise alignment errors between algorithm-derived correspondences and manually annotated landmarks on hippocampal surfaces reconstructed using different shape representation methods. Supplementary Figure S2 shows representative three hippocampal surfaces reconstructed by ARMM, SPHARM-PDM, cm-rep, and ds-rep, compared with ground-truth surfaces extracted from anatomical labels. While all methods produced anatomically plausible reconstructions, ARMM and SPHARM-PDM exhibited higher geometric fidelity to the original surfaces, as indicated by the similarity of curvature maps. In contrast, ds-rep showed substantial deviation from ground-truth due to its reduced surface resolution (low mesh vertexes).

We tested alignment accuracy based on data from ADNI. Because manual landmark is time-consuming and requiring high intra-rater consistency, a representative set of 30 subjects was randomly selected to cover the range of inter-individual shape variability. For each selected case, four anatomically identifiable landmarks were manually placed on both the ground-truth and reconstructed surfaces for error quantification. Across all methods and subjects, this resulted in 600 annotated points. The left panel of Supplementary Figure S2 illustrates the spatial layout of these landmarks, including the anterior and posterior extremities, medial folding apex, and lateral curvature center.

Alignment errors were quantified as Euclidean distances between algorithm-generated and manually annotated points. As summarized in the right panel of Supplementary Figure S2, ARMM achieved the lowest average correspondence errors across all landmarks, with a maximum mean error of 1.66 mm at the anterior folding apex. Errors at other locations remained below 1.1 mm on average, with standard deviations of less than 1 mm. In comparison, cm-rep and ds-rep showed moderate alignment accuracy, with maximum errors reaching 4.64 mm and 3.41 mm, respectively. SPHARM-PDM exhibited the largest variability, with a maximum error of 9.97 mm, likely due to pole flipping. Detailed correspondence errors for each method across all landmark locations are reported in Supplementary Table S4.

5. Evaluations of ARMM measurement reproducibility compared with hippounfold

Measurement reproducibility was quantified using the intraclass correlation coefficient (ICC), which was selected for its ability to evaluate both consistency and absolute agreement between repeated measurements. The ICC was calculated using a two-way mixed-effects model (ICC(3,1)), based on the formula:

$$\mathrm{ICC}(3,1) = \frac{\mathrm{MS_R} - \mathrm{MS_E}}{\mathrm{MS_R} + (\mathrm{k}-1)\mathrm{MS_E}},$$

where MSR is the mean square for rows (subjects), MSE is the residual mean square, and k is the number of measurements per subject. Note that the ICC values closer to 1 indicate stronger measurement stability.

To complement the reproducibility assessment and evaluate the information efficiency of extracted features, we performed principal component analysis to assess feature compactness. Compactness reflects how efficiently a feature set captures the underlying variance in the data, with higher compactness indicating that fewer features are needed to represent the same information. Comparable information density and stability among feature sets in different methods can confirm that the observed reproducibility reflects methodological reliability rather than incidental differences in shape representation.

In the experiment, we used longitudinal MRI data from cognitive normal subjects in ADNI, including 87 individuals with two scans acquired within one year; all morphological features were extracted at both time points for reliability analysis. The analysis evaluated both local thickness measurement stability (local ICC and variance ICC) and global reliability (Mean ICC and Global ICC). The numerical distribution and spatial patterns of ICC values are visualized in Supplementary Figure. S3, and detailed values are provided in Supplementary Tables S5 and S6. To benchmark substructural morphometric reproducibility, we compared ARMM against Hippunfold, currently the most advanced hippocampal thickness measurement method, using the same dataset.

Results demonstrated that the Inf_thickness and Sup_thickness (representing inferior and superior hippocampal thickness, respectively) showed ICC values around 0.6, comparable to those of Hippunfold. In contrast, width-related features, including Lat_width, Med_width, and Width, as well as Length, achieved mean ICC values above 0.91. Hippunfold only achieved values above 0.9 for hipp_surface and gyrification. Overall, ARMM exhibited smaller standard deviations in local ICC values than Hippunfold.

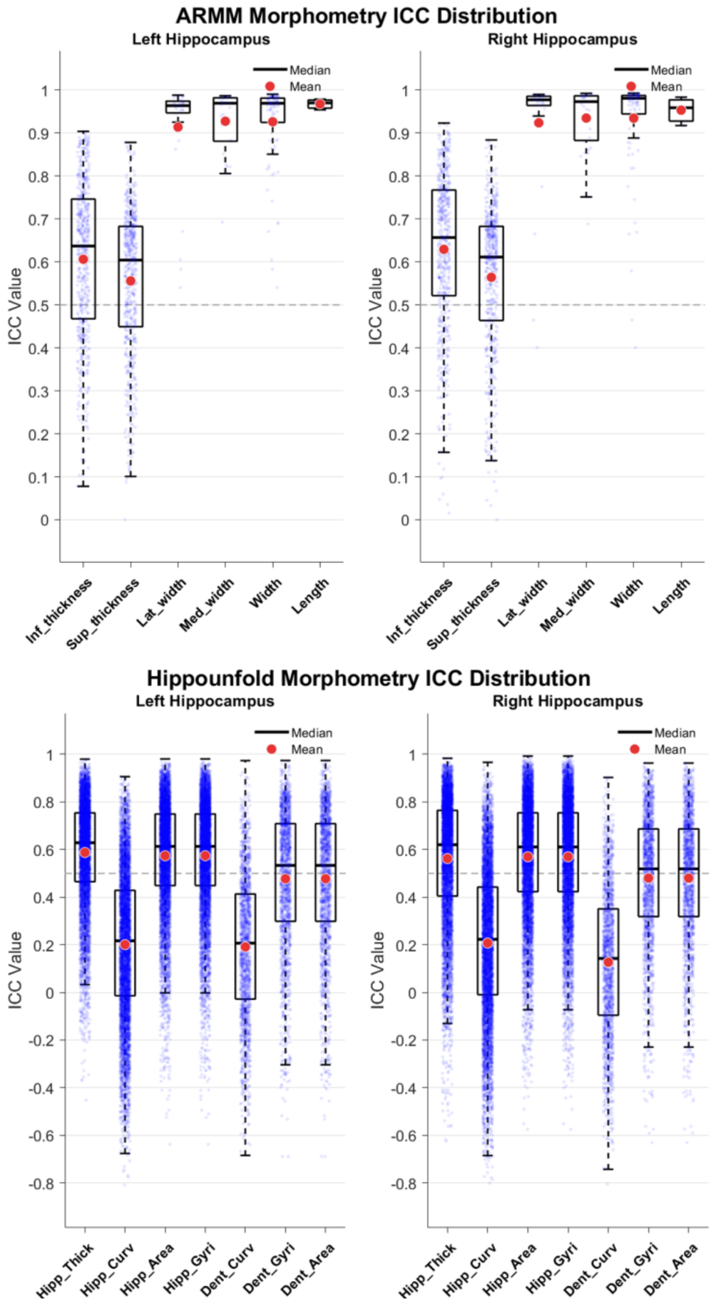

**Supplementary Fig. S3. ICC distributions of subfield measurements from ARMM and Hippunfold.**

#### 6. Evaluations of GAMLSS model specifications

We assessed the robustness of the normative modelling framework by systematically comparing several GAMLSS configurations that varied in how the scale component was parameterized and how age was modelled. In all configurations, the location parameter (μ) was defined consistently, using the same linear predictor formulation:

$$\eta_{\mu} = f_{sex}[age] + \beta_{sex}[sex] + b_{site}$$

where $f_{sex}(\cdot)$ denotes a sex-stratified smooth function of age. Sex was modelled as a fixed effect, while site was incorporated as a random intercept to account for inter-site variability. Age-related effects were captured using penalized B-splines. Multiple model configurations were evaluated; in configurations 1-6, the scale parameter (σ) was specified as:

$$\text{Config-1: } \eta_{\sigma} = \gamma_0 + \gamma_1[age], \eta_{\nu} = \delta_0, \eta_{\tau} = \kappa_0$$

$$\text{Config-2: } \eta_{\sigma} = \gamma_0, \eta_{\nu} = \delta_0, \eta_{\tau} = \kappa_0$$

$$\text{Config-3: } \eta_{\sigma} = \gamma_0 + \gamma_1[age] + \gamma_{sex}[sex], \eta_{\nu} = \delta_0, \eta_{\tau} = \kappa_0$$

$$\text{Config-4: } \eta_{\sigma} = \gamma_0 + \gamma_1[age], \eta_{\nu} = \delta_0 + \delta_1[age], \eta_{\tau} = \kappa_0 + \kappa_1[age]$$

$$\text{Config-5: } \eta_{\sigma} = \gamma_0 + \gamma_1[age], \eta_{\nu} = \delta_0, \eta_{\tau} = \kappa_0 + \kappa_1[age]$$

$$\text{Config-6: } \eta_{\sigma} = \gamma_0 + \gamma_1[age], \eta_{\nu} = \delta_0 + \delta_1[age], \eta_{\tau} = \kappa_0$$

Configurations 7–12 paralleled Configurations 1-6, respectively, with linear age terms replaced by their log-transformed counterparts in the corresponding model components. The results are summarized in Supplementary Table S7. Across all specifications, normative estimates remained highly consistent. The resulting Z-score distributions were centered close to zero with comparable variance across models, and higher-order distributional properties, including skewness and kurtosis, exhibited only minimal variation. To evaluate model fit and predictive performance, we randomly selected 100 features from both hemispheres as a representative subset for benchmarking. Model fit and predictive performance metrics, including logarithmic score, $R^2$, MAE, MSE, RMSE, AIC, and BIC, were highly comparable across all model specifications, further indicating that the normative modelling results were robust to alternative parameterizations of the scale component (σ) and to different treatments of age. Consistent with the distributional analyses, Z-score statistics remained well-calibrated across configurations, with means close to zero, standard deviations near unity, and minimal variation in skewness and kurtosis. Among the evaluated models, config-1 showed a slight but consistent advantage across multiple criteria, including lower information criteria (AIC/BIC) and competitive predictive performance, while maintaining well-calibrated Z-score distributions. Given its stable and overall favorable performance across a broad range of evaluation metrics, config-1 was selected as the final GAMLSS specification for all subsequent analyses.

#### 7. Normative trajectories of lamellar thickness across the lifespan

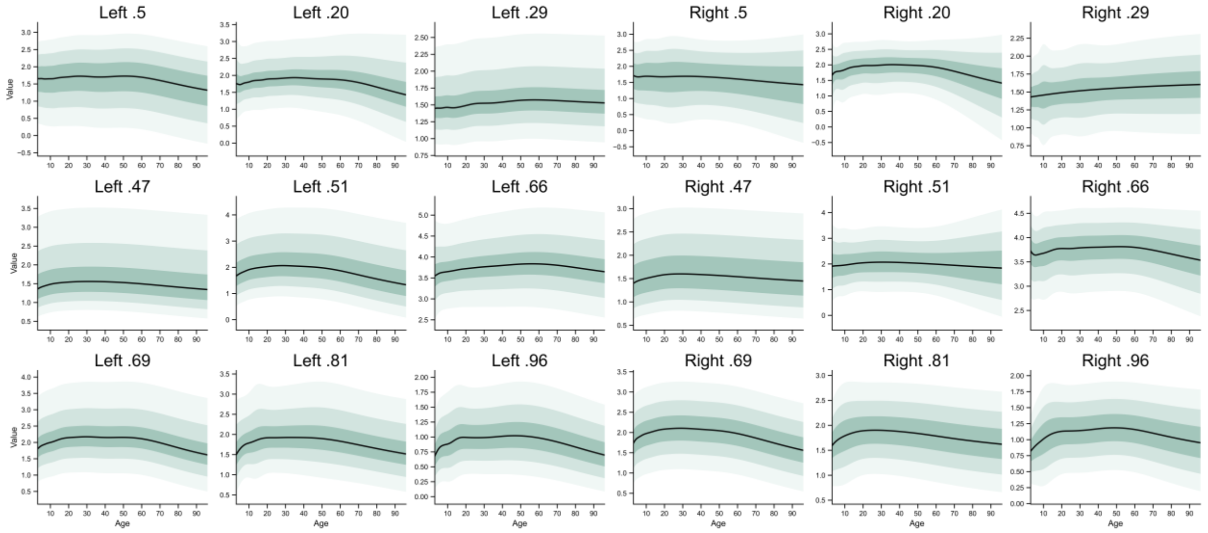

**Supplementary Fig. S4. Centile trajectories of lamellar thickness across the lifespan.** Centile curves for several lamellar thickness features randomly selected across the hippocampus, shown separately for the left and right hemispheres. Solid lines indicate the median (50th percentile), and shaded bands represent variability across the population (e.g., 5th–95th percentiles).

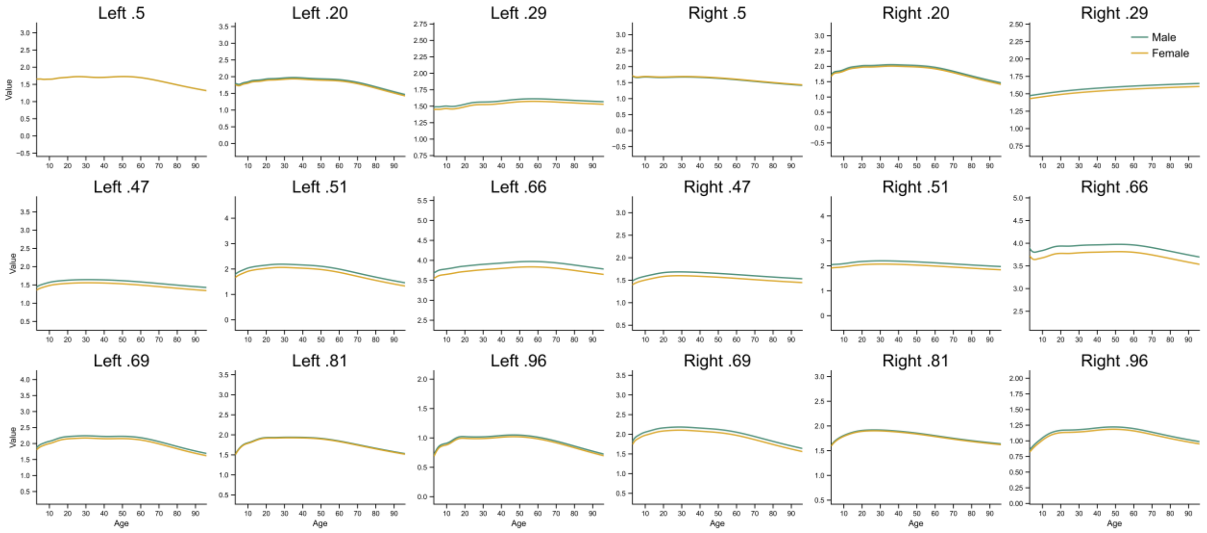

**Supplementary Fig. S5. Sex differences in lamellar thickness trajectories.** Sex-stratified lifespan trajectories of lamellar thickness for the same representative locations as in Supplementary Fig. S4. Male and female curves are overlaid to illustrate sex-specific effects.

To provide an intuitive illustration of the normative trajectories at a fine spatial scale, we randomly selected a subset of lamellar locations and visualized their centile curves and sex differences (Supplementary Figs. S4–S5). Overall, most lamellar thickness features exhibited a characteristic non-linear trajectory across the lifespan, with an initial increase followed by a gradual decline. In contrast, a subset of features showed an increase followed by a plateau, suggesting relative stability during later adulthood. This pattern indicates that certain lamellar locations may be less sensitive to developmental and ageing-related changes. Such reduced sensitivity could arise from the ultra-fine spatial granularity of the representation or may reflect inherently weak or heterogeneous local morphological variation. Comparisons between the left and right hippocampus revealed largely consistent global trends, supporting the overall symmetry of hippocampal organization. However, subtle lateralized differences were also observed. For example, at lamella index 96, the left hippocampus showed a decline after approximately 60 years of age, whereas the right side remained relatively stable, suggesting region-specific hemispheric divergence in ageing trajectories. Sex differences were present but not uniform across features. While some lamellar locations exhibited modest but consistent differences between males and females, others showed nearly overlapping trajectories, indicating that sex effects are spatially heterogeneous and not a dominant driver for all regions. Together, these results highlight both the shared global structure and the fine-grained variability of hippocampal morphology across space, sex, and the lifespan.

#### 8. Normative trajectories of lamellar width across the lifespan

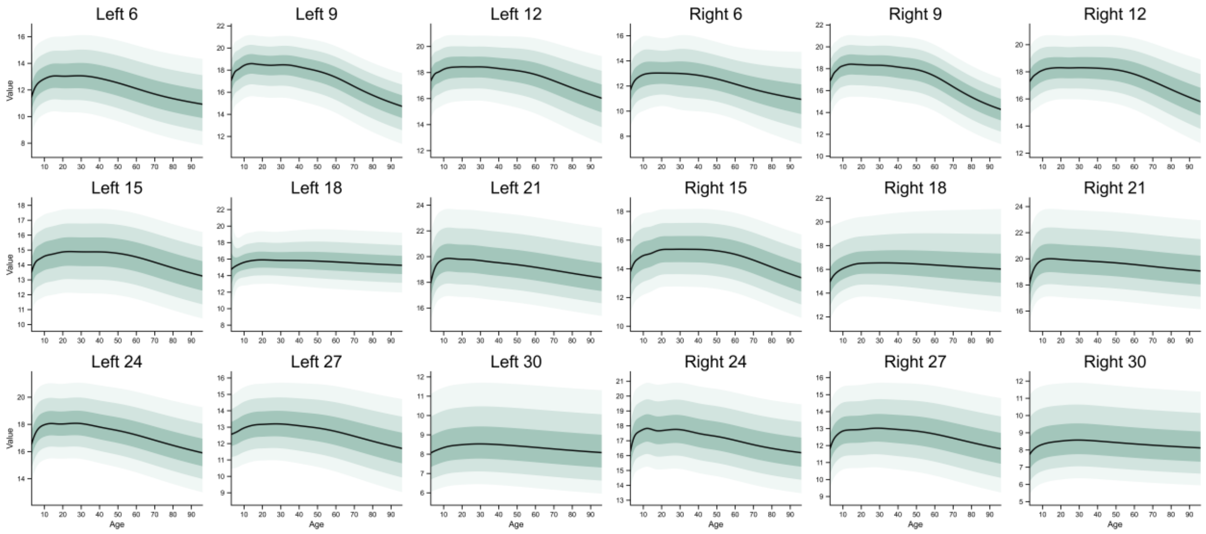

**Supplementary Fig. S6. Centile trajectories of lamellar width across the lifespan.** Centile curves for several lamellar width features randomly selected across the hippocampus, shown separately for the left and right hemispheres. Solid lines indicate the median (50th percentile), and shaded bands represent variability across the population (e.g., 5th–95th percentiles).

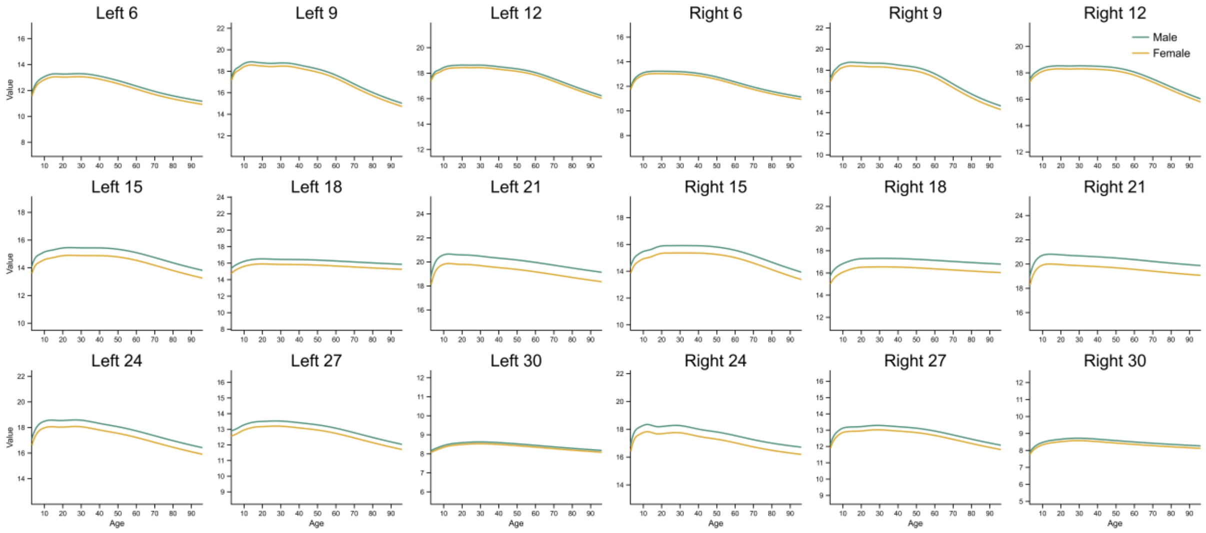

**Supplementary Fig. S7. Sex differences in lamellar width trajectories.** Sex-stratified lifespan trajectories of lamellar width for the same representative locations as in Supplementary Fig. S6. Male and female curves are overlaid to illustrate sex-specific effects.

Similarly, we visualized lamellar width at representative locations to examine its lifespan trajectories and sex differences (Supplementary Figs. S6–S7). Compared with lamellar thickness, lamellar width showed a stronger association with age, with more pronounced and consistent trajectories across the lifespan. Most features exhibited clear non-linear patterns, typically characterized by an early increase followed by a gradual decline, indicating that width may be more sensitive to developmental and ageing-related processes. In addition, sex differences were generally more evident for lamellar width than for thickness. Several regions displayed consistent offsets between males and females across a wide age range, suggesting that width captures more robust sex-related variation in hippocampal morphology. As with lamellar thickness, the overall trajectories were largely consistent between the left and right hippocampus, supporting a globally symmetric organizational pattern. Nevertheless, subtle regional discrepancies were observed at specific lamellar locations, indicating fine-grained hemispheric differences that are not apparent at coarser spatial scales.

#### 9. Deviation distributions comparison across groups

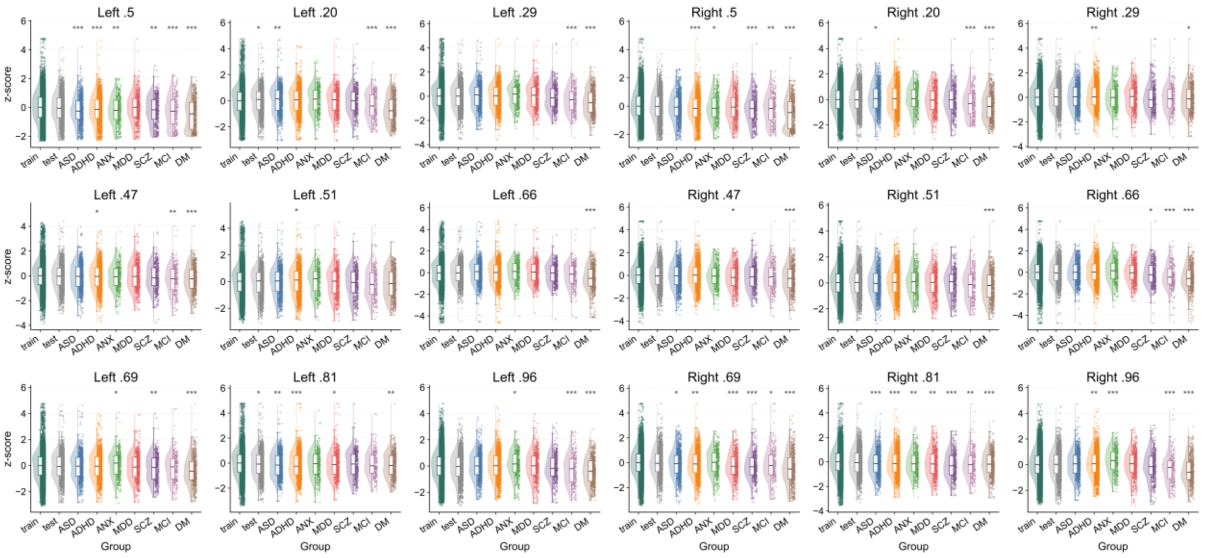

**Supplementary Fig. S8. Z-score distributions of lamellar thickness across groups.** Violin and box plots show the distribution of z-scores for representative lamellar thickness features in the training set, test set, and multiple clinical cohorts (ASD, ADHD, ANX, MDD, SCZ, MCI, DM), separately for the left and right hippocampus.

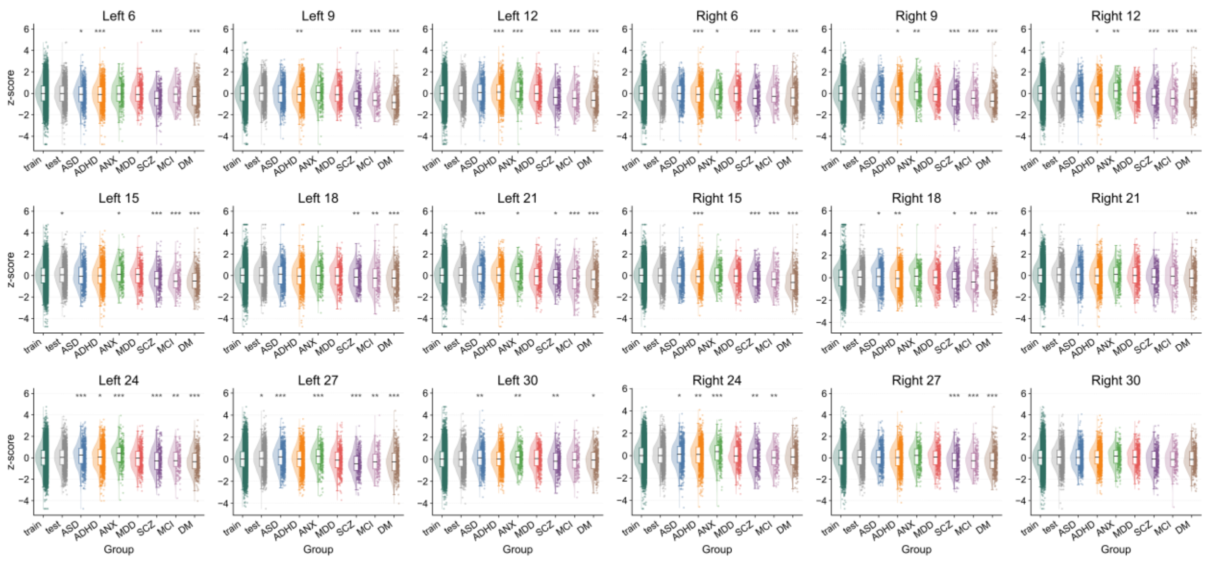

**Supplementary Fig. S9. Z-score distributions of lamellar width across groups.** Violin and box plots show the distribution of z-scores for representative lamellar width features across the same groups. Compared with thickness, width features exhibit clearer group-wise shifts in certain regions, while still maintaining consistent distributions between training and test sets.

We further examined the distribution of z-scores across training, test, and clinical cohorts for a representative subset of lamella-based features (Supplementary Fig. S8–S9). Overall, both the training and test sets exhibited approximately zero-centered Gaussian-like distributions, with comparable spread and no systematic shift between the two sets. Statistical comparisons revealed that most features did not show significant differences between training and test samples, indicating that the normative model generalizes well to unseen data and does not exhibit evident overfitting. In contrast, clinical cohorts showed varying degrees of deviation from the normative reference, with significance observed in a subset of features depending on the disorder. Notably, these alterations were not uniformly distributed across features or diagnostic groups, suggesting substantial heterogeneity in hippocampal geometry across disorders. The absence of consistent, shared patterns across all conditions further indicates that disease-related effects are spatially specific rather than globally uniform.

#### 10. Classification comparison across different types of features

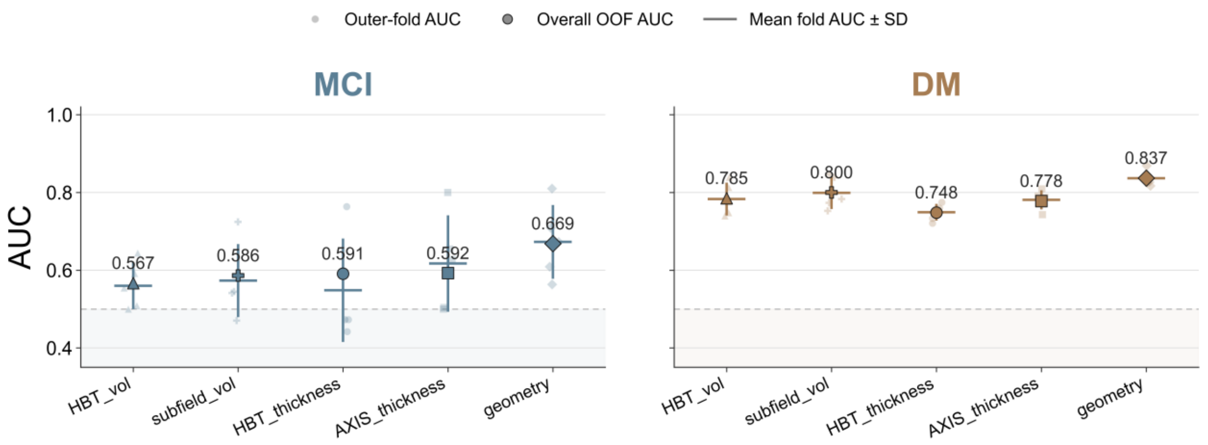

**Supplementary Fig. S10. Comparison of classification performance across feature representations.** Comparison of classification performance (AUC) for MCI (left) and dementia (DM, right) using different hippocampal feature representations, including HBT volume, subfield volume, HBT-level thickness, axis-based thickness, and the proposed geometry representation. Points indicate outer-fold AUCs from cross-validation, with larger markers denoting overall out-of-fold (OOF) AUC.

To further contextualize the performance of the proposed geometry-based representation, we additionally compared it with conventional volume-derived features, including hippocampal head–body–tail (HBT) volumes and subfield volumes obtained from FreeSurfer segmentation. The hippocampal subfield volumes were defined based on FreeSurfer segmentation, including CA1 (head and body), CA3 (head and body), CA4 (head and body), dentate gyrus (GC-ML-DG; head and body), subiculum (head), presubiculum (head and body), parasubiculum, molecular layer (head and body), hippocampal fissure, fimbria, and the hippocampus–amygdala transition area (HATA). Across both classification tasks, geometry-based features consistently achieved the highest discriminative performance. For MCI classification, volume-based features showed relatively limited sensitivity, with HBT volume and subfield volume yielding AUCs of 0.567 and 0.586, respectively. Thickness-based representations (HBT and axis-based) provided modest improvements (AUC $\approx$ 0.59), whereas the geometry representation achieved a notably higher AUC of 0.669. A similar pattern was observed for dementia (DM), where volume features already demonstrated relatively strong performance (HBT: 0.785; subfield: 0.800), consistent with the well-known sensitivity of volumetric atrophy to advanced neurodegeneration. However, the geometry representation still outperformed all alternatives, achieving the highest AUC (0.837), surpassing both volume-based and thickness-based features. These results suggest that although volumetric measures are sensitive to disease-related changes, particularly in later stages, they remain limited in capturing fine-grained and spatially heterogeneous morphological alterations. In contrast, the geometry-based representation integrates lamella-resolved thickness, width, and long-axis organization, enabling a more comprehensive characterization of hippocampal structural variation. This likely underlies its superior performance, especially in earlier disease stages such as MCI, where subtle and localized alterations are not fully reflected by bulk volume measures.

#### 11. Sensitivity Analysis of cross-cohort transfer sample size

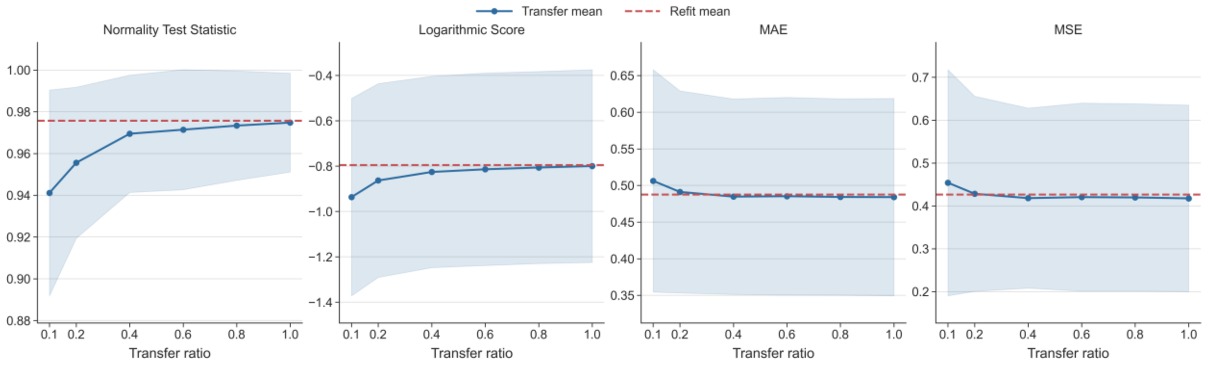

**Supplementary Fig. S11. Sample-size effects on cross-cohort transfer stability of the GAMLSS-based normative model.** Transfer stability was quantified using normality test score, mean absolute error (MAE), mean squared error (MSE), and log-likelihood (Logarithmic Score).

To evaluate the sample-size requirements for cross-cohort transfer in a real-world clinical setting, we performed a calibration analysis using the ADNI cohort. A total of 211 cognitively normal (CN) participants were used as the calibration pool, from which varying proportions were sampled to estimate study-specific location offsets (μ) while keeping all other distributional parameters fixed. For each sampling ratio, calibration subsets were repeatedly drawn and the transfer procedure was conducted to assess robustness across runs. Model performance in the held-out data was quantified using the Normality Test Statistic, Logarithmic Score, mean absolute error (MAE), and mean squared error (MSE).

As shown in Supplementary Fig. S11, all evaluation metrics exhibited systematic improvement with increasing calibration sample size. The Normality Test Statistic progressively increased and approached the refit baseline, indicating improved alignment between transferred distributions and empirical data. Similarly, the Logarithmic Score showed a monotonic increase, reflecting better likelihood fit under larger calibration subsets. In parallel, both MAE and MSE decreased with increasing sample size, with the most substantial gains observed at lower sampling ratios. Notably, all metrics demonstrated rapid improvement at small sample sizes, followed by gradual stabilization beyond approximately 40–60% of the calibration pool, suggesting diminishing returns with further increases.

Importantly, even with relatively small calibration subsets, the transferred model achieved performance close to the refit baseline, indicating strong robustness of the normative framework under limited calibration data. Overall, these results demonstrate that stable and reliable cross-cohort transfer can be achieved with a moderate number of calibration samples in heterogeneous clinical datasets such as ADNI, supporting the practical applicability of the framework in real-world scenarios where large calibration cohorts may not be available.

### 12. Cross-cohort transferability of normative models

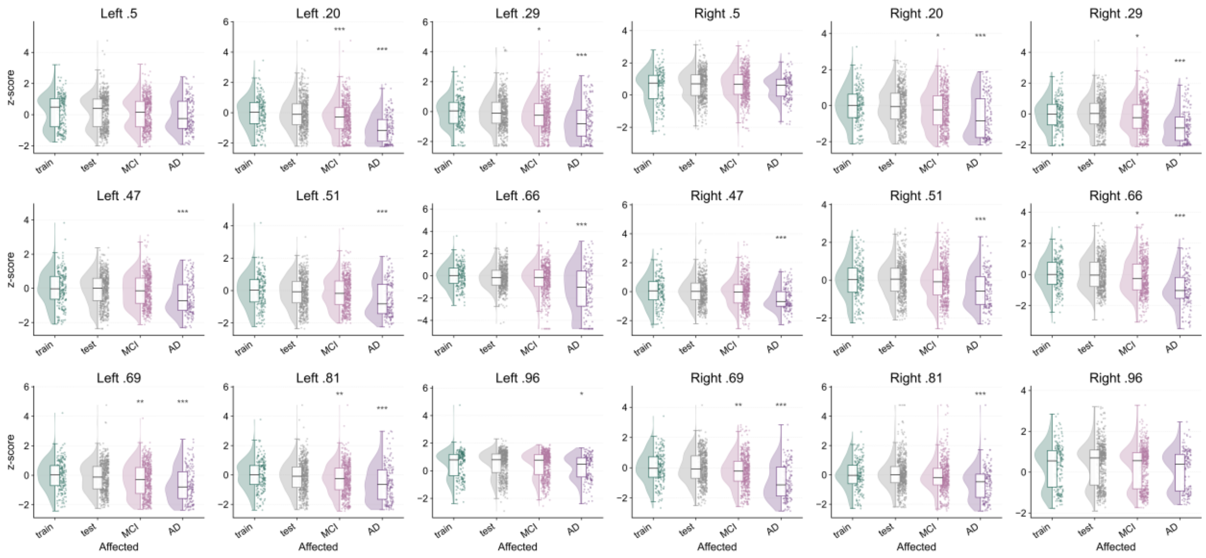

**Supplementary Fig. S12. Z-score distributions after transfer to the ADNI cohort on lamellar thickness across training data, test data and clinical groups (MCI and AD).**

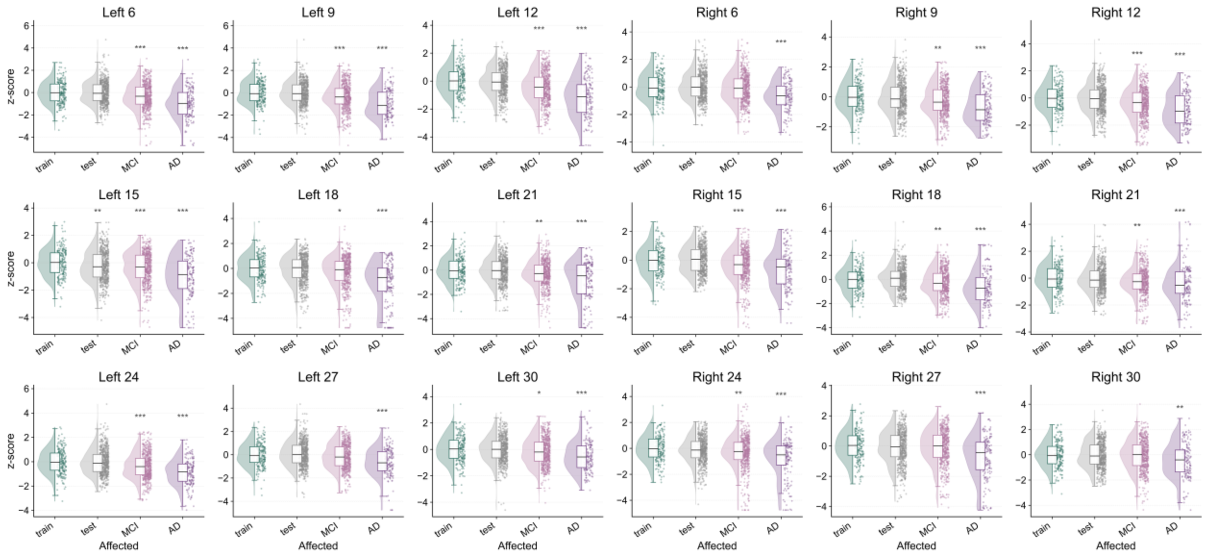

**Supplementary Fig. S13. Z-score distributions after transfer to the ADNI cohort on lamellar width across training data, test data and clinical groups (MCI and AD).**

We examined the distribution of z-scores across training data, test set, and clinical groups in ADNI. The test set comprised cognitively normal (CN) individuals from both partial baseline and longitudinal follow-up visits. Across both lamellar thickness and lateral-medial width features, z-score distributions in the training and test CN groups were highly comparable, with both centered around zero and exhibiting similar dispersion patterns. No systematic shifts or significant differences were observed between these two groups, indicating that the transferred model preserves calibration and maintains consistent normalization performance in unseen data, even under longitudinal sampling. In contrast, clinical groups exhibited clear deviations from the normative distribution. Individuals with mild cognitive impairment (MCI) showed moderate shifts in z-score distributions relative to controls, while AD cases demonstrated more pronounced and widespread deviations. These effects were consistently observed across multiple lamellar locations and in both hemispheres, supporting the sensitivity of the geometry-based representation to disease-related alterations. Notably, while thickness features showed relatively subtle but spatially heterogeneous deviations, width features exhibited stronger and more consistent shifts across disease stages, aligning with their higher sensitivity to age- and disease-related structural variation observed in the main analyses. Overall, these results confirm that the normative framework, after transfer, achieves robust cross-cohort generalization while retaining the ability to capture clinically meaningful deviations.

### Supplementary Tables

#### 13. Supplementary Table S1. Sources of all the studies used in the study

| Datasets | Sources | Comments | References |
|---|---|---|---|
| Autism Brain Imaging Dataset Exchange (ABIDE-1) | http://fcon_1000.projects.nitrc.org/ | Primary support for the work by Adriana Di Martino was provided by the NIMH (K23MH087770) and the Leon Levy Foundation. Primary support for the work by Michael P. Milham and the INDI team was provided by gifts from Joseph P. Healy and the Stavros Niarchos Foundation to the Child Mind Institute, as well as by an NIMH award to MPM (R03MH096321). | [4] |
| Autism Brain Imaging Dataset Exchange II (ABIDE-2) | http://fcon_1000.projects.nitrc.org/ | Primary support for the work by Adriana Di Martino and her team was provided by the National Institute of Mental Health (NIMH 5R21MH107045). Primary support for the work by Michael P. Milham and his team provided by the National Institute of Mental Health (NIMH 5R21MH107045); Nathan S. Kline Institute of Psychiatric Research). Additional Support was provided by gifts from Joseph P. Healey, Phyllis Green and Randolph Cowen to the Child Mind Institute. | [4] |
| ADHD-200 | https://fcon_1000.projects.nitrc.org/indi/adhd200/ | Data used in the preparation of this article were obtained from the ADHD-200 Sample (http://fcon_1000.projects.nitrc.org/indi/adhd200/). The ADHD-200 Sample was contributed by the ADHD-200 Consortium. | [5] |
| Cam-CAN | https://camcan-archive.mrc-cbu.cam.ac.uk/dataaccess/ | Data collection and sharing for this project was provided by the Cambridge Centre for Ageing and Neuroscience (CamCAN). CamCAN funding was provided by the UK Biotechnology and Biological Sciences Research Council (grant number BB/H008217/1), together with support from the UK Medical Research Council and University of Cambridge, UK. | [6,7] |
| Consortium for Reliability and Reproducibility (CORR) | http://fcon_1000.projects.nitrc.org/ | The National Institute on Drug Abuse (NIDA) and the National Natural Science Foundation of China (NSFC) have been instrumental in the CoRR collaboration providing the necessary funding and manpower to build the foundation of the project along with the Child Mind Institute, the Institute of Psychology, Chinese Academy of Sciences and the Nathan Kline Institute. | [8] |
| The Center for Biomedical Research Excellence (COBRE) | https://fcon_1000.projects.nitrc.org/indi/retro/cobre.html | The imaging data and phenotypic information was collected and shared by the Mind Research Network and the University of New Mexico funded by a National Institute of Health Center of Biomedical Research Excellence (COBRE) grant 1P20RR021938-01A2 | [9,10] |

| | | | |
|---|---|---|---|
| FCON1000 | https://fcon_1000.projects.nitrc.org/fcpClassic/FcpTable.html | This study used data from the 1000 Functional Connectomes Project (FCP). We thank the investigators and participants for making these data publicly available. | [11] |
| Healthy Brain Network (HBN) | https://fcon_1000.projects.nitrc.org/indi/cmi_healthy_brain_network/ | The Healthy Brain Network and its collaborative initiatives are supported by philanthropic contributions from the following individuals, foundations and organizations: Margaret Bilotti; Brooklyn Nets; Agapi and Bruce Burkard; James Chang; Phyllis Green and Randolph Cowen; Grieve Family Fund; Susan Miller and Byron Grote; Sarah and Geoff Gund; George Hall; Jonathan M. Harris Family Foundation; Joseph P. Healey; The Hearst Foundations; Eve and Ross Jaffe; Howard & Irene Levine Family Foundation; Rachael and Marshall Levine; George and Nitzia Logothetis; Christine and Richard Mack; Julie Minskoff; Valerie Mnuchin; Morgan Stanley Foundation; Amy and John Phelan; Roberts Family Foundation; Jim and Linda Robinson Foundation, Inc.; Linda and Richard Schaps; Zibby Schwarzman; Abigail Pogrebin and David Shapiro; Stavros Niarchos Foundation; Preethi Krishna and Ram Sundaram; Amy and John Weinberg; Donors to the 2013 Child Advocacy Award Dinner Auction; Donors to the 2012 Brant Art Auction. | [12] |
| NKIRS | https://fcon_1000.projects.nitrc.org/indi/enhanced/index.html | Principal support for the enhanced NKI-RS project is provided by the NIMH BRAINS R01MH094639-01 (PI Milham). Funding for key personnel also provided in part by the New York State Office of Mental Health and Research Foundation for Mental Hygiene. Funding for the decompression and augmentation of administrative and phenotypic protocols provided by a grant from the Child Mind Institute (1FDN2012-1). Additional personnel support provided by the Center for the Developing Brain at the Child Mind Institute, as well as NIMH R01MH081218, R01MH083246, and R21MH084126. Project support also provided by the NKI Center for Advanced Brain Imaging (CABI), the Brain Research Foundation, and the Stavros Niarchos Foundation. | [13,14] |
| SALD | https://fcon_1000.projects.nitrc.org/indi/retro/sald.html | This data repository was supported by the National Natural Science Foundation of China (31470981; 31571137; 31500885), National Outstanding young people plan, the Program for the Top Young Talents by Chongqing, the Fundamental Research Funds for the Central Universities (SWU1509383,SWU1509451,SWU1609177), Natural Science Foundation of Chongqing (cstc2015jcyjA10106), Fok Ying Tung Education Foundation (151023) , General Financial Grant from the China Postdoctoral Science Foundation (2015M572423, 2015M580767), Special Funds from the Chongqing Postdoctoral Science Foundation (Xm2015037, Xm2016044), Key research for Humanities and social sciences of Ministry of Education (14JJD880009). | [15] |
| SLIM | https://fcon_1000.projects.nitrc.org/indi/retro/southwestuni_qiu_index.html | This data repository was supported by: The National Natural Science Foundation of China (31271087; 31470981; 31571137; 31500885). National Outstanding young people planthe Program for the Top Young Talents by Chongqing, the Fundamental Research Funds for the Central Universities (SWU1509383,SWU1509451). Natural Science Foundation of Chongqing (cstc2015jcyjA10106). Fok Ying Tung Education Foundation (151023). General Financial Grant from the China Postdoctoral Science Foundation (2015M572423, 2015M580767). Special Funds from the Chongqing Postdoctoral Science Foundation (Xm2015037). Key research for Humanities and social sciences of Ministry of Education(14JJD880009). | [16] |
| Enhanced Nathan Kline Institute - Rockland Sample (NKI) | http://fcon_1000.projects.nitrc.org/ | Principal support for the enhanced NKI-RS project is provided by the NIMH BRAINS R01MH094639-01 (PI Milham). Funding for key personnel also provided in part by the New York State Office of Mental Health and Research Foundation for Mental Hygiene. Funding for the decompression and augmentation of administrative and phenotypic protocols provided by a grant from the Child Mind Institute (1FDN2012-1). Additional personnel support provided by the Center for the Developing Brain at the Child Mind Institute, as well as NIMH R01MH081218, R01MH083246, and R21MH084126. Project support also provided by the NKI Center for Advanced Brain Imaging (CABI), the Brain Research Foundation, and the Stavros Niarchos Foundation. | [17] |
| SRPBS-OPEN | https://bicr-resource.atr.jp/srpbsopen/ | Data used in the preparation of this work were obtained from the DecNef Project Brain Data Repository (https://bicr-resource.atr.jp/srpbsopen/) gathered by a consortium as part of the Japanese Strategic Research Program for the Promotion of Brain Science (SRPBS) supported by the Japanese Advanced Research and Development Programs for Medical Innovation (AMED) | [18] |

| | | | |
|---|---|---|---|
| SRPBS-1600 | https://bicr-resource.atr.jp/srpbs1600/ | Data used in the preparation of this work were obtained from the DecNef Project Brain Data Repository (https://bicr-resource.atr.jp/srpbsopen/) gathered by a consortium as part of the Japanese Strategic Research Program for the Promotion of Brain Science (SRPBS) supported by the Japanese Advanced Research and Development Programs for Medical Innovation (AMED) | 18 |
| HCP1200 | https://www.humanconnectome.org/study/hcp-young-adult/document/1200-subjects-data-release | Data were provided by the Human Connectome Project, WU-Minn Consortium (Principal Investigators: David Van Essen and Kamil Ugurbil; 1U54MH091657) funded by the 16 NIH Institutes and Centers that support the NIH Blueprint for Neuroscience Research; and by the McDonnell Center for Systems Neuroscience at Washington University. | 19 |
| HCP-Aging | https://www.humanconnectome.org/study/hcp-lifespan-aging | Data, methods used, and/or research reported in this publication were provided in whole or in part by the Aging Adult Vulnerability and Resiliency in the Aging Adult Brain Connectome (AABC) project (U19AG073585) and the Human Connectome Project in Aging (HCP-A, U01AG052564) funded by the National Institute of Aging of the National Institutes of Health. HCP-A was further supported by funds provided by the McDonnell Center for Neuroscience at Washington University in St. Louis. | 20 |
| HCP-Dev | https://www.humanconnectome.org/study/hcp-lifespan-development/ | Research reported in this publication was supported by the National Institute Of Mental Health of the National Institutes of Health under Award Number U01MH109589 and by funds provided by the McDonnell Center for Systems Neuroscience at Washington University in St. Louis. The HCP-Development 2.0 Release data used in this report came from DOI: 10.15154/1520708. | 21 |
| ADNI | https://adni.loni.usc.edu/ | Data collection and sharing for this project was funded by the Alzheimer's Disease Neuroimaging Initiative(ADNI) (National Institutes of Health Grant U01 AG024904) and DOD ADNI (Department of Defense awardnumber W81XWH-12-2-0012). ADNI is funded by the National Institute on Aging, the National Institute ofBiomedical Imaging and Bioengineering, and through generous contributions from the following: AbbVie,Alzheimer's Association; Alzheimer's Drug Discovery Foundation; Araclon Biotech; BioClinica, Inc.; Biogen;Bristol-Myers Squibb Company; CereSpir, Inc.; Cogstate; Eisai Inc.; Elan Pharmaceuticals, Inc.; Eli Lilly andCompany; EuroImmun; F. Hoffmann-La Roche Ltd and its affiliated company Genentech, Inc.; Fujirebio; GEHealthcare; IXICO Ltd.; Janssen Alzheimer Immunotherapy Research & Development, LLC.; Johnson &Johnson Pharmaceutical Research & Development LLC.; Lumosity; Lundbeck; Merck & Co., Inc.; MesoScale Diagnostics, LLC.; NeuroRx Research; Neurotrack Technologies; Novartis PharmaceuticalsCorporation; Pfizer Inc.; Piramal Imaging; Servier; Takeda Pharmaceutical Company; and TransitionTherapeutics. The Canadian Institutes of Health Research is providing funds to support ADNI clinical sitesin Canada. Private sector contributions are facilitated by the Foundation for the National Institutes of Health(www.fnih.org). The grantee organization is the Northern California Institute for Research and Education,and the study is coordinated by the Alzheimer's Therapeutic Research Institute at the University of SouthernCalifornia. ADNI data are disseminated by the Laboratory for Neuro Imaging at the University of SouthernCalifornia. | 22 |
| AIBL | https://aibl.org.au/ | Australian Imaging Biomarkers and Lifestyle flagship study of ageing (AIBL) was funded by the Commonwealth Scientific and Industrial Research Organisation (CSIRO), which was made available at the ADNI databas (http://www.loni.usc.edu/ADNI). The AIBL researchers contributed data but did not participate in analysis or writing of this report. AIBL researchers are listed at http://www.aibl.csiro.au. Correspondence should be addressed to Christopher Rowe. | 23 |

| | | | |
|---|---|---|---|
| GSP | https://dataverse.harvard.edu/dataverse/GSP | Data were obtained from the Brain Genomics Superstruct Project (GSP), supported by the National Institute of Mental Health (R01MH079799). | 24 |
| IXI | https://brain-development.org/ixi-dataset/ | | |
| Open Access Series of Imaging Studies (OASIS) | http://www.oasis-brains.org/ | Data were provided by OASIS 3: Longitudinal Multimodal Neuroimaging: Principal Investigators: T. Benzinger, D. Marcus, J. Morris; NIH P30 AG066444, P50 AG00561, P30 NS09857781, P01 AG026276, P01 AG003991, R01 AG043434, UL1 TR000448, R01 EB009352. AV-45 doses were provided by Avid Radiopharmaceuticals, a wholly owned subsidiary of Eli Lilly. Supported by grants P50 AG05681, P01 AG03991, R01 AG021910, P50 MH071616, U24 RR021382, R01 MH56584. | 25 |
| Pediatric Imaging, Neurocognition and Genetics (PING) | http://pingstudy.ucsd.edu/ | Data used in the preparation of this article were obtained from the Pediatric Imaging, Neurocognition and Genetics (PING) Study database (www.chd.ucsd.edu/research/ping-study.html, now shared through the NIMH Data Archive (NDA)). PING was a multisite, cross-sectional study that recruited more than 1,700 participants aged 3 to 20 years. The study was supported by award number RC2DA029475 from the National Institute on Drug Abuse with additional support for data sharing provided by the Eunice Kennedy Shriver National Institute of Child Health & Human Development under award number R01HD061414. A list of participating sites and study investigators can be found at https://ping-dataportal.ucsd.edu/sharing/Authors10222012.pdf. PING investigators designed and implemented the study and/or provided data but did not necessarily participate in analysis or writing of this report. This publication is solely the responsibility of the authors and does not necessarily represent the views of the National Institutes of Health or PING investigators. | 26 |
| PNC | https://www.med.upenn.edu/bbl/philadelphianeurodevelopmentalcohort.html | Data were obtained from the Philadelphia Neurodevelopmental Cohort (PNC), supported by the National Institute of Mental Health under award numbers RC2MH089983 and RC2MH089924. | 27 |
| PPMI | https://www.ppmi-info.org/ | Data used in the preparation of this article were obtained from the Parkinson's Progression Markers Initiative (PPMI) database (https://www.ppmi-info.org/access-data-specimens/download-data). For up-to-date information, see https://www.ppmi-info.org. PPMI, a public–private partnership, is funded by the Michael J. Fox Foundation for Parkinson's Research and its funding partners (https://www.ppmi-info.org/fundingpartners). | 28 |

| | | | |
|---|---|---|---|
| TCP | https://openneuro.org/datasets/ds005237 | This work was supported by the National Institute of Mental Health (R01MH123245 to AJH and R01MH120080 to AJH and BTTY). | [29] |
| OpenNeuro | https://openneuro.org/datasets/ | These sub-datasets were obtained from the OpenNeuro database, including ds000030, ds000053, ds000119, ds000202, ds000208, ds000222, ds000228, ds000240, ds000243, ds001131, ds001408, ds001486, ds001734, ds001747, ds001748, ds001796, ds001838, ds001848, ds001894, ds002116, ds002236, ds002330, ds002345, ds002382, ds002385, ds002424, ds002643, ds002647, ds002717, ds002731, ds002785, ds002790, ds002837, ds002843, ds002886, ds003037, ds003097, ds003126, ds003138, ds003242, ds003346, ds003416, ds003436, ds003469, ds003481, ds003499, ds003508, ds003568, ds003592, ds003612, ds003643, ds003701, ds003709, ds003717, ds003745, ds003798, ds003826, ds003831, ds003877, ds003974, ds003988, ds004044, ds004146, ds004169, ds004173, ds004199, ds004215, ds004217, ds004261, ds004285, ds004302, ds004349, ds004466, ds004469, ds004512, ds004556, ds004589, ds004604, ds004636, ds004648, ds004697, ds004711, ds004718, ds004725, ds004746, ds004837. | [30–120] |

### 14. Supplementary Table S2. Overview of the included datasets

| | Train Set | | | | Test Set | | | | Clinical Set | | | |
|---|---|---|---|---|---|---|---|---|---|---|---|---|
| Site | N | F/M | Age (mean) | Age (std) | N | F/M | Age (mean) | Age (std) | N | F/M | Age (mean) | Age (std) |
| ABIDE-NYU-Siemens_Allegra_3T | 51 | 39/12 | 15.47 | 6.39 | 54 | 40/14 | 16.13 | 6.16 | 79 | 68/11 | 14.52 | 6.97 |
| ABIDE-UM_1-GE_Signa_3T | 26 | 18/8 | 14.08 | 3.17 | 29 | 20/9 | 14.06 | 3.25 | 55 | 46/9 | 12.72 | 2.4 |
| ABIDE-USM-Siemens_TrioTim_3T | 22 | 22/0 | 21.16 | 7.67 | 21 | 21/0 | 21.58 | 7.81 | 58 | 58/0 | 22.65 | 7.73 |
| ABIDE2-ABIDEII-KKI_1-Philips_Achieva_3.0T | 34 | 24/10 | 10.23 | 1.04 | 43 | 30/13 | 10.25 | 1.1 | 56 | 41/15 | 10.31 | 1.51 |
| ADHD200-KKI- | 22 | 13/9 | 10.55 | 1.13 | 21 | 12/9 | 10.38 | 1.42 | 22 | 12/10 | 10.22 | 1.56 |
| ADHD200-NYU-SIEMENS_MAGNETOM_Allegra_syngo_MR_2004A | 36 | 15/21 | 12.12 | 3.08 | 29 | 15/14 | 12.04 | 3.22 | 150 | 114/36 | 10.96 | 2.68 |
| ADHD200-Peking_1-SIEMENS_MAGNETOM_TrioTim_syngo_MR_B15 | 28 | 7/21 | 11.09 | 1.72 | 35 | 12/23 | 11.03 | 1.77 | 48 | 36/12 | 11.24 | 2.13 |
| ADHD200-Pittsburgh-SIEMENS_MAGNETOM_TrioTim_syngo_MR_B15 | 24 | 13/11 | 14.12 | 2.66 | 23 | 15/8 | 14.54 | 2.61 | 4 | 3/1 | 15.4 | 1.4 |
| AIBL-Brain Research Institute-Siemens_TrioTim | 42 | 18/24 | 73.52 | 7.3 | 47 | 23/24 | 74.49 | 6.2 | 64 | 26/38 | 75.95 | 7.56 |
| AIBL-Royal Melbourne Hosp 3T-Siemens_TrioTim | 42 | 14/28 | 73.57 | 6.06 | 44 | 20/24 | 72.52 | 5.51 | 15 | 11/4 | 74.4 | 5.96 |
| AIBL-SKG Radiology Subiaco-Siemens_Verio | 24 | 6/18 | 71.75 | 5.05 | 18 | 4/14 | 73.22 | 5.15 | 14 | 7/7 | 75.71 | 6.58 |
| CAMCAN | 647 | 319/328 | 54.65 | 18.6 | | | | | | | | |
| COBRE-SIEMENS_TrioTim | 42 | 29/13 | 35.9 | 12.26 | 32 | 22/10 | 35.72 | 10.81 | 72 | 58/14 | 38.17 | 13.89 |
| CORR-BMB_1-Siemens_TrioTim | 50 | 24/26 | 30.83 | 7.09 | | | | | | | | |
| CORR-BNU_1-Siemens_TrioTim | 53 | 26/27 | 23 | 2.3 | | | | | | | | |
| CORR-BNU_2-Siemens_TrioTim | 61 | 33/28 | 21.32 | 0.86 | | | | | | | | |
| CORR-BNU_3-Siemens_TrioTim | 48 | 24/24 | 22.54 | 2.15 | | | | | | | | |
| CORR-IPCAS_7-Siemens_TrioTim | 70 | 30/40 | 11.67 | 3.07 | | | | | | | | |
| CORR-MRN-Siemens_TrioTim | 54 | 27/27 | 24.88 | 11.02 | | | | | | | | |
| CORR-NYU_2-Siemens_Allegra | 185 | 115/70 | 20.22 | 11.53 | | | | | | | | |
| CORR-SWU_4-Siemens_TrioTim | 234 | 119/115 | 20.04 | 1.28 | | | | | | | | |
| CORR-UM-Siemens_TrioTim | 80 | 22/58 | 65.36 | 6.26 | | | | | | | | |
| CORR-UPSM_1-Siemens_TrioTim | 100 | 52/48 | 15.14 | 2.8 | | | | | | | | |

| | | | | | | | | | | | | |
|---|---|---|---|---|---|---|---|---|---|---|---|---|
| FCON1000_Beijing_Zang | 196 | 196/0 | 21.16 | 1.84 | | | | | | | | |
| FCON1000_Cambridge_Buckner | 197 | 197/0 | 21.04 | 2.32 | | | | | | | | |
| FCON1000_ICBM | 85 | 85/0 | 44.02 | 17.96 | | | | | | | | |
| FCON1000_Milwaukee_b | 46 | 46/0 | 53.59 | 5.79 | | | | | | | | |
| FCON1000_Oulu | 102 | 102/0 | 21.52 | 0.58 | | | | | | | | |
| GSP-B | 218 | 98/120 | 22.31 | 3.26 | | | | | | | | |
| GSP-D | 58 | 22/36 | 21.9 | 2.88 | | | | | | | | |
| GSP-E | 697 | 302/395 | 21.29 | 2.66 | | | | | | | | |
| HBN-Site-CBIC-Siemens_Prisma_fit | 157 | 95/62 | 10.48 | 3.72 | 149 | 93/56 | 10.15 | 3.45 | 923 | 637/286 | 10.4 | 3.4 |
| HBN-Site-CUNY-Siemens_Prisma | 113 | 66/47 | 10.58 | 3.57 | 116 | 74/42 | 10.58 | 3.31 | 262 | 173/89 | 10.14 | 3.13 |
| HBN-Site-RU-Siemens_TrioTim | 110 | 65/45 | 10.08 | 3.64 | 103 | 67/36 | 10.3 | 3.93 | 634 | 429/205 | 10.56 | 3.48 |
| HCP1200 | 1109 | 503/606 | 28.81 | 3.7 | | | | | | | | |
| HCPAging-MGH-Siemens_Prisma_fit | 162 | 79/83 | 61.56 | 16.01 | | | | | | | | |
| HCPAging-UCLA-Siemens_Prisma_fit | 147 | 63/84 | 56.76 | 13.66 | | | | | | | | |
| HCPAging-UMinn-Siemens_Prisma | 202 | 90/112 | 61.68 | 16.79 | | | | | | | | |
| HCPAging-WashU-Siemens_Prisma | 186 | 80/106 | 61.04 | 15.88 | | | | | | | | |
| HCPDev-Harvard-Siemens_Prisma_fit | 209 | 97/112 | 14.94 | 3.87 | | | | | | | | |
| HCPDev-UCLA-Siemens_Prisma_fit | 127 | 59/68 | 13.96 | 4.03 | | | | | | | | |
| HCPDev-UMinn-Siemens_Prisma | 163 | 71/92 | 13.95 | 3.91 | | | | | | | | |
| HCPDev-WashU-Siemens_Prisma | 137 | 69/68 | 14.57 | 4.45 | | | | | | | | |
| IXI-Guy's_Hospital_1.5T_Philips | 313 | 138/175 | 50.71 | 15.94 | | | | | | | | |
| IXI-Hammersmith_Hospital_3T_Philips | 181 | 87/94 | 47.36 | 16.71 | | | | | | | | |
| IXI-Institute_of_Psychiatry_1.5T_GE | 68 | 24/44 | 42.38 | 16.6 | | | | | | | | |
| NKIRS_Siemens_TrioTim | 235 | 86/149 | 41.98 | 22.23 | 247 | 83/164 | 43.52 | 22.15 | 58 | 24/34 | 39.4 | 20.34 |
| OASIS3-OASIS3-Siemens_Biograph-mMR_3.0T | 106 | 51/55 | 69.24 | 7.86 | 104 | 50/54 | 70.04 | 7.79 | 75 | 34/41 | 75.18 | 6.48 |
| OASIS3-OASIS3-Siemens_MAGNETOM-Vida_3.0T | 42 | 17/25 | 69.18 | 7.97 | 55 | 28/27 | 70.86 | 8.35 | 26 | 16/10 | 72.99 | 6.85 |
| OASIS3-OASIS3-Siemens_TrioTim_3.0T | 275 | 111/164 | 68.46 | 9.49 | 295 | 120/175 | 69.22 | 9.36 | 201 | 114/87 | 76.15 | 7.95 |
| OpenNeuro-NIH_FMRIF-GE_DISCOVERY_MR750 | 29 | 11/18 | 15.66 | 1.11 | 23 | 12/11 | 16 | 1.21 | 64 | 19/45 | 16.67 | 1.11 |
| PING-PING-Achieva | 64 | 34/30 | 13.37 | 4.36 | | | | | | | | |
| PING-PING-DISCOVERY MR750 | 121 | 55/66 | 8.55 | 3.37 | | | | | | | | |

| | | | | | | | | | | | | |
|---|---|---|---|---|---|---|---|---|---|---|---|---|
| PING-PING-PING | 118 | 61/57 | 14.94 | 4.29 | | | | | | | | |
| PING-PING-SIGNA HDx | 103 | 57/46 | 14.58 | 4.17 | | | | | | | | |
| PING-PING-TrioTim | 354 | 188/166 | 11.8 | 5.17 | | | | | | | | |
| PNC_HUP6_Siemens_TrioTim | 1424 | 696/728 | 14.53 | 3.54 | | | | | | | | |
| PNC_MR6_Siemens_TrioTim | 159 | 58/101 | 14.61 | 3.5 | | | | | | | | |
| PPMI_18_GE-SIGNA-Architect | 54 | 25/29 | 67.18 | 5.23 | | | | | | | | |
| PPMI_307_Siemens-PMOD | 47 | 27/20 | 62.13 | 10.62 | | | | | | | | |
| PPMI_34_Siemens-PMOD | 112 | 41/71 | 62.64 | 7.94 | | | | | | | | |
| PPMI_34_Siemens-Verio | 103 | 36/67 | 66.13 | 5.38 | | | | | | | | |
| PPMI_73_Siemens-PMOD | 71 | 30/41 | 61.74 | 6.15 | | | | | | | | |
| PPMI_88_Siemens-Prisma-fit | 49 | 21/28 | 65.29 | 6.63 | | | | | | | | |
| SRPBS_1600-ATR-SIEMENS_Verio | 77 | 60/17 | 22.68 | 1.98 | | | | | | | | |
| SRPBS_1600-Center_of_Innovation_in_Hiroshima_University-SIEMENS_MAGNETOM_Verio.Dot | 62 | 25/37 | 51.06 | 14.18 | 62 | 21/41 | 52.66 | 12.73 | 68 | 29/39 | 44.9 | 12.43 |
| SRPBS_1600-Hiroshima_University_Hospital-GE_Signa_HDxt | 55 | 21/34 | 38.65 | 13.04 | 61 | 21/40 | 36.8 | 12.72 | 73 | 38/35 | 42.71 | 12.01 |
| SRPBS_1600-Kyoto_university-SIEMENS_TimTrio | 79 | 43/36 | 36.57 | 13.3 | 80 | 50/30 | 36.45 | 13.95 | 61 | 31/30 | 41.7 | 11.18 |
| SRPBS_1600-Kyoto_university-SIEMENS_Trio | 33 | 20/13 | 29.48 | 8.24 | 42 | 28/14 | 28.43 | 9.72 | 47 | 26/21 | 37.89 | 9.79 |
| SRPBS_1600-Showa_university-SIEMENS_Verio | 47 | 38/9 | 28.19 | 8.38 | 53 | 47/6 | 28.74 | 7.49 | 133 | 114/19 | 33.68 | 8.74 |
| SRPBS_1600-University_of_Tokyo-GE_Discovery_MR750w | 92 | 43/49 | 35.52 | 17.45 | 78 | 35/43 | 35.67 | 17.67 | 108 | 69/39 | 36.14 | 11.45 |
| TCP-1-Siemens_Magnetom_3T_Prisma | 30 | 12/18 | 30.4 | 12.47 | 28 | 13/15 | 31.57 | 13.62 | 10 | 3/7 | 40.3 | 18.86 |
| ds000030-ds000030-35343 | 57 | 32/25 | 31.79 | 8.8 | 45 | 23/22 | 31.6 | 9.06 | 46 | 25/21 | 35.15 | 9.85 |
| ds000053-ds000053-SIEMENS_Skyra | 59 | 28/31 | 22.92 | 3.37 | | | | | | | | |
| ds000119-ds000119-Siemens_Allegra_3T | 69 | 29/40 | 16.16 | 4.77 | | | | | | | | |
| ds000202-ds000202-Philips_Achieva_3T | 87 | 0/87 | 22.15 | 2.74 | | | | | | | | |
| ds000208-Northwestern_University_Chicago-Siemens_Trio_3T | 75 | 35/40 | 57.95 | 6.88 | | | | | | | | |
| ds000222-University_Medicine_Berlin-Siemens_Magnetom_Trio_3T | 68 | 30/38 | 45.16 | 20.11 | | | | | | | | |
| ds000228-ds000228-Siemens_TrioTim | 155 | 71/84 | 10.56 | 8.07 | | | | | | | | |
| ds000240-SC3T-Siemens_Prisma | 63 | 28/35 | 48.98 | 24.41 | | | | | | | | |
| ds000243-Washington_University_St_Louis-Siemens_MAGNETOM_Tim_Trio_3T | 120 | 59/61 | 24.74 | 2.39 | | | | | | | | |

| | | | | | | | | | | | | |
|---|---|---|---|---|---|---|---|---|---|---|---|---|
| ds001131-ds001131-Siemens_Skyra_3T | 54 | 27/27 | 21.02 | 3.44 | | | | | | | | |
| ds001408-Federal_Treatment_Rehab_Center_Moscow-General Electrics_Discovery_MR750 | 42 | 17/25 | 26.69 | 6.42 | | | | | | | | |
| ds001486-ds001486-Siemens_Trio_Tim_3T | 132 | 62/70 | 11.26 | 1.46 | | | | | | | | |
| ds001734-ds001734-Siemens_Prisma_3T | 108 | 48/60 | 25.55 | 3.59 | | | | | | | | |
| ds001747-Brigham_Young_University-Siemens_TrioTim | 91 | 40/51 | 21.67 | 2.51 | | | | | | | | |
| ds001748-St_Lucia_Campus-Siemens_TrioTim | 42 | 22/20 | 18.83 | 8.49 | | | | | | | | |
| ds001796-UoR-Siemens_Prisma_fit | 62 | 14/48 | 32.1 | 7.62 | | | | | | | | |
| ds001838-Robarts_Research_Institute-Siemens_Prisma_fit | 52 | 26/26 | 24.25 | 4.23 | | | | | | | | |
| ds001848-Robarts_Research_Institute-Siemens_Prisma_fit | 52 | 22/30 | 23.56 | 4.34 | | | | | | | | |
| ds001894-Northwestern_CAMRI-Siemens_Magnetom_Trio_Tim_3T | 188 | 99/89 | 10.48 | 1.61 | | | | | | | | |
| ds002116-Robarts_Research_Institute-Siemens_Prisma_fit | 57 | 35/22 | 9.32 | 2.76 | | | | | | | | |
| ds002236-Evanston_Hospital-GE_SIGNA_EXCITE | 91 | 52/39 | 11.36 | 2.08 | | | | | | | | |
| ds002330-Queens_University_Ontario-Siemens_Magnetom_Tim_Trio | 65 | 29/36 | 26.55 | 4.3 | | | | | | | | |
| ds002345-Princeton University-Siemens_Prisma | 120 | 43/77 | 22.53 | 5.48 | | | | | | | | |
| ds002345-Princeton University-Siemens_Skyra | 214 | 93/121 | 21.6 | 4.14 | | | | | | | | |
| ds002382-Washington_University_St_Louis-Siemens_Prisma_3T | 60 | 24/36 | 47.95 | 23.84 | | | | | | | | |
| ds002385-UPMC_Presbyterian_University_Hospital-Siemens_Biograph_mMR | 127 | 56/71 | 19.37 | 5.07 | | | | | | | | |
| ds002424-ds002424-ds002424 | 23 | 16/7 | 10.67 | 0.92 | 21 | 14/7 | 10.26 | 1.01 | 35 | 35/0 | 10.29 | 0.94 |
| ds002643-Spinoza Centre for Neuroimaging, location REC-Philips_Achieva_3T | 78 | 38/40 | 24.81 | 4.92 | | | | | | | | |
| ds002647-University_of_Southern_California-Siemens_Prisma_fit_3T | 80 | 0/80 | 34.83 | 20.05 | | | | | | | | |
| ds002717-Weizmann-Siemens_TrioTim | 55 | 0/55 | 35.2 | 5.67 | | | | | | | | |
| ds002731-Beijing Normal University-Siemens | 59 | 31/28 | 21.25 | 1.45 | | | | | | | | |
| ds002785-PIOP1-Philips_Achieva_3T | 208 | 88/120 | 22.19 | 1.8 | | | | | | | | |
| ds002790-PIOP2-Philips_Achieva_dStream_3T | 224 | 96/128 | 21.96 | 1.79 | | | | | | | | |
| ds002837-PIOP2-Philips_Achieva_dStream_3T | 86 | 44/42 | 26.73 | 10.06 | | | | | | | | |
| ds002843-University_of_Pennsylvania-Siemens_Trio_3T | 166 | 98/68 | 24.52 | 4.49 | | | | | | | | |
| ds002886-CAMRI-Siemens_Trio_Tim_3T | 56 | 24/32 | 11.19 | 1.64 | | | | | | | | |

| | | | | |
|---|---|---|---|---|
| ds003097-ID-1000-Philips_Intera_3T | 925 | 444/481 | 22.85 | 1.71 |
| ds003126-ID-1000-Philips_Intera_3T | 58 | 35/23 | 9.05 | 0.69 |
| ds003138-KFU-Siemens_Skyra | 53 | 22/31 | 23.92 | 3.5 |
| ds003242-MIT-Siemens_Prisma_fit | 89 | 30/59 | 27.1 | 5.41 |
| ds003346-I.N._Psiquiatria-Philips_Ingenia | 61 | 50/11 | 30.43 | 8.22 |
| ds003416-1B-Philips_Achieva_3T | 90 | 52/38 | 8.06 | 5.26 |
| ds003436-Az.Osp._Udine-Philips_Achieva | 49 | 19/30 | 23.14 | 3.76 |
| ds003469-ds003469-ds003469 | 81 | 31/50 | 24.06 | 5.31 |
| ds003481-INSTITUTO_DE_NEUROBIOLOGIA-GE_DISCOVERY_MR750 | 143 | 66/77 | 23.43 | 3.72 |
| ds003499-NYU_Center_for_Brain_Imaging-Siemens_Prisma | 90 | 43/47 | 15.92 | 5.02 |
| ds003508-Lund MR 7T-Philips_Achieva | 56 | 15/41 | 22.75 | 2.04 |
| ds003568-NIH_FMRIF-GE_DISCOVERY_MR750 | 49 | 18/31 | 15.88 | 1.78 |
| ds003592-ds003592-ds003592 | 277 | 120/157 | 40.09 | 22.94 |
| ds003612-TU_Graz-Siemens_MAGNETOM_Vida | 50 | 25/25 | 24.08 | 2.8 |
| ds003643-ds003643-ds003643 | 80 | 38/42 | 20.71 | 3.09 |
| ds003701-RUBIC-Siemens_TrioTim | 80 | 36/44 | 22.69 | 4.05 |
| ds003709-NIH_FMRIF-GE_DISCOVERY_MR750 | 51 | 11/40 | 15.57 | 1.36 |
| ds003717-ds003717-ds003717 | 60 | 15/45 | 22.42 | 3.24 |
| ds003745-Temple_University_-_Weiss_Hall-Siemens_Prisma | 50 | 23/27 | 45.28 | 23.58 |
| ds003798-Caltech Tim32-Siemens_TrioTim | 73 | 41/32 | 27.16 | 4.95 |
| ds003826-Malopolskie_Centrum_Biotechnologii-Siemens_Skyra | 108 | 40/68 | 24.3 | 3.52 |
| ds003831 | 68 | 22/46 | 38.81 | 14.31 |
| ds003877 | 54 | 27/27 | 6.65 | 0.43 |
| ds003974 | 42 | 19/23 | 22.74 | 4.83 |
| ds003988 | 46 | 22/24 | 25.8 | 4.88 |
| ds004044-BeiJing_Normal_University-Siemens_Prisma | 55 | 25/30 | 22.65 | 2.16 |
| ds004146-St_Lucia_Campus-Siemens_Prisma_fit | 381 | 194/187 | 10.93 | 1.43 |
| ds004169-ds004169-Siemens_MedSpec_4T | 1114 | 434/680 | 21.11 | 4.09 |
| ds004173-RCNS-Siemens_Prisma | 109 | 36/73 | 28.08 | 11.26 |

| | | | | |
|---|---|---|---|---|
| ds004199-Life&Brain - NeuroCognition-3T-Siemens_TrioTim | 114 | 61/53 | 6.94 | 2.48 |
| ds004215-NIH FMRIF-GE_DISCOVERY_MR750 | 155 | 53/102 | 34.05 | 12.75 |
| ds004217-Princeton_University_-Neuroscience_Institute-Siemens_Skyra | 53 | 20/33 | 20.6 | 3.36 |
| ds004261-BeiJing_Normal_University-Siemens_Prisma | 55 | 26/29 | 21.49 | 1.95 |
| ds004285-ds004285-ds004285 | 56 | 14/42 | 42.11 | 23.85 |
| ds004349-University_of_Oregon-Siemens_Skyra | 50 | 19/31 | 19.6 | 1.99 |
| ds004466-CISC-Siemens_Prisma | 123 | 28/95 | 35.67 | 13.3 |
| ds004469-ds004469-Philips_Achieva | 66 | 18/48 | 31.83 | 11.84 |
| ds004512-Palmetto_Health_Richland-Siemens_Prisma_fit | 170 | 100/70 | 61.58 | 11.46 |
| ds004556-KFU-Siemens_Skyra | 53 | 29/24 | 29.98 | 7.03 |
| ds004589-Brigham_Young_University-Siemens_TrioTim | 98 | 43/55 | 22.48 | 3.84 |
| ds004604-ULB_ERASME-GE_SIGNA_PET_MR | 50 | 27/23 | 33.62 | 10.49 |
| ds004636-ds004636-ds004636 | 106 | 37/69 | 23.86 | 5.62 |
| ds004648-IATM-Philips_Ingenia | 60 | 60/0 | 28.35 | 5.14 |
| ds004697-PSU-SLEIC-Siemens_Prisma_fit | 83 | 41/42 | 7.81 | 0.59 |
| ds004711-ds004711-ds004711 | 185 | 88/97 | 44.94 | 19.23 |
| ds004718-The Hong Kong Polytechnic University-Siemens_MAGNETOM_Prisma | 50 | 11/39 | 69.08 | 3.58 |
| ds004725-ds004725-ds004725 | 84 | 40/44 | 46.79 | 24.87 |
| ds004746-Intermountain_Neuroimaging_Consortium-Siemens_Prisma_fit | 390 | 162/228 | 36.12 | 3.4 |
| ds004837-ds004837-ds004837 | 59 | 40/19 | 23.72 | 4.47 |

15. Supplementary Table S3. Cross-sectional and longitudinal shape errors between reconstructed surface and the ground truth in observations

| Shape errors | m-s-rep | cm-rep | ds-rep | SPHARM-PDM |
|---|---|---|---|---|
| **t0-Q (mm)** | 0.003±0.002 | 0.05±0.13 | 0.13±0.11 | 0.01±0.001 |
| **t0-q (mm)** | 0.78±0.07 | 4.22±38.43 | 4.22±38.43 | 1.07±0.16 |
| **t0-HD (mm)** | 1.63±0.31 | 19.11±8.61 | 10.59±8.36 | 15.88±9.88 |
| **t0-Ae ($mm^2$)** | 0.83±0.43 | 12.40±14.31 | 29.85±16.44 | 1.73±0.48 |
| **t0-Ce (1/mm)** | 0.08±0.05 | 0.10±0.53 | 0.18±0.54 | 0.10±0.05 |
| **t1-Q (mm)** | 0.003±0.001 | 0.05±0.10 | 0.14±0.11 | 0.01±0.001 |
| **t1-q (mm)** | 0.73±0.05 | 3.55±29.50 | 8.82±9.94 | 1.02±0.16 |
| **t1-HD (mm)** | 1.63±0.48 | 18.54±30.07 | 10.80±8.55 | 15.77±9.84 |
| **t1-Ae ($mm^2$)** | 0.82±0.46 | 13.18±13.45 | 33.34±12.45 | 1.78±0.50 |
| **t1-Ce (1/mm)** | 0.06±0.08 | 0.07±0.07 | 0.07±0.07 | 0.06±0.04 |
| **t2-Q (mm)** | 0.003±0.001 | 0.03±0.02 | 0.14±0.11 | 0.01±0.001 |
| **t2-q (mm)** | 0.70±0.06 | 1.34±1.13 | 9.08±10.05 | 1.07±1.09 |
| **t2-HD (mm)** | 1.62±0.42 | 16.17±9.86 | 11.01±8.65 | 15.62±9.95 |
| **t2-Ae ($mm^2$)** | 0.82±0.49 | 13.27±13.85 | 32.94±12.42 | 1.60±3.30 |
| **t2-Ce (1/mm)** | 0.05±0.08 | 0.06±0.07 | 0.14±0.03 | 0.06±0.02 |

### 16. Supplementary Table S4. Correspondence error of different shape models

| Location | ARMM | cm-rep | ds-rep | SPHARM-PDM |
|---|---|---|---|---|
| **1** | 1.09 ± 0.51 mm | 6.25 ± 6.19 mm | 3.50 ± 0.92 mm | 34.05 ± 16.06 mm |
| **2** | 1.14 ± 0.54 mm | 3.30 ± 3.87 mm | 3.79 ± 1.03 mm | 6.84 ± 2.63 mm |
| **3** | 1.55 ± 0.61 mm | 4.12 ± 6.63 mm | 2.99 ± 1.22 mm | 29.80 ± 11.95 mm |
| **4** | 1.66 ± 0.78 mm | 4.64 ± 5.05 mm | 3.41 ± 1.42 mm | 9.97 ± 6.06 mm |

### 17. Supplementary Table S5.Test-Retest Reliability of ARMM Measurements

| Side | Category | MeanICC | VarICC | GlobalICC |
|---|---|---|---|---|
| **Left** | InfThickness | 0.6060 | 0.0332 | 0.7675 |
| **Left** | SupThickness | 0.5558 | 0.0284 | 0.8163 |
| **Left** | LatWidth | 0.9136 | 0.0161 | 0.9679 |
| **Left** | VenWidth | 0.9272 | 0.0058 | 0.9715 |
| **Left** | Width | 0.9260 | 0.0094 | 0.9733 |
| **Left** | Length | 0.9674 | 0.0002 | 0.9699 |
| **Right** | InfThickness | 0.6291 | 0.0327 | 0.7497 |
| **Right** | SupThickness | 0.5642 | 0.0287 | 0.8177 |
| **Right** | LatWidth | 0.9235 | 0.0217 | 0.9861 |
| **Right** | VenWidth | 0.9345 | 0.0059 | 0.9833 |
| **Right** | Width | 0.9344 | 0.0121 | 0.9882 |
| **Right** | Length | 0.9530 | 0.0011 | 0.9586 |

### 18. Supplementary Table S6. Test-Retest Reliability of Hippounfold Measurements

| Side | Category | MeanICC | VarICC | GlobalICC |
|---|---|---|---|---|
| **Left** | hipp_thickness | 0.5878 | 0.0480 | 0.7869 |
| **Left** | hipp_curvature | 0.2008 | 0.0933 | 0.0259 |
| **Left** | hipp_surfarea | 0.5739 | 0.0534 | 0.9928 |
| **Left** | hipp_gyrification | 0.5739 | 0.0534 | 0.9760 |
| **Left** | dentate_curvature | 0.1912 | 0.0902 | 0.0959 |
| **Left** | dentate_gyrification | 0.4776 | 0.0849 | 0.9559 |
| **Left** | dentate_surfarea | 0.4776 | 0.0849 | 0.9585 |
| **Right** | hipp_thickness | 0.5621 | 0.0680 | 0.7587 |
| **Right** | hipp_curvature | 0.2072 | 0.0989 | 0.7042 |
| **Right** | hipp_surfarea | 0.5696 | 0.0592 | 0.9968 |
| **Right** | hipp_gyrification | 0.5696 | 0.0592 | 0.9823 |
| **Right** | dentate_curvature | 0.1271 | 0.0952 | 0.9350 |
| **Right** | dentate_gyrification | 0.4803 | 0.0743 | 0.8483 |
| **Right** | dentate_surfarea | 0.4803 | 0.0743 | 0.9229 |

### 19. Supplementary Table S7. Comparison of model fitting across different function configs

| | **config-1** | **config-2** | **config-3** | **config-4** | **config-5** | **config-6** | **config-7** | **config-8** | **config-9** | **config-10** | **config-11** | **config-12** |
|---|---|---|---|---|---|---|---|---|---|---|---|---|
| **Z Scores Mean** | -0.025 | -0.023 | -0.023 | -0.023 | -0.023 | **-0.021** | -0.023 | -0.023 | -0.023 | -0.022 | -0.022 | -0.022 |
| **Z Scores SD** | 0.980 | **0.989** | 0.978 | 0.982 | 0.982 | 0.981 | 0.980 | 0.989 | 0.978 | 0.982 | 0.983 | 0.981 |
| **Skewness** | 0.068 | 0.063 | 0.064 | 0.064 | 0.057 | 0.060 | 0.063 | 0.063 | 0.063 | 0.068 | 0.067 | 0.060 |
| **Excess Kurtosis** | 0.431 | 0.391 | 0.426 | 0.446 | 0.418 | 0.432 | 0.412 | 0.391 | 0.423 | 0.462 | 0.448 | 0.431 |
| **Normality Test Statistic** | **0.989** | 0.988 | **0.989** | **0.989** | **0.989** | **0.989** | **0.989** | 0.988 | **0.989** | **0.989** | **0.989** | **0.989** |
| **Logarithmic Score** | -0.617 | -0.628 | -0.628 | -0.629 | -0.628 | -0.627 | -0.628 | -0.628 | -0.628 | -0.627 | -0.627 | -0.626 |
| **R Squared** | **0.032** | 0.030 | 0.030 | 0.030 | 0.030 | 0.030 | 0.031 | 0.030 | 0.031 | **0.032** | 0.030 | 0.030 |
| **MAE** | **0.537** | 0.541 | 0.541 | 0.545 | 0.541 | 0.541 | 0.541 | 0.541 | 0.541 | 0.544 | 0.541 | 0.541 |
| **MSE** | 1.272 | 1.260 | 1.260 | 1.282 | 1.260 | 1.260 | **1.259** | 1.260 | **1.259** | 1.280 | **1.259** | 1.260 |
| **RMSE** | **0.752** | 0.754 | 0.753 | 0.760 | 0.753 | 0.754 | 0.753 | 0.753 | 0.753 | 0.758 | 0.753 | 0.753 |
| **AIC** | **27700.987** | 28214.539 | 28140.189 | 28178.635 | 28097.666 | 28108.591 | 28158.149 | 28213.986 | 28139.795 | 28110.416 | 28097.682 | 28110.220 |
| **BIC** | **28934.369** | 29414.394 | 29369.523 | 29471.550 | 29359.678 | 29360.250 | 29374.198 | 29408.063 | 29360.765 | 29406.952 | 29354.203 | 29351.139 |

#### 20. Supplementary Table S8. Group-wise comparison of normative z-scores between ASD and test HC

| | Left | | | Right | | |
|---|---|---|---|---|---|---|
| | effect size | P | corrected P | effect size | P | corrected P |
| Lamellar Width 1 | 0.050 | 0.034 | 0.108 | 0.026 | 0.269 | 0.404 |
| Lamellar Width 2 | -0.003 | 0.887 | 0.947 | -0.001 | 0.972 | 0.985 |
| Lamellar Width 3 | -0.045 | 0.055 | 0.144 | -0.020 | 0.390 | 0.543 |
| Lamellar Width 4 | -0.049 | 0.038 | 0.114 | -0.144 | 0.000 | 0.000 |
| Lamellar Width 5 | -0.054 | 0.023 | 0.081 | -0.045 | 0.056 | 0.144 |
| Lamellar Width 6 | -0.036 | 0.129 | 0.254 | -0.001 | 0.978 | 0.989 |
| Lamellar Width 7 | 0.015 | 0.532 | 0.676 | 0.012 | 0.600 | 0.729 |
| Lamellar Width 8 | -0.002 | 0.947 | 0.985 | -0.015 | 0.531 | 0.676 |
| Lamellar Width 9 | -0.014 | 0.538 | 0.680 | -0.023 | 0.326 | 0.461 |
| Lamellar Width 10 | 0.019 | 0.408 | 0.554 | -0.012 | 0.611 | 0.736 |
| Lamellar Width 11 | 0.028 | 0.232 | 0.361 | -0.010 | 0.665 | 0.778 |
| Lamellar Width 12 | 0.042 | 0.077 | 0.179 | 0.003 | 0.888 | 0.947 |
| Lamellar Width 13 | 0.006 | 0.791 | 0.883 | 0.010 | 0.661 | 0.777 |
| Lamellar Width 14 | -0.024 | 0.318 | 0.453 | -0.020 | 0.402 | 0.550 |
| Lamellar Width 15 | -0.028 | 0.233 | 0.361 | -0.008 | 0.729 | 0.843 |
| Lamellar Width 16 | -0.001 | 0.969 | 0.985 | -0.008 | 0.741 | 0.853 |
| Lamellar Width 17 | -0.004 | 0.863 | 0.938 | 0.020 | 0.400 | 0.550 |
| Lamellar Width 18 | 0.032 | 0.171 | 0.306 | 0.045 | 0.057 | 0.144 |
| Lamellar Width 19 | 0.033 | 0.155 | 0.288 | 0.071 | 0.003 | 0.015 |
| Lamellar Width 20 | 0.036 | 0.128 | 0.252 | 0.034 | 0.145 | 0.275 |
| Lamellar Width 21 | 0.061 | 0.010 | 0.039 | 0.008 | 0.749 | 0.854 |
| Lamellar Width 22 | 0.083 | 0.000 | 0.003 | 0.038 | 0.106 | 0.219 |
| Lamellar Width 23 | 0.091 | 0.000 | 0.001 | 0.046 | 0.050 | 0.138 |
| Lamellar Width 24 | 0.095 | 0.000 | 0.001 | 0.038 | 0.104 | 0.216 |
| Lamellar Width 25 | 0.044 | 0.063 | 0.155 | 0.026 | 0.264 | 0.402 |
| Lamellar Width 26 | 0.040 | 0.087 | 0.193 | 0.005 | 0.832 | 0.908 |
| Lamellar Width 27 | 0.067 | 0.004 | 0.021 | 0.008 | 0.744 | 0.853 |

| | | | | | | |
|---|---|---|---|---|---|---|
| Lamellar Width 28 | 0.114 | 0.000 | 0.000 | 0.046 | 0.053 | 0.141 |
| Lamellar Width 29 | 0.099 | 0.000 | 0.000 | 0.021 | 0.363 | 0.507 |
| Lamellar Width 30 | 0.065 | 0.006 | 0.025 | -0.001 | 0.954 | 0.985 |
| Lamellar Width 31 | 0.034 | 0.143 | 0.273 | 0.039 | 0.102 | 0.214 |
| Long-axis Length | 0.089 | 0.000 | 0.002 | 0.099 | 0.000 | 0.000 |
| Lamellar Thickness 2 | -0.003 | 0.895 | 0.951 | -0.028 | 0.239 | 0.368 |
| Lamellar Thickness 6 | -0.041 | 0.083 | 0.188 | 0.067 | 0.005 | 0.022 |
| Lamellar Thickness 10 | -0.075 | 0.001 | 0.009 | 0.014 | 0.546 | 0.684 |
| Lamellar Thickness 14 | -0.001 | 0.950 | 0.985 | -0.011 | 0.655 | 0.773 |
| Lamellar Thickness 18 | -0.038 | 0.110 | 0.226 | 0.049 | 0.039 | 0.114 |
| Lamellar Thickness 22 | 0.046 | 0.051 | 0.139 | 0.032 | 0.179 | 0.314 |
| Lamellar Thickness 26 | 0.089 | 0.000 | 0.002 | 0.089 | 0.000 | 0.002 |
| Lamellar Thickness 30 | 0.109 | 0.000 | 0.000 | 0.065 | 0.006 | 0.025 |
| Lamellar Thickness 34 | 0.030 | 0.198 | 0.332 | 0.043 | 0.067 | 0.163 |
| Lamellar Thickness 38 | 0.064 | 0.007 | 0.028 | 0.010 | 0.668 | 0.779 |
| Lamellar Thickness 42 | 0.066 | 0.005 | 0.024 | 0.031 | 0.183 | 0.316 |
| Lamellar Thickness 46 | 0.042 | 0.075 | 0.176 | 0.016 | 0.488 | 0.630 |
| Lamellar Thickness 50 | 0.026 | 0.268 | 0.404 | 0.031 | 0.189 | 0.322 |
| Lamellar Thickness 54 | 0.020 | 0.403 | 0.550 | 0.044 | 0.063 | 0.155 |
| Lamellar Thickness 58 | 0.007 | 0.767 | 0.869 | 0.033 | 0.157 | 0.288 |
| Lamellar Thickness 62 | -0.011 | 0.634 | 0.758 | 0.028 | 0.229 | 0.358 |
| Lamellar Thickness 66 | 0.040 | 0.092 | 0.201 | 0.035 | 0.135 | 0.261 |
| Lamellar Thickness 70 | -0.024 | 0.308 | 0.449 | 0.002 | 0.918 | 0.968 |
| Lamellar Thickness 74 | -0.100 | 0.000 | 0.000 | -0.097 | 0.000 | 0.001 |
| Lamellar Thickness 78 | -0.078 | 0.001 | 0.007 | -0.107 | 0.000 | 0.000 |
| Lamellar Thickness 82 | -0.046 | 0.049 | 0.138 | -0.068 | 0.004 | 0.021 |
| Lamellar Thickness 86 | -0.039 | 0.098 | 0.210 | -0.087 | 0.000 | 0.002 |
| Lamellar Thickness 90 | -0.053 | 0.024 | 0.082 | -0.052 | 0.026 | 0.086 |
| Lamellar Thickness 94 | -0.030 | 0.207 | 0.337 | -0.050 | 0.032 | 0.102 |
| Lamellar Thickness 98 | 0.045 | 0.057 | 0.144 | 0.030 | 0.196 | 0.329 |
| Lamellar Thickness 102 | 0.061 | 0.010 | 0.039 | 0.006 | 0.802 | 0.892 |
| Lamellar Thickness 106 | 0.032 | 0.169 | 0.303 | 0.017 | 0.473 | 0.619 |

| | | | | | | |
|---|---|---|---|---|---|---|
| Lamellar Thickness 110 | 0.049 | 0.039 | 0.114 | 0.053 | 0.024 | 0.082 |
| Lamellar Thickness 109 | 0.019 | 0.413 | 0.558 | 0.089 | 0.000 | 0.002 |
| Lamellar Thickness 105 | 0.017 | 0.462 | 0.607 | 0.066 | 0.005 | 0.023 |
| Lamellar Thickness 101 | -0.001 | 0.951 | 0.985 | 0.034 | 0.153 | 0.288 |
| Lamellar Thickness 97 | 0.013 | 0.592 | 0.726 | 0.049 | 0.038 | 0.114 |
| Lamellar Thickness 93 | -0.018 | 0.443 | 0.588 | -0.029 | 0.212 | 0.341 |
| Lamellar Thickness 89 | -0.025 | 0.283 | 0.422 | -0.044 | 0.063 | 0.155 |
| Lamellar Thickness 85 | -0.023 | 0.326 | 0.461 | -0.047 | 0.045 | 0.132 |
| Lamellar Thickness 81 | -0.036 | 0.122 | 0.245 | -0.053 | 0.025 | 0.084 |
| Lamellar Thickness 77 | -0.054 | 0.023 | 0.081 | -0.080 | 0.001 | 0.005 |
| Lamellar Thickness 73 | -0.049 | 0.036 | 0.112 | -0.057 | 0.016 | 0.058 |
| Lamellar Thickness 69 | 0.000 | 0.983 | 0.990 | -0.039 | 0.100 | 0.213 |
| Lamellar Thickness 65 | 0.030 | 0.203 | 0.334 | -0.025 | 0.289 | 0.429 |
| Lamellar Thickness 61 | 0.078 | 0.001 | 0.007 | 0.029 | 0.213 | 0.341 |
| Lamellar Thickness 57 | 0.094 | 0.000 | 0.001 | 0.052 | 0.028 | 0.092 |
| Lamellar Thickness 53 | 0.090 | 0.000 | 0.002 | 0.094 | 0.000 | 0.001 |
| Lamellar Thickness 49 | 0.077 | 0.001 | 0.007 | 0.100 | 0.000 | 0.000 |
| Lamellar Thickness 45 | 0.070 | 0.003 | 0.016 | 0.111 | 0.000 | 0.000 |
| Lamellar Thickness 41 | 0.077 | 0.001 | 0.007 | 0.141 | 0.000 | 0.000 |
| Lamellar Thickness 37 | 0.088 | 0.000 | 0.002 | 0.106 | 0.000 | 0.000 |
| Lamellar Thickness 33 | 0.053 | 0.024 | 0.082 | 0.068 | 0.004 | 0.020 |
| Lamellar Thickness 29 | 0.041 | 0.084 | 0.190 | 0.018 | 0.447 | 0.590 |
| Lamellar Thickness 25 | 0.019 | 0.416 | 0.560 | -0.005 | 0.824 | 0.906 |
| Lamellar Thickness 21 | -0.004 | 0.874 | 0.943 | -0.006 | 0.786 | 0.881 |
| Lamellar Thickness 17 | -0.071 | 0.003 | 0.015 | 0.004 | 0.873 | 0.943 |
| Lamellar Thickness 13 | -0.056 | 0.018 | 0.065 | -0.039 | 0.096 | 0.207 |
| Lamellar Thickness 9 | -0.060 | 0.010 | 0.041 | -0.053 | 0.023 | 0.081 |
| Lamellar Thickness 5 | -0.100 | 0.000 | 0.000 | -0.024 | 0.308 | 0.449 |
| Lamellar Thickness 1 | 0.126 | 0.000 | 0.000 | 0.076 | 0.001 | 0.008 |
| Lamellar Thickness 4 | 0.057 | 0.015 | 0.057 | 0.031 | 0.183 | 0.316 |
| Lamellar Thickness 8 | 0.063 | 0.007 | 0.030 | 0.042 | 0.073 | 0.171 |
| Lamellar Thickness 12 | 0.034 | 0.154 | 0.288 | -0.001 | 0.963 | 0.985 |

| | | | | | | |
|---|---|---|---|---|---|---|
| Lamellar Thickness 16 | 0.099 | 0.000 | 0.000 | 0.033 | 0.158 | 0.288 |
| Lamellar Thickness 20 | 0.033 | 0.160 | 0.290 | 0.047 | 0.046 | 0.134 |
| Lamellar Thickness 24 | 0.012 | 0.615 | 0.738 | 0.087 | 0.000 | 0.002 |
| Lamellar Thickness 28 | 0.067 | 0.005 | 0.022 | 0.027 | 0.249 | 0.381 |
| Lamellar Thickness 32 | 0.024 | 0.302 | 0.446 | 0.016 | 0.506 | 0.650 |
| Lamellar Thickness 36 | 0.014 | 0.541 | 0.680 | 0.040 | 0.089 | 0.198 |
| Lamellar Thickness 40 | -0.011 | 0.638 | 0.758 | 0.017 | 0.481 | 0.625 |
| Lamellar Thickness 44 | 0.036 | 0.131 | 0.255 | 0.011 | 0.639 | 0.758 |
| Lamellar Thickness 48 | 0.049 | 0.037 | 0.114 | 0.030 | 0.209 | 0.338 |
| Lamellar Thickness 52 | 0.067 | 0.004 | 0.021 | 0.029 | 0.220 | 0.348 |
| Lamellar Thickness 56 | 0.081 | 0.001 | 0.004 | 0.024 | 0.311 | 0.450 |
| Lamellar Thickness 60 | 0.030 | 0.201 | 0.333 | 0.002 | 0.942 | 0.985 |
| Lamellar Thickness 64 | 0.000 | 0.998 | 0.998 | 0.006 | 0.812 | 0.896 |
| Lamellar Thickness 68 | 0.006 | 0.806 | 0.893 | 0.005 | 0.827 | 0.906 |
| Lamellar Thickness 72 | 0.043 | 0.067 | 0.163 | 0.026 | 0.266 | 0.403 |
| Lamellar Thickness 76 | 0.095 | 0.000 | 0.001 | 0.053 | 0.023 | 0.081 |
| Lamellar Thickness 80 | 0.031 | 0.181 | 0.316 | 0.040 | 0.090 | 0.198 |
| Lamellar Thickness 84 | -0.007 | 0.781 | 0.879 | 0.018 | 0.438 | 0.583 |
| Lamellar Thickness 88 | -0.007 | 0.750 | 0.854 | 0.041 | 0.078 | 0.179 |
| Lamellar Thickness 92 | 0.024 | 0.308 | 0.449 | 0.008 | 0.728 | 0.843 |
| Lamellar Thickness 96 | 0.036 | 0.128 | 0.252 | 0.002 | 0.941 | 0.985 |
| Lamellar Thickness 100 | -0.045 | 0.057 | 0.144 | -0.041 | 0.078 | 0.179 |
| Lamellar Thickness 104 | -0.096 | 0.000 | 0.001 | -0.013 | 0.567 | 0.700 |
| Lamellar Thickness 108 | -0.069 | 0.004 | 0.020 | -0.007 | 0.781 | 0.879 |
| Lamellar Thickness 112 | 0.031 | 0.187 | 0.321 | 0.024 | 0.317 | 0.453 |
| Lamellar Thickness 111 | 0.014 | 0.562 | 0.700 | 0.066 | 0.005 | 0.022 |
| Lamellar Thickness 107 | 0.015 | 0.528 | 0.676 | 0.089 | 0.000 | 0.002 |
| Lamellar Thickness 103 | 0.030 | 0.205 | 0.335 | 0.075 | 0.001 | 0.008 |
| Lamellar Thickness 99 | 0.043 | 0.068 | 0.163 | 0.078 | 0.001 | 0.007 |
| Lamellar Thickness 95 | 0.061 | 0.010 | 0.039 | 0.052 | 0.028 | 0.092 |
| Lamellar Thickness 91 | 0.037 | 0.119 | 0.240 | 0.039 | 0.100 | 0.213 |
| Lamellar Thickness 87 | 0.071 | 0.003 | 0.015 | 0.046 | 0.051 | 0.140 |

| | | | | | | |
|---|---|---|---|---|---|---|
| Lamellar Thickness 83 | 0.046 | 0.052 | 0.140 | 0.033 | 0.158 | 0.288 |
| Lamellar Thickness 79 | 0.038 | 0.111 | 0.227 | 0.037 | 0.114 | 0.231 |
| Lamellar Thickness 75 | 0.029 | 0.214 | 0.341 | 0.022 | 0.341 | 0.479 |
| Lamellar Thickness 71 | 0.068 | 0.004 | 0.020 | 0.071 | 0.003 | 0.015 |
| Lamellar Thickness 67 | 0.122 | 0.000 | 0.000 | 0.067 | 0.004 | 0.021 |
| Lamellar Thickness 63 | 0.121 | 0.000 | 0.000 | 0.058 | 0.014 | 0.055 |
| Lamellar Thickness 59 | 0.077 | 0.001 | 0.007 | 0.046 | 0.048 | 0.138 |
| Lamellar Thickness 55 | 0.049 | 0.039 | 0.114 | 0.001 | 0.964 | 0.985 |
| Lamellar Thickness 51 | 0.012 | 0.600 | 0.729 | -0.013 | 0.587 | 0.723 |
| Lamellar Thickness 47 | 0.004 | 0.878 | 0.944 | 0.012 | 0.603 | 0.730 |
| Lamellar Thickness 43 | 0.017 | 0.482 | 0.625 | -0.028 | 0.226 | 0.356 |
| Lamellar Thickness 39 | 0.019 | 0.427 | 0.572 | -0.085 | 0.000 | 0.003 |
| Lamellar Thickness 35 | 0.032 | 0.174 | 0.309 | -0.075 | 0.001 | 0.008 |
| Lamellar Thickness 31 | 0.031 | 0.195 | 0.329 | -0.014 | 0.565 | 0.700 |
| Lamellar Thickness 27 | 0.059 | 0.012 | 0.047 | -0.034 | 0.143 | 0.273 |
| Lamellar Thickness 23 | 0.030 | 0.200 | 0.333 | -0.003 | 0.905 | 0.958 |
| Lamellar Thickness 19 | 0.020 | 0.399 | 0.550 | 0.024 | 0.315 | 0.453 |
| Lamellar Thickness 15 | 0.046 | 0.049 | 0.138 | 0.042 | 0.072 | 0.171 |
| Lamellar Thickness 11 | 0.032 | 0.177 | 0.313 | 0.001 | 0.964 | 0.985 |
| Lamellar Thickness 7 | 0.000 | 0.994 | 0.998 | -0.045 | 0.055 | 0.144 |
| Lamellar Thickness 3 | 0.130 | 0.000 | 0.000 | 0.061 | 0.009 | 0.038 |

#### 21. Supplementary Table S9. Group-wise comparison of normative z-scores between ADHD and test HC

| | Left | | | Right | | |
|---|---|---|---|---|---|---|
| | effect size | P | corrected P | effect size | P | corrected P |
| Lamellar Width 1 | 0.024 | 0.207 | 0.308 | -0.009 | 0.629 | 0.733 |
| Lamellar Width 2 | -0.024 | 0.197 | 0.301 | -0.025 | 0.177 | 0.278 |
| Lamellar Width 3 | 0.006 | 0.767 | 0.821 | -0.037 | 0.050 | 0.109 |
| Lamellar Width 4 | -0.043 | 0.022 | 0.057 | -0.115 | 0.000 | 0.000 |

| | | | | | | |
|---|---|---|---|---|---|---|
| Lamellar Width 5 | -0.052 | 0.005 | 0.018 | -0.106 | 0.000 | 0.000 |
| Lamellar Width 6 | -0.046 | 0.015 | 0.042 | -0.069 | 0.000 | 0.001 |
| Lamellar Width 7 | -0.012 | 0.512 | 0.623 | -0.030 | 0.106 | 0.189 |
| Lamellar Width 8 | -0.039 | 0.039 | 0.087 | -0.047 | 0.013 | 0.038 |
| Lamellar Width 9 | -0.056 | 0.003 | 0.011 | -0.031 | 0.095 | 0.176 |
| Lamellar Width 10 | -0.010 | 0.579 | 0.686 | -0.029 | 0.128 | 0.217 |
| Lamellar Width 11 | 0.027 | 0.148 | 0.241 | -0.029 | 0.127 | 0.217 |
| Lamellar Width 12 | 0.057 | 0.002 | 0.010 | -0.032 | 0.091 | 0.172 |
| Lamellar Width 13 | 0.013 | 0.483 | 0.594 | -0.050 | 0.008 | 0.025 |
| Lamellar Width 14 | -0.007 | 0.695 | 0.770 | -0.054 | 0.004 | 0.014 |
| Lamellar Width 15 | -0.033 | 0.079 | 0.153 | -0.048 | 0.010 | 0.030 |
| Lamellar Width 16 | -0.026 | 0.162 | 0.256 | -0.043 | 0.023 | 0.058 |
| Lamellar Width 17 | -0.029 | 0.124 | 0.214 | -0.027 | 0.149 | 0.242 |
| Lamellar Width 18 | -0.022 | 0.234 | 0.341 | -0.053 | 0.005 | 0.017 |
| Lamellar Width 19 | -0.034 | 0.071 | 0.144 | -0.038 | 0.045 | 0.099 |
| Lamellar Width 20 | -0.023 | 0.220 | 0.323 | -0.058 | 0.002 | 0.009 |
| Lamellar Width 21 | 0.006 | 0.743 | 0.799 | -0.046 | 0.015 | 0.041 |
| Lamellar Width 22 | 0.017 | 0.354 | 0.465 | 0.002 | 0.926 | 0.942 |
| Lamellar Width 23 | 0.014 | 0.457 | 0.567 | 0.039 | 0.037 | 0.084 |
| Lamellar Width 24 | 0.015 | 0.434 | 0.545 | 0.034 | 0.070 | 0.142 |
| Lamellar Width 25 | 0.007 | 0.705 | 0.777 | 0.044 | 0.019 | 0.051 |
| Lamellar Width 26 | -0.007 | 0.714 | 0.779 | 0.003 | 0.873 | 0.915 |
| Lamellar Width 27 | -0.017 | 0.368 | 0.479 | -0.013 | 0.505 | 0.618 |
| Lamellar Width 28 | 0.016 | 0.383 | 0.492 | 0.009 | 0.647 | 0.744 |
| Lamellar Width 29 | 0.024 | 0.201 | 0.302 | 0.009 | 0.636 | 0.738 |
| Lamellar Width 30 | 0.015 | 0.435 | 0.545 | -0.008 | 0.652 | 0.744 |
| Lamellar Width 31 | -0.041 | 0.029 | 0.071 | -0.019 | 0.307 | 0.414 |
| Long-axis Length | 0.044 | 0.020 | 0.053 | 0.032 | 0.095 | 0.176 |
| Lamellar Thickness 2 | 0.009 | 0.651 | 0.744 | -0.043 | 0.021 | 0.055 |
| Lamellar Thickness 6 | 0.010 | 0.608 | 0.711 | 0.056 | 0.003 | 0.011 |
| Lamellar Thickness 10 | -0.047 | 0.013 | 0.036 | 0.007 | 0.709 | 0.779 |
| Lamellar Thickness 14 | 0.021 | 0.262 | 0.367 | -0.001 | 0.945 | 0.958 |

| | | | | | | |
|---|---|---|---|---|---|---|
| Lamellar Thickness 18 | 0.020 | 0.284 | 0.393 | -0.033 | 0.075 | 0.148 |
| Lamellar Thickness 22 | 0.072 | 0.000 | 0.001 | 0.005 | 0.771 | 0.822 |
| Lamellar Thickness 26 | 0.156 | 0.000 | 0.000 | 0.046 | 0.014 | 0.040 |
| Lamellar Thickness 30 | 0.072 | 0.000 | 0.001 | 0.051 | 0.007 | 0.022 |
| Lamellar Thickness 34 | 0.037 | 0.051 | 0.109 | 0.044 | 0.019 | 0.050 |
| Lamellar Thickness 38 | 0.044 | 0.019 | 0.051 | 0.035 | 0.063 | 0.133 |
| Lamellar Thickness 42 | 0.057 | 0.002 | 0.009 | 0.021 | 0.254 | 0.362 |
| Lamellar Thickness 46 | 0.051 | 0.007 | 0.022 | 0.007 | 0.715 | 0.779 |
| Lamellar Thickness 50 | 0.022 | 0.244 | 0.351 | 0.009 | 0.650 | 0.744 |
| Lamellar Thickness 54 | 0.008 | 0.681 | 0.763 | 0.020 | 0.292 | 0.398 |
| Lamellar Thickness 58 | 0.003 | 0.885 | 0.916 | 0.021 | 0.264 | 0.369 |
| Lamellar Thickness 62 | -0.028 | 0.137 | 0.230 | 0.038 | 0.043 | 0.095 |
| Lamellar Thickness 66 | 0.001 | 0.958 | 0.967 | 0.045 | 0.017 | 0.046 |
| Lamellar Thickness 70 | 0.024 | 0.198 | 0.301 | -0.011 | 0.558 | 0.670 |
| Lamellar Thickness 74 | -0.040 | 0.036 | 0.082 | -0.061 | 0.001 | 0.005 |
| Lamellar Thickness 78 | -0.078 | 0.000 | 0.000 | -0.124 | 0.000 | 0.000 |
| Lamellar Thickness 82 | -0.032 | 0.091 | 0.172 | -0.078 | 0.000 | 0.000 |
| Lamellar Thickness 86 | -0.002 | 0.911 | 0.934 | -0.061 | 0.001 | 0.005 |
| Lamellar Thickness 90 | -0.063 | 0.001 | 0.004 | -0.101 | 0.000 | 0.000 |
| Lamellar Thickness 94 | -0.055 | 0.003 | 0.012 | -0.083 | 0.000 | 0.000 |
| Lamellar Thickness 98 | -0.012 | 0.523 | 0.633 | 0.034 | 0.073 | 0.146 |
| Lamellar Thickness 102 | 0.008 | 0.654 | 0.744 | -0.024 | 0.203 | 0.303 |
| Lamellar Thickness 106 | 0.007 | 0.725 | 0.783 | -0.033 | 0.075 | 0.148 |
| Lamellar Thickness 110 | 0.064 | 0.001 | 0.003 | -0.007 | 0.695 | 0.770 |
| Lamellar Thickness 109 | 0.055 | 0.004 | 0.013 | 0.056 | 0.003 | 0.011 |
| Lamellar Thickness 105 | 0.027 | 0.157 | 0.250 | 0.031 | 0.105 | 0.188 |
| Lamellar Thickness 101 | -0.007 | 0.717 | 0.779 | 0.021 | 0.260 | 0.367 |
| Lamellar Thickness 97 | 0.002 | 0.903 | 0.932 | 0.037 | 0.051 | 0.109 |
| Lamellar Thickness 93 | -0.024 | 0.194 | 0.300 | -0.018 | 0.351 | 0.464 |
| Lamellar Thickness 89 | -0.066 | 0.000 | 0.002 | -0.014 | 0.463 | 0.572 |
| Lamellar Thickness 85 | -0.079 | 0.000 | 0.000 | -0.034 | 0.069 | 0.141 |
| Lamellar Thickness 81 | -0.058 | 0.002 | 0.009 | -0.033 | 0.078 | 0.153 |

| | | | | | | |
|---|---|---|---|---|---|---|
| Lamellar Thickness 77 | -0.058 | 0.002 | 0.009 | -0.074 | 0.000 | 0.001 |
| Lamellar Thickness 73 | -0.041 | 0.028 | 0.069 | -0.035 | 0.065 | 0.135 |
| Lamellar Thickness 69 | 0.008 | 0.661 | 0.749 | -0.039 | 0.037 | 0.084 |
| Lamellar Thickness 65 | 0.035 | 0.066 | 0.136 | -0.011 | 0.559 | 0.670 |
| Lamellar Thickness 61 | 0.083 | 0.000 | 0.000 | 0.029 | 0.126 | 0.217 |
| Lamellar Thickness 57 | 0.090 | 0.000 | 0.000 | 0.041 | 0.028 | 0.069 |
| Lamellar Thickness 53 | 0.087 | 0.000 | 0.000 | 0.071 | 0.000 | 0.001 |
| Lamellar Thickness 49 | 0.083 | 0.000 | 0.000 | 0.063 | 0.001 | 0.004 |
| Lamellar Thickness 45 | 0.102 | 0.000 | 0.000 | 0.098 | 0.000 | 0.000 |
| Lamellar Thickness 41 | 0.099 | 0.000 | 0.000 | 0.148 | 0.000 | 0.000 |
| Lamellar Thickness 37 | 0.057 | 0.003 | 0.010 | 0.125 | 0.000 | 0.000 |
| Lamellar Thickness 33 | 0.071 | 0.000 | 0.001 | 0.083 | 0.000 | 0.000 |
| Lamellar Thickness 29 | 0.024 | 0.197 | 0.301 | 0.031 | 0.102 | 0.184 |
| Lamellar Thickness 25 | 0.011 | 0.574 | 0.683 | 0.018 | 0.338 | 0.449 |
| Lamellar Thickness 21 | 0.008 | 0.676 | 0.761 | -0.015 | 0.430 | 0.545 |
| Lamellar Thickness 17 | 0.010 | 0.601 | 0.707 | -0.027 | 0.151 | 0.242 |
| Lamellar Thickness 13 | -0.002 | 0.906 | 0.932 | 0.029 | 0.122 | 0.213 |
| Lamellar Thickness 9 | -0.022 | 0.248 | 0.355 | -0.016 | 0.388 | 0.496 |
| Lamellar Thickness 5 | -0.051 | 0.007 | 0.022 | -0.075 | 0.000 | 0.000 |
| Lamellar Thickness 1 | 0.070 | 0.000 | 0.001 | 0.060 | 0.002 | 0.007 |
| Lamellar Thickness 4 | 0.040 | 0.032 | 0.076 | -0.005 | 0.795 | 0.845 |
| Lamellar Thickness 8 | 0.077 | 0.000 | 0.000 | 0.090 | 0.000 | 0.000 |
| Lamellar Thickness 12 | -0.023 | 0.218 | 0.322 | -0.010 | 0.581 | 0.686 |
| Lamellar Thickness 16 | 0.017 | 0.358 | 0.469 | 0.002 | 0.916 | 0.935 |
| Lamellar Thickness 20 | -0.001 | 0.969 | 0.972 | 0.021 | 0.265 | 0.369 |
| Lamellar Thickness 24 | -0.029 | 0.123 | 0.214 | 0.008 | 0.688 | 0.768 |
| Lamellar Thickness 28 | -0.056 | 0.003 | 0.012 | -0.033 | 0.077 | 0.150 |
| Lamellar Thickness 32 | -0.050 | 0.008 | 0.025 | -0.011 | 0.567 | 0.678 |
| Lamellar Thickness 36 | -0.031 | 0.099 | 0.181 | 0.015 | 0.436 | 0.545 |
| Lamellar Thickness 40 | -0.028 | 0.142 | 0.237 | 0.040 | 0.032 | 0.076 |
| Lamellar Thickness 44 | 0.023 | 0.221 | 0.323 | 0.075 | 0.000 | 0.000 |
| Lamellar Thickness 48 | -0.003 | 0.880 | 0.916 | 0.076 | 0.000 | 0.000 |

| | | | | | | |
|---|---|---|---|---|---|---|
| Lamellar Thickness 52 | 0.030 | 0.113 | 0.198 | 0.055 | 0.003 | 0.012 |
| Lamellar Thickness 56 | 0.040 | 0.033 | 0.076 | 0.073 | 0.000 | 0.001 |
| Lamellar Thickness 60 | 0.058 | 0.002 | 0.008 | 0.041 | 0.029 | 0.071 |
| Lamellar Thickness 64 | -0.001 | 0.973 | 0.973 | 0.003 | 0.863 | 0.909 |
| Lamellar Thickness 68 | -0.019 | 0.301 | 0.407 | -0.024 | 0.200 | 0.301 |
| Lamellar Thickness 72 | -0.024 | 0.199 | 0.301 | -0.022 | 0.236 | 0.342 |
| Lamellar Thickness 76 | 0.027 | 0.144 | 0.238 | 0.035 | 0.065 | 0.135 |
| Lamellar Thickness 80 | 0.045 | 0.018 | 0.048 | -0.003 | 0.882 | 0.916 |
| Lamellar Thickness 84 | 0.007 | 0.720 | 0.779 | -0.040 | 0.032 | 0.076 |
| Lamellar Thickness 88 | 0.020 | 0.286 | 0.394 | 0.033 | 0.084 | 0.161 |
| Lamellar Thickness 92 | 0.025 | 0.185 | 0.288 | 0.040 | 0.034 | 0.078 |
| Lamellar Thickness 96 | 0.048 | 0.010 | 0.030 | 0.027 | 0.145 | 0.238 |
| Lamellar Thickness 100 | -0.019 | 0.316 | 0.423 | -0.020 | 0.294 | 0.399 |
| Lamellar Thickness 104 | -0.060 | 0.001 | 0.006 | -0.030 | 0.111 | 0.197 |
| Lamellar Thickness 108 | -0.053 | 0.005 | 0.017 | -0.008 | 0.672 | 0.759 |
| Lamellar Thickness 112 | 0.027 | 0.151 | 0.242 | -0.016 | 0.381 | 0.492 |
| Lamellar Thickness 111 | 0.042 | 0.026 | 0.065 | 0.047 | 0.012 | 0.034 |
| Lamellar Thickness 107 | 0.027 | 0.149 | 0.242 | 0.028 | 0.135 | 0.227 |
| Lamellar Thickness 103 | 0.015 | 0.415 | 0.529 | 0.071 | 0.000 | 0.001 |
| Lamellar Thickness 99 | 0.032 | 0.087 | 0.166 | 0.085 | 0.000 | 0.000 |
| Lamellar Thickness 95 | 0.030 | 0.106 | 0.189 | 0.067 | 0.000 | 0.002 |
| Lamellar Thickness 91 | 0.073 | 0.000 | 0.001 | 0.094 | 0.000 | 0.000 |
| Lamellar Thickness 87 | 0.098 | 0.000 | 0.000 | 0.077 | 0.000 | 0.000 |
| Lamellar Thickness 83 | 0.081 | 0.000 | 0.000 | 0.065 | 0.001 | 0.003 |
| Lamellar Thickness 79 | 0.082 | 0.000 | 0.000 | 0.062 | 0.001 | 0.005 |
| Lamellar Thickness 75 | 0.069 | 0.000 | 0.001 | 0.049 | 0.009 | 0.028 |
| Lamellar Thickness 71 | 0.078 | 0.000 | 0.000 | 0.073 | 0.000 | 0.001 |
| Lamellar Thickness 67 | 0.112 | 0.000 | 0.000 | 0.096 | 0.000 | 0.000 |
| Lamellar Thickness 63 | 0.117 | 0.000 | 0.000 | 0.138 | 0.000 | 0.000 |
| Lamellar Thickness 59 | 0.096 | 0.000 | 0.000 | 0.128 | 0.000 | 0.000 |
| Lamellar Thickness 55 | 0.078 | 0.000 | 0.000 | 0.066 | 0.000 | 0.002 |
| Lamellar Thickness 51 | 0.043 | 0.024 | 0.060 | 0.019 | 0.321 | 0.428 |

| | | | | | | |
|---|---|---|---|---|---|---|
| Lamellar Thickness 47 | -0.021 | 0.258 | 0.366 | 0.038 | 0.045 | 0.099 |
| Lamellar Thickness 43 | -0.031 | 0.101 | 0.184 | 0.004 | 0.812 | 0.860 |
| Lamellar Thickness 39 | -0.015 | 0.438 | 0.546 | -0.050 | 0.008 | 0.025 |
| Lamellar Thickness 35 | 0.001 | 0.960 | 0.967 | -0.025 | 0.177 | 0.278 |
| Lamellar Thickness 31 | 0.052 | 0.006 | 0.018 | 0.016 | 0.383 | 0.492 |
| Lamellar Thickness 27 | 0.054 | 0.004 | 0.014 | 0.013 | 0.506 | 0.618 |
| Lamellar Thickness 23 | 0.070 | 0.000 | 0.001 | 0.035 | 0.061 | 0.128 |
| Lamellar Thickness 19 | 0.082 | 0.000 | 0.000 | 0.031 | 0.095 | 0.176 |
| Lamellar Thickness 15 | 0.096 | 0.000 | 0.000 | 0.052 | 0.006 | 0.020 |
| Lamellar Thickness 11 | 0.078 | 0.000 | 0.000 | 0.020 | 0.290 | 0.398 |
| Lamellar Thickness 7 | 0.038 | 0.041 | 0.092 | -0.003 | 0.865 | 0.909 |
| Lamellar Thickness 3 | 0.086 | 0.000 | 0.000 | 0.055 | 0.003 | 0.012 |

#### 22. Supplementary Table S10. Group-wise comparison of normative z-scores between ANX and test HC

| | Left | | | Right | | |
|---|---|---|---|---|---|---|
| | effect size | P | corrected P | effect size | P | corrected P |
| Lamellar Width 1 | 0.090 | 0.008 | 0.029 | 0.079 | 0.020 | 0.061 |
| Lamellar Width 2 | -0.001 | 0.983 | 0.990 | 0.033 | 0.325 | 0.459 |
| Lamellar Width 3 | 0.059 | 0.084 | 0.169 | 0.044 | 0.198 | 0.328 |
| Lamellar Width 4 | -0.045 | 0.189 | 0.316 | -0.060 | 0.077 | 0.160 |
| Lamellar Width 5 | -0.052 | 0.122 | 0.223 | -0.085 | 0.012 | 0.041 |
| Lamellar Width 6 | -0.001 | 0.983 | 0.990 | -0.078 | 0.021 | 0.062 |
| Lamellar Width 7 | 0.015 | 0.656 | 0.747 | 0.033 | 0.325 | 0.459 |
| Lamellar Width 8 | 0.014 | 0.679 | 0.761 | 0.035 | 0.298 | 0.435 |
| Lamellar Width 9 | 0.015 | 0.661 | 0.748 | 0.113 | 0.001 | 0.005 |
| Lamellar Width 10 | 0.080 | 0.018 | 0.057 | 0.112 | 0.001 | 0.006 |
| Lamellar Width 11 | 0.096 | 0.004 | 0.019 | 0.109 | 0.001 | 0.007 |
| Lamellar Width 12 | 0.142 | 0.000 | 0.000 | 0.111 | 0.001 | 0.006 |
| Lamellar Width 13 | 0.093 | 0.006 | 0.023 | 0.063 | 0.064 | 0.140 |

| | | | | | | |
|---|---|---|---|---|---|---|
| Lamellar Width 14 | 0.095 | 0.005 | 0.021 | 0.020 | 0.564 | 0.677 |
| Lamellar Width 15 | 0.057 | 0.093 | 0.183 | 0.025 | 0.455 | 0.595 |
| Lamellar Width 16 | 0.063 | 0.064 | 0.140 | 0.026 | 0.446 | 0.586 |
| Lamellar Width 17 | 0.043 | 0.201 | 0.329 | 0.040 | 0.243 | 0.380 |
| Lamellar Width 18 | 0.053 | 0.118 | 0.221 | 0.039 | 0.251 | 0.386 |
| Lamellar Width 19 | 0.011 | 0.736 | 0.806 | 0.021 | 0.542 | 0.658 |
| Lamellar Width 20 | 0.023 | 0.491 | 0.618 | -0.012 | 0.718 | 0.793 |
| Lamellar Width 21 | 0.070 | 0.038 | 0.093 | 0.015 | 0.654 | 0.747 |
| Lamellar Width 22 | 0.115 | 0.001 | 0.004 | 0.087 | 0.010 | 0.036 |
| Lamellar Width 23 | 0.125 | 0.000 | 0.002 | 0.127 | 0.000 | 0.001 |
| Lamellar Width 24 | 0.152 | 0.000 | 0.000 | 0.149 | 0.000 | 0.000 |
| Lamellar Width 25 | 0.137 | 0.000 | 0.001 | 0.150 | 0.000 | 0.000 |
| Lamellar Width 26 | 0.104 | 0.002 | 0.011 | 0.068 | 0.044 | 0.103 |
| Lamellar Width 27 | 0.101 | 0.003 | 0.013 | 0.047 | 0.162 | 0.281 |
| Lamellar Width 28 | 0.119 | 0.000 | 0.003 | 0.044 | 0.192 | 0.319 |
| Lamellar Width 29 | 0.091 | 0.007 | 0.028 | 0.070 | 0.038 | 0.094 |
| Lamellar Width 30 | 0.097 | 0.004 | 0.018 | 0.033 | 0.332 | 0.467 |
| Lamellar Width 31 | -0.002 | 0.953 | 0.966 | -0.037 | 0.276 | 0.412 |
| Long-axis Length | 0.106 | 0.002 | 0.009 | 0.047 | 0.166 | 0.284 |
| Lamellar Thickness 2 | 0.040 | 0.234 | 0.368 | 0.035 | 0.301 | 0.435 |
| Lamellar Thickness 6 | 0.126 | 0.000 | 0.001 | 0.085 | 0.012 | 0.041 |
| Lamellar Thickness 10 | -0.018 | 0.589 | 0.698 | 0.071 | 0.036 | 0.091 |
| Lamellar Thickness 14 | 0.037 | 0.276 | 0.412 | 0.021 | 0.533 | 0.652 |
| Lamellar Thickness 18 | -0.024 | 0.476 | 0.609 | 0.000 | 0.991 | 0.993 |
| Lamellar Thickness 22 | 0.086 | 0.011 | 0.039 | 0.115 | 0.001 | 0.005 |
| Lamellar Thickness 26 | 0.149 | 0.000 | 0.000 | 0.094 | 0.006 | 0.023 |
| Lamellar Thickness 30 | 0.059 | 0.079 | 0.164 | 0.046 | 0.174 | 0.296 |
| Lamellar Thickness 34 | 0.033 | 0.323 | 0.459 | 0.114 | 0.001 | 0.005 |
| Lamellar Thickness 38 | 0.077 | 0.023 | 0.067 | 0.121 | 0.000 | 0.003 |
| Lamellar Thickness 42 | 0.144 | 0.000 | 0.000 | 0.053 | 0.122 | 0.223 |
| Lamellar Thickness 46 | 0.105 | 0.002 | 0.010 | 0.000 | 0.993 | 0.993 |
| Lamellar Thickness 50 | 0.080 | 0.019 | 0.060 | 0.016 | 0.637 | 0.737 |

| | | | | | | |
|---|---|---|---|---|---|---|
| Lamellar Thickness 54 | 0.079 | 0.020 | 0.062 | 0.019 | 0.585 | 0.696 |
| Lamellar Thickness 58 | 0.076 | 0.025 | 0.072 | 0.037 | 0.270 | 0.411 |
| Lamellar Thickness 62 | 0.070 | 0.039 | 0.094 | 0.075 | 0.027 | 0.074 |
| Lamellar Thickness 66 | 0.071 | 0.035 | 0.089 | 0.084 | 0.013 | 0.044 |
| Lamellar Thickness 70 | 0.045 | 0.185 | 0.312 | 0.060 | 0.075 | 0.157 |
| Lamellar Thickness 74 | 0.020 | 0.547 | 0.662 | -0.036 | 0.292 | 0.431 |
| Lamellar Thickness 78 | -0.028 | 0.404 | 0.546 | -0.154 | 0.000 | 0.000 |
| Lamellar Thickness 82 | 0.005 | 0.883 | 0.928 | -0.073 | 0.031 | 0.080 |
| Lamellar Thickness 86 | -0.022 | 0.512 | 0.632 | -0.064 | 0.060 | 0.134 |
| Lamellar Thickness 90 | -0.057 | 0.094 | 0.183 | -0.047 | 0.163 | 0.282 |
| Lamellar Thickness 94 | -0.059 | 0.083 | 0.169 | -0.036 | 0.287 | 0.426 |
| Lamellar Thickness 98 | -0.031 | 0.365 | 0.497 | 0.039 | 0.251 | 0.386 |
| Lamellar Thickness 102 | 0.004 | 0.910 | 0.939 | -0.035 | 0.299 | 0.435 |
| Lamellar Thickness 106 | 0.013 | 0.700 | 0.775 | -0.074 | 0.029 | 0.079 |
| Lamellar Thickness 110 | 0.049 | 0.148 | 0.261 | -0.050 | 0.143 | 0.254 |
| Lamellar Thickness 109 | 0.026 | 0.435 | 0.580 | 0.023 | 0.500 | 0.622 |
| Lamellar Thickness 105 | -0.040 | 0.232 | 0.368 | -0.017 | 0.622 | 0.725 |
| Lamellar Thickness 101 | -0.033 | 0.334 | 0.467 | 0.010 | 0.768 | 0.828 |
| Lamellar Thickness 97 | 0.003 | 0.939 | 0.959 | 0.064 | 0.060 | 0.134 |
| Lamellar Thickness 93 | -0.026 | 0.439 | 0.580 | 0.024 | 0.475 | 0.609 |
| Lamellar Thickness 89 | -0.058 | 0.085 | 0.169 | 0.012 | 0.729 | 0.801 |
| Lamellar Thickness 85 | -0.051 | 0.132 | 0.238 | -0.057 | 0.095 | 0.183 |
| Lamellar Thickness 81 | -0.007 | 0.826 | 0.874 | -0.059 | 0.081 | 0.167 |
| Lamellar Thickness 77 | -0.020 | 0.556 | 0.670 | -0.101 | 0.003 | 0.014 |
| Lamellar Thickness 73 | -0.009 | 0.779 | 0.838 | -0.008 | 0.812 | 0.866 |
| Lamellar Thickness 69 | 0.094 | 0.006 | 0.023 | 0.035 | 0.296 | 0.435 |
| Lamellar Thickness 65 | 0.149 | 0.000 | 0.000 | 0.073 | 0.031 | 0.080 |
| Lamellar Thickness 61 | 0.164 | 0.000 | 0.000 | 0.110 | 0.001 | 0.007 |
| Lamellar Thickness 57 | 0.138 | 0.000 | 0.001 | 0.064 | 0.060 | 0.134 |
| Lamellar Thickness 53 | 0.142 | 0.000 | 0.000 | 0.075 | 0.026 | 0.074 |
| Lamellar Thickness 49 | 0.158 | 0.000 | 0.000 | 0.085 | 0.012 | 0.041 |
| Lamellar Thickness 45 | 0.184 | 0.000 | 0.000 | 0.089 | 0.009 | 0.031 |

| | | | | | | |
|---|---|---|---|---|---|---|
| Lamellar Thickness 41 | 0.172 | 0.000 | 0.000 | 0.153 | 0.000 | 0.000 |
| Lamellar Thickness 37 | 0.137 | 0.000 | 0.001 | 0.091 | 0.007 | 0.028 |
| Lamellar Thickness 33 | 0.075 | 0.027 | 0.074 | 0.025 | 0.468 | 0.607 |
| Lamellar Thickness 29 | 0.073 | 0.032 | 0.082 | -0.019 | 0.579 | 0.692 |
| Lamellar Thickness 25 | 0.056 | 0.099 | 0.187 | -0.004 | 0.895 | 0.934 |
| Lamellar Thickness 21 | 0.043 | 0.201 | 0.329 | 0.016 | 0.627 | 0.729 |
| Lamellar Thickness 17 | 0.043 | 0.206 | 0.335 | -0.011 | 0.750 | 0.815 |
| Lamellar Thickness 13 | 0.014 | 0.673 | 0.757 | 0.023 | 0.494 | 0.619 |
| Lamellar Thickness 9 | -0.034 | 0.313 | 0.451 | -0.017 | 0.606 | 0.713 |
| Lamellar Thickness 5 | -0.081 | 0.017 | 0.056 | -0.061 | 0.075 | 0.157 |
| Lamellar Thickness 1 | 0.062 | 0.067 | 0.144 | 0.053 | 0.120 | 0.223 |
| Lamellar Thickness 4 | 0.073 | 0.031 | 0.080 | 0.069 | 0.041 | 0.098 |
| Lamellar Thickness 8 | 0.079 | 0.020 | 0.061 | 0.067 | 0.049 | 0.113 |
| Lamellar Thickness 12 | -0.050 | 0.138 | 0.248 | -0.032 | 0.344 | 0.475 |
| Lamellar Thickness 16 | 0.018 | 0.603 | 0.711 | 0.024 | 0.480 | 0.612 |
| Lamellar Thickness 20 | 0.032 | 0.345 | 0.475 | 0.041 | 0.233 | 0.368 |
| Lamellar Thickness 24 | -0.013 | 0.699 | 0.775 | -0.024 | 0.487 | 0.618 |
| Lamellar Thickness 28 | -0.110 | 0.001 | 0.006 | -0.042 | 0.220 | 0.353 |
| Lamellar Thickness 32 | -0.138 | 0.000 | 0.001 | -0.051 | 0.133 | 0.239 |
| Lamellar Thickness 36 | -0.101 | 0.003 | 0.014 | -0.037 | 0.275 | 0.412 |
| Lamellar Thickness 40 | -0.125 | 0.000 | 0.002 | -0.015 | 0.662 | 0.748 |
| Lamellar Thickness 44 | -0.002 | 0.948 | 0.965 | 0.073 | 0.032 | 0.082 |
| Lamellar Thickness 48 | 0.007 | 0.826 | 0.874 | 0.056 | 0.098 | 0.187 |
| Lamellar Thickness 52 | 0.031 | 0.366 | 0.497 | 0.059 | 0.082 | 0.167 |
| Lamellar Thickness 56 | 0.069 | 0.043 | 0.102 | 0.068 | 0.045 | 0.105 |
| Lamellar Thickness 60 | 0.113 | 0.001 | 0.005 | 0.076 | 0.025 | 0.072 |
| Lamellar Thickness 64 | 0.048 | 0.154 | 0.270 | 0.069 | 0.044 | 0.103 |
| Lamellar Thickness 68 | 0.015 | 0.648 | 0.744 | 0.050 | 0.143 | 0.254 |
| Lamellar Thickness 72 | 0.042 | 0.213 | 0.344 | 0.013 | 0.694 | 0.774 |
| Lamellar Thickness 76 | 0.024 | 0.471 | 0.608 | 0.137 | 0.000 | 0.001 |
| Lamellar Thickness 80 | 0.017 | 0.612 | 0.717 | 0.101 | 0.003 | 0.013 |
| Lamellar Thickness 84 | 0.004 | 0.895 | 0.934 | 0.023 | 0.501 | 0.622 |

| | | | | | | |
|---|---|---|---|---|---|---|
| Lamellar Thickness 88 | 0.023 | 0.491 | 0.618 | 0.139 | 0.000 | 0.000 |
| Lamellar Thickness 92 | 0.038 | 0.260 | 0.399 | 0.161 | 0.000 | 0.000 |
| Lamellar Thickness 96 | 0.120 | 0.000 | 0.003 | 0.158 | 0.000 | 0.000 |
| Lamellar Thickness 100 | -0.008 | 0.812 | 0.866 | 0.011 | 0.740 | 0.808 |
| Lamellar Thickness 104 | -0.056 | 0.098 | 0.187 | 0.006 | 0.863 | 0.911 |
| Lamellar Thickness 108 | -0.081 | 0.016 | 0.054 | -0.003 | 0.920 | 0.943 |
| Lamellar Thickness 112 | 0.051 | 0.131 | 0.238 | -0.061 | 0.073 | 0.155 |
| Lamellar Thickness 111 | 0.021 | 0.531 | 0.652 | 0.026 | 0.439 | 0.580 |
| Lamellar Thickness 107 | -0.042 | 0.216 | 0.347 | -0.032 | 0.350 | 0.481 |
| Lamellar Thickness 103 | 0.021 | 0.534 | 0.652 | 0.010 | 0.766 | 0.828 |
| Lamellar Thickness 99 | 0.045 | 0.184 | 0.311 | 0.068 | 0.047 | 0.108 |
| Lamellar Thickness 95 | 0.093 | 0.006 | 0.023 | 0.077 | 0.023 | 0.068 |
| Lamellar Thickness 91 | 0.155 | 0.000 | 0.000 | 0.215 | 0.000 | 0.000 |
| Lamellar Thickness 87 | 0.160 | 0.000 | 0.000 | 0.127 | 0.000 | 0.001 |
| Lamellar Thickness 83 | 0.100 | 0.003 | 0.014 | 0.133 | 0.000 | 0.001 |
| Lamellar Thickness 79 | 0.130 | 0.000 | 0.001 | 0.104 | 0.002 | 0.011 |
| Lamellar Thickness 75 | 0.090 | 0.008 | 0.028 | 0.092 | 0.006 | 0.025 |
| Lamellar Thickness 71 | 0.074 | 0.030 | 0.080 | 0.110 | 0.001 | 0.007 |
| Lamellar Thickness 67 | 0.151 | 0.000 | 0.000 | 0.143 | 0.000 | 0.000 |
| Lamellar Thickness 63 | 0.170 | 0.000 | 0.000 | 0.134 | 0.000 | 0.001 |
| Lamellar Thickness 59 | 0.132 | 0.000 | 0.001 | 0.115 | 0.001 | 0.005 |
| Lamellar Thickness 55 | 0.075 | 0.027 | 0.074 | 0.063 | 0.065 | 0.140 |
| Lamellar Thickness 51 | 0.071 | 0.037 | 0.092 | 0.025 | 0.462 | 0.602 |
| Lamellar Thickness 47 | -0.033 | 0.325 | 0.459 | 0.003 | 0.919 | 0.943 |
| Lamellar Thickness 43 | -0.077 | 0.023 | 0.068 | -0.079 | 0.020 | 0.061 |
| Lamellar Thickness 39 | -0.079 | 0.019 | 0.061 | -0.159 | 0.000 | 0.000 |
| Lamellar Thickness 35 | -0.058 | 0.085 | 0.169 | -0.103 | 0.002 | 0.011 |
| Lamellar Thickness 31 | -0.037 | 0.273 | 0.412 | -0.091 | 0.007 | 0.028 |
| Lamellar Thickness 27 | 0.027 | 0.434 | 0.580 | -0.040 | 0.245 | 0.381 |
| Lamellar Thickness 23 | 0.057 | 0.093 | 0.183 | 0.027 | 0.420 | 0.565 |
| Lamellar Thickness 19 | 0.156 | 0.000 | 0.000 | -0.004 | 0.902 | 0.937 |
| Lamellar Thickness 15 | 0.113 | 0.001 | 0.005 | 0.032 | 0.338 | 0.471 |

| | | | | | | |
|---|---|---|---|---|---|---|
| Lamellar Thickness 11 | 0.097 | 0.004 | 0.019 | -0.004 | 0.904 | 0.937 |
| Lamellar Thickness 7 | 0.063 | 0.062 | 0.137 | 0.016 | 0.648 | 0.744 |
| Lamellar Thickness 3 | 0.070 | 0.039 | 0.095 | 0.056 | 0.099 | 0.187 |

### 23. Supplementary Table S11. Group-wise comparison of normative z-scores between MDD and test HC

| | Left | | | Right | | |
|---|---|---|---|---|---|---|
| | effect size | P | corrected P | effect size | P | corrected P |
| Lamellar Width 1 | 0.026 | 0.370 | 0.773 | 0.065 | 0.027 | 0.242 |
| Lamellar Width 2 | -0.016 | 0.580 | 0.860 | 0.087 | 0.003 | 0.093 |
| Lamellar Width 3 | -0.024 | 0.422 | 0.795 | 0.051 | 0.083 | 0.435 |
| Lamellar Width 4 | -0.020 | 0.487 | 0.830 | -0.017 | 0.552 | 0.860 |
| Lamellar Width 5 | -0.011 | 0.705 | 0.887 | -0.078 | 0.008 | 0.154 |
| Lamellar Width 6 | -0.031 | 0.288 | 0.672 | -0.031 | 0.294 | 0.672 |
| Lamellar Width 7 | -0.033 | 0.262 | 0.645 | -0.033 | 0.267 | 0.645 |
| Lamellar Width 8 | -0.068 | 0.020 | 0.207 | -0.068 | 0.020 | 0.207 |
| Lamellar Width 9 | -0.046 | 0.120 | 0.493 | -0.035 | 0.230 | 0.645 |
| Lamellar Width 10 | -0.011 | 0.697 | 0.887 | -0.037 | 0.206 | 0.645 |
| Lamellar Width 11 | 0.023 | 0.443 | 0.813 | -0.026 | 0.367 | 0.773 |
| Lamellar Width 12 | 0.018 | 0.550 | 0.860 | 0.004 | 0.897 | 0.957 |
| Lamellar Width 13 | -0.007 | 0.804 | 0.922 | -0.002 | 0.950 | 0.974 |
| Lamellar Width 14 | -0.034 | 0.241 | 0.645 | -0.028 | 0.332 | 0.724 |
| Lamellar Width 15 | -0.014 | 0.636 | 0.866 | -0.025 | 0.385 | 0.775 |
| Lamellar Width 16 | -0.048 | 0.105 | 0.460 | -0.048 | 0.104 | 0.460 |
| Lamellar Width 17 | -0.052 | 0.078 | 0.419 | -0.015 | 0.608 | 0.863 |
| Lamellar Width 18 | -0.026 | 0.369 | 0.773 | 0.004 | 0.883 | 0.954 |
| Lamellar Width 19 | -0.014 | 0.644 | 0.866 | -0.005 | 0.856 | 0.948 |
| Lamellar Width 20 | -0.069 | 0.019 | 0.207 | -0.020 | 0.495 | 0.833 |
| Lamellar Width 21 | -0.052 | 0.078 | 0.419 | -0.001 | 0.960 | 0.976 |
| Lamellar Width 22 | -0.021 | 0.486 | 0.830 | 0.032 | 0.276 | 0.656 |
| Lamellar Width 23 | -0.025 | 0.401 | 0.784 | 0.024 | 0.416 | 0.792 |

| | | | | | | |
|---|---|---|---|---|---|---|
| Lamellar Width 24 | -0.058 | 0.048 | 0.312 | 0.005 | 0.855 | 0.948 |
| Lamellar Width 25 | -0.085 | 0.004 | 0.101 | -0.009 | 0.767 | 0.902 |
| Lamellar Width 26 | -0.078 | 0.008 | 0.154 | 0.000 | 0.998 | 1.000 |
| Lamellar Width 27 | -0.060 | 0.041 | 0.291 | -0.016 | 0.593 | 0.860 |
| Lamellar Width 28 | -0.018 | 0.542 | 0.860 | 0.016 | 0.597 | 0.860 |
| Lamellar Width 29 | 0.022 | 0.457 | 0.820 | 0.010 | 0.726 | 0.902 |
| Lamellar Width 30 | 0.046 | 0.121 | 0.493 | 0.004 | 0.884 | 0.954 |
| Lamellar Width 31 | 0.032 | 0.272 | 0.653 | 0.076 | 0.010 | 0.169 |
| Long-axis Length | 0.065 | 0.028 | 0.243 | 0.034 | 0.250 | 0.645 |
| Lamellar Thickness 2 | -0.012 | 0.682 | 0.880 | 0.024 | 0.418 | 0.792 |
| Lamellar Thickness 6 | 0.001 | 0.963 | 0.977 | 0.023 | 0.441 | 0.813 |
| Lamellar Thickness 10 | -0.009 | 0.750 | 0.902 | -0.006 | 0.842 | 0.945 |
| Lamellar Thickness 14 | -0.015 | 0.612 | 0.864 | -0.025 | 0.403 | 0.784 |
| Lamellar Thickness 18 | -0.002 | 0.933 | 0.974 | 0.058 | 0.047 | 0.312 |
| Lamellar Thickness 22 | 0.061 | 0.039 | 0.282 | 0.100 | 0.001 | 0.046 |
| Lamellar Thickness 26 | 0.013 | 0.649 | 0.866 | 0.035 | 0.236 | 0.645 |
| Lamellar Thickness 30 | 0.043 | 0.145 | 0.521 | 0.059 | 0.045 | 0.310 |
| Lamellar Thickness 34 | -0.017 | 0.559 | 0.860 | 0.070 | 0.017 | 0.200 |
| Lamellar Thickness 38 | -0.013 | 0.652 | 0.866 | 0.045 | 0.124 | 0.493 |
| Lamellar Thickness 42 | 0.022 | 0.450 | 0.816 | -0.009 | 0.769 | 0.902 |
| Lamellar Thickness 46 | -0.005 | 0.872 | 0.951 | -0.013 | 0.666 | 0.866 |
| Lamellar Thickness 50 | -0.034 | 0.249 | 0.645 | -0.026 | 0.380 | 0.775 |
| Lamellar Thickness 54 | 0.013 | 0.668 | 0.866 | -0.017 | 0.572 | 0.860 |
| Lamellar Thickness 58 | 0.005 | 0.868 | 0.951 | -0.019 | 0.507 | 0.839 |
| Lamellar Thickness 62 | 0.015 | 0.605 | 0.862 | -0.052 | 0.079 | 0.419 |
| Lamellar Thickness 66 | 0.042 | 0.153 | 0.531 | 0.010 | 0.743 | 0.902 |
| Lamellar Thickness 70 | 0.017 | 0.561 | 0.860 | -0.019 | 0.506 | 0.839 |
| Lamellar Thickness 74 | -0.016 | 0.576 | 0.860 | -0.050 | 0.088 | 0.452 |
| Lamellar Thickness 78 | -0.021 | 0.471 | 0.825 | -0.056 | 0.058 | 0.353 |
| Lamellar Thickness 82 | -0.045 | 0.127 | 0.493 | -0.043 | 0.143 | 0.520 |
| Lamellar Thickness 86 | -0.048 | 0.102 | 0.460 | -0.048 | 0.100 | 0.460 |
| Lamellar Thickness 90 | -0.036 | 0.216 | 0.645 | -0.071 | 0.015 | 0.185 |

| | | | | | | |
|---|---|---|---|---|---|---|
| Lamellar Thickness 94 | -0.024 | 0.416 | 0.792 | -0.098 | 0.001 | 0.046 |
| Lamellar Thickness 98 | -0.002 | 0.945 | 0.974 | 0.017 | 0.560 | 0.860 |
| Lamellar Thickness 102 | -0.037 | 0.212 | 0.645 | -0.009 | 0.771 | 0.902 |
| Lamellar Thickness 106 | 0.029 | 0.331 | 0.724 | 0.003 | 0.916 | 0.969 |
| Lamellar Thickness 110 | -0.002 | 0.934 | 0.974 | 0.021 | 0.466 | 0.823 |
| Lamellar Thickness 109 | 0.022 | 0.459 | 0.820 | 0.036 | 0.216 | 0.645 |
| Lamellar Thickness 105 | -0.010 | 0.736 | 0.902 | 0.030 | 0.305 | 0.683 |
| Lamellar Thickness 101 | -0.027 | 0.357 | 0.773 | 0.017 | 0.567 | 0.860 |
| Lamellar Thickness 97 | 0.002 | 0.959 | 0.976 | 0.036 | 0.221 | 0.645 |
| Lamellar Thickness 93 | -0.033 | 0.258 | 0.645 | -0.036 | 0.219 | 0.645 |
| Lamellar Thickness 89 | -0.035 | 0.239 | 0.645 | -0.045 | 0.127 | 0.493 |
| Lamellar Thickness 85 | -0.043 | 0.148 | 0.521 | -0.061 | 0.039 | 0.282 |
| Lamellar Thickness 81 | -0.040 | 0.175 | 0.587 | -0.049 | 0.096 | 0.454 |
| Lamellar Thickness 77 | -0.072 | 0.014 | 0.185 | -0.095 | 0.001 | 0.051 |
| Lamellar Thickness 73 | -0.037 | 0.211 | 0.645 | -0.112 | 0.000 | 0.020 |
| Lamellar Thickness 69 | -0.031 | 0.296 | 0.672 | -0.125 | 0.000 | 0.006 |
| Lamellar Thickness 65 | 0.006 | 0.843 | 0.945 | -0.048 | 0.101 | 0.460 |
| Lamellar Thickness 61 | 0.033 | 0.258 | 0.645 | -0.004 | 0.893 | 0.957 |
| Lamellar Thickness 57 | 0.033 | 0.266 | 0.645 | -0.054 | 0.065 | 0.377 |
| Lamellar Thickness 53 | 0.045 | 0.124 | 0.493 | 0.016 | 0.579 | 0.860 |
| Lamellar Thickness 49 | 0.049 | 0.096 | 0.454 | -0.021 | 0.478 | 0.825 |
| Lamellar Thickness 45 | 0.044 | 0.138 | 0.516 | 0.007 | 0.810 | 0.922 |
| Lamellar Thickness 41 | 0.031 | 0.296 | 0.672 | 0.013 | 0.653 | 0.866 |
| Lamellar Thickness 37 | 0.034 | 0.253 | 0.645 | 0.015 | 0.599 | 0.860 |
| Lamellar Thickness 33 | 0.004 | 0.897 | 0.957 | -0.019 | 0.517 | 0.846 |
| Lamellar Thickness 29 | 0.022 | 0.462 | 0.822 | -0.013 | 0.658 | 0.866 |
| Lamellar Thickness 25 | 0.050 | 0.091 | 0.452 | 0.013 | 0.657 | 0.866 |
| Lamellar Thickness 21 | 0.020 | 0.491 | 0.832 | 0.026 | 0.375 | 0.775 |
| Lamellar Thickness 17 | 0.014 | 0.635 | 0.866 | 0.052 | 0.076 | 0.419 |
| Lamellar Thickness 13 | -0.014 | 0.623 | 0.866 | 0.071 | 0.015 | 0.185 |
| Lamellar Thickness 9 | -0.012 | 0.694 | 0.887 | 0.013 | 0.655 | 0.866 |
| Lamellar Thickness 5 | 0.005 | 0.852 | 0.948 | -0.031 | 0.287 | 0.672 |

| | | | | | | |
|---|---|---|---|---|---|---|
| Lamellar Thickness 1 | 0.068 | 0.021 | 0.207 | -0.049 | 0.093 | 0.453 |
| Lamellar Thickness 4 | -0.029 | 0.325 | 0.720 | 0.024 | 0.407 | 0.787 |
| Lamellar Thickness 8 | -0.043 | 0.148 | 0.521 | -0.007 | 0.808 | 0.922 |
| Lamellar Thickness 12 | 0.009 | 0.754 | 0.902 | 0.010 | 0.739 | 0.902 |
| Lamellar Thickness 16 | 0.009 | 0.767 | 0.902 | -0.025 | 0.397 | 0.783 |
| Lamellar Thickness 20 | -0.013 | 0.654 | 0.866 | -0.033 | 0.264 | 0.645 |
| Lamellar Thickness 24 | -0.016 | 0.588 | 0.860 | 0.004 | 0.878 | 0.954 |
| Lamellar Thickness 28 | -0.011 | 0.712 | 0.887 | -0.002 | 0.949 | 0.974 |
| Lamellar Thickness 32 | 0.010 | 0.742 | 0.902 | 0.034 | 0.253 | 0.645 |
| Lamellar Thickness 36 | 0.025 | 0.395 | 0.783 | 0.008 | 0.785 | 0.916 |
| Lamellar Thickness 40 | 0.011 | 0.709 | 0.887 | -0.014 | 0.629 | 0.866 |
| Lamellar Thickness 44 | 0.054 | 0.065 | 0.377 | -0.002 | 0.948 | 0.974 |
| Lamellar Thickness 48 | 0.041 | 0.168 | 0.575 | -0.009 | 0.758 | 0.902 |
| Lamellar Thickness 52 | 0.031 | 0.293 | 0.672 | -0.016 | 0.578 | 0.860 |
| Lamellar Thickness 56 | 0.045 | 0.126 | 0.493 | -0.050 | 0.090 | 0.452 |
| Lamellar Thickness 60 | 0.033 | 0.261 | 0.645 | -0.064 | 0.029 | 0.246 |
| Lamellar Thickness 64 | -0.008 | 0.798 | 0.922 | -0.066 | 0.024 | 0.225 |
| Lamellar Thickness 68 | 0.027 | 0.361 | 0.773 | -0.014 | 0.642 | 0.866 |
| Lamellar Thickness 72 | -0.023 | 0.444 | 0.813 | 0.003 | 0.915 | 0.969 |
| Lamellar Thickness 76 | -0.005 | 0.864 | 0.951 | 0.026 | 0.385 | 0.775 |
| Lamellar Thickness 80 | -0.039 | 0.181 | 0.593 | -0.023 | 0.427 | 0.798 |
| Lamellar Thickness 84 | -0.020 | 0.500 | 0.838 | -0.013 | 0.659 | 0.866 |
| Lamellar Thickness 88 | 0.063 | 0.034 | 0.257 | 0.009 | 0.768 | 0.902 |
| Lamellar Thickness 92 | 0.079 | 0.007 | 0.154 | 0.015 | 0.600 | 0.860 |
| Lamellar Thickness 96 | 0.040 | 0.179 | 0.591 | 0.016 | 0.587 | 0.860 |
| Lamellar Thickness 100 | 0.001 | 0.970 | 0.980 | 0.035 | 0.230 | 0.645 |
| Lamellar Thickness 104 | -0.012 | 0.691 | 0.887 | 0.039 | 0.187 | 0.604 |
| Lamellar Thickness 108 | -0.009 | 0.756 | 0.902 | 0.017 | 0.573 | 0.860 |
| Lamellar Thickness 112 | 0.036 | 0.224 | 0.645 | 0.057 | 0.052 | 0.333 |
| Lamellar Thickness 111 | 0.026 | 0.383 | 0.775 | 0.035 | 0.227 | 0.645 |
| Lamellar Thickness 107 | 0.022 | 0.446 | 0.813 | 0.092 | 0.002 | 0.061 |
| Lamellar Thickness 103 | 0.016 | 0.589 | 0.860 | 0.072 | 0.014 | 0.185 |

| | | | | | | |
|---|---|---|---|---|---|---|
| Lamellar Thickness 99 | 0.025 | 0.394 | 0.783 | 0.075 | 0.011 | 0.169 |
| Lamellar Thickness 95 | 0.072 | 0.015 | 0.185 | 0.063 | 0.031 | 0.254 |
| Lamellar Thickness 91 | 0.076 | 0.009 | 0.169 | 0.082 | 0.005 | 0.128 |
| Lamellar Thickness 87 | 0.106 | 0.000 | 0.029 | 0.073 | 0.012 | 0.185 |
| Lamellar Thickness 83 | 0.043 | 0.141 | 0.520 | 0.062 | 0.034 | 0.257 |
| Lamellar Thickness 79 | 0.057 | 0.053 | 0.333 | -0.019 | 0.520 | 0.846 |
| Lamellar Thickness 75 | 0.006 | 0.835 | 0.943 | -0.011 | 0.705 | 0.887 |
| Lamellar Thickness 71 | 0.021 | 0.476 | 0.825 | 0.009 | 0.753 | 0.902 |
| Lamellar Thickness 67 | 0.046 | 0.116 | 0.493 | 0.008 | 0.793 | 0.921 |
| Lamellar Thickness 63 | 0.054 | 0.065 | 0.377 | 0.044 | 0.137 | 0.516 |
| Lamellar Thickness 59 | 0.038 | 0.198 | 0.632 | 0.018 | 0.549 | 0.860 |
| Lamellar Thickness 55 | 0.021 | 0.478 | 0.825 | -0.036 | 0.218 | 0.645 |
| Lamellar Thickness 51 | 0.001 | 0.978 | 0.985 | -0.014 | 0.641 | 0.866 |
| Lamellar Thickness 47 | 0.019 | 0.529 | 0.856 | -0.044 | 0.136 | 0.516 |
| Lamellar Thickness 43 | 0.003 | 0.929 | 0.974 | -0.030 | 0.306 | 0.683 |
| Lamellar Thickness 39 | 0.000 | 1.000 | 1.000 | -0.066 | 0.023 | 0.225 |
| Lamellar Thickness 35 | -0.033 | 0.256 | 0.645 | -0.063 | 0.032 | 0.257 |
| Lamellar Thickness 31 | 0.019 | 0.513 | 0.843 | 0.006 | 0.825 | 0.935 |
| Lamellar Thickness 27 | 0.005 | 0.870 | 0.951 | 0.015 | 0.599 | 0.860 |
| Lamellar Thickness 23 | 0.013 | 0.666 | 0.866 | 0.027 | 0.363 | 0.773 |
| Lamellar Thickness 19 | 0.034 | 0.247 | 0.645 | 0.086 | 0.003 | 0.093 |
| Lamellar Thickness 15 | 0.033 | 0.255 | 0.645 | 0.040 | 0.170 | 0.575 |
| Lamellar Thickness 11 | 0.003 | 0.931 | 0.974 | -0.007 | 0.807 | 0.922 |
| Lamellar Thickness 7 | -0.002 | 0.937 | 0.974 | -0.011 | 0.712 | 0.887 |
| Lamellar Thickness 3 | 0.095 | 0.001 | 0.051 | -0.017 | 0.559 | 0.860 |

24. Supplementary Table S12. Group-wise comparison of normative z-scores between SCZ and test HC

| | Left | | | Right | | |
|---|---|---|---|---|---|---|
| | effect size | P | corrected P | effect size | P | corrected P |
| Lamellar Width 1 | 0.011 | 0.747 | 0.812 | -0.048 | 0.159 | 0.267 |
| Lamellar Width 2 | -0.049 | 0.142 | 0.242 | -0.136 | 0.000 | 0.000 |
| Lamellar Width 3 | -0.097 | 0.004 | 0.012 | -0.109 | 0.001 | 0.005 |
| Lamellar Width 4 | -0.154 | 0.000 | 0.000 | -0.043 | 0.201 | 0.322 |
| Lamellar Width 5 | -0.154 | 0.000 | 0.000 | -0.144 | 0.000 | 0.000 |
| Lamellar Width 6 | -0.242 | 0.000 | 0.000 | -0.225 | 0.000 | 0.000 |
| Lamellar Width 7 | -0.255 | 0.000 | 0.000 | -0.240 | 0.000 | 0.000 |
| Lamellar Width 8 | -0.286 | 0.000 | 0.000 | -0.239 | 0.000 | 0.000 |
| Lamellar Width 9 | -0.270 | 0.000 | 0.000 | -0.243 | 0.000 | 0.000 |
| Lamellar Width 10 | -0.259 | 0.000 | 0.000 | -0.228 | 0.000 | 0.000 |
| Lamellar Width 11 | -0.211 | 0.000 | 0.000 | -0.208 | 0.000 | 0.000 |
| Lamellar Width 12 | -0.167 | 0.000 | 0.000 | -0.167 | 0.000 | 0.000 |
| Lamellar Width 13 | -0.201 | 0.000 | 0.000 | -0.172 | 0.000 | 0.000 |
| Lamellar Width 14 | -0.183 | 0.000 | 0.000 | -0.191 | 0.000 | 0.000 |
| Lamellar Width 15 | -0.191 | 0.000 | 0.000 | -0.154 | 0.000 | 0.000 |
| Lamellar Width 16 | -0.148 | 0.000 | 0.000 | -0.132 | 0.000 | 0.000 |
| Lamellar Width 17 | -0.150 | 0.000 | 0.000 | -0.133 | 0.000 | 0.000 |
| Lamellar Width 18 | -0.098 | 0.004 | 0.012 | -0.092 | 0.007 | 0.019 |
| Lamellar Width 19 | -0.096 | 0.004 | 0.014 | -0.053 | 0.119 | 0.206 |
| Lamellar Width 20 | -0.120 | 0.000 | 0.002 | -0.110 | 0.001 | 0.005 |
| Lamellar Width 21 | -0.093 | 0.006 | 0.016 | -0.073 | 0.030 | 0.071 |
| Lamellar Width 22 | -0.130 | 0.000 | 0.001 | -0.095 | 0.005 | 0.015 |
| Lamellar Width 23 | -0.123 | 0.000 | 0.001 | -0.113 | 0.001 | 0.004 |
| Lamellar Width 24 | -0.160 | 0.000 | 0.000 | -0.108 | 0.001 | 0.005 |

| | | | | | | |
|---|---|---|---|---|---|---|
| Lamellar Width 25 | -0.215 | 0.000 | 0.000 | -0.172 | 0.000 | 0.000 |
| Lamellar Width 26 | -0.226 | 0.000 | 0.000 | -0.172 | 0.000 | 0.000 |
| Lamellar Width 27 | -0.242 | 0.000 | 0.000 | -0.152 | 0.000 | 0.000 |
| Lamellar Width 28 | -0.209 | 0.000 | 0.000 | -0.134 | 0.000 | 0.000 |
| Lamellar Width 29 | -0.164 | 0.000 | 0.000 | -0.147 | 0.000 | 0.000 |
| Lamellar Width 30 | -0.101 | 0.003 | 0.009 | -0.108 | 0.001 | 0.005 |
| Lamellar Width 31 | -0.064 | 0.055 | 0.119 | 0.027 | 0.420 | 0.533 |
| Long-axis Length | -0.134 | 0.000 | 0.000 | -0.110 | 0.001 | 0.005 |
| Lamellar Thickness 2 | -0.100 | 0.003 | 0.010 | -0.177 | 0.000 | 0.000 |
| Lamellar Thickness 6 | -0.036 | 0.282 | 0.411 | 0.072 | 0.033 | 0.076 |
| Lamellar Thickness 10 | -0.023 | 0.495 | 0.605 | 0.031 | 0.363 | 0.488 |
| Lamellar Thickness 14 | 0.011 | 0.742 | 0.812 | 0.025 | 0.468 | 0.581 |
| Lamellar Thickness 18 | -0.009 | 0.788 | 0.843 | -0.059 | 0.081 | 0.155 |
| Lamellar Thickness 22 | 0.104 | 0.002 | 0.007 | -0.009 | 0.798 | 0.851 |
| Lamellar Thickness 26 | 0.169 | 0.000 | 0.000 | 0.007 | 0.841 | 0.877 |
| Lamellar Thickness 30 | 0.104 | 0.002 | 0.007 | 0.041 | 0.221 | 0.348 |
| Lamellar Thickness 34 | 0.107 | 0.001 | 0.005 | 0.030 | 0.368 | 0.488 |
| Lamellar Thickness 38 | 0.093 | 0.006 | 0.017 | -0.021 | 0.529 | 0.627 |
| Lamellar Thickness 42 | 0.113 | 0.001 | 0.003 | -0.062 | 0.066 | 0.134 |
| Lamellar Thickness 46 | 0.108 | 0.001 | 0.005 | -0.028 | 0.414 | 0.530 |
| Lamellar Thickness 50 | -0.031 | 0.355 | 0.482 | -0.016 | 0.628 | 0.717 |
| Lamellar Thickness 54 | -0.024 | 0.470 | 0.581 | -0.028 | 0.404 | 0.524 |
| Lamellar Thickness 58 | -0.068 | 0.042 | 0.093 | -0.118 | 0.000 | 0.002 |
| Lamellar Thickness 62 | -0.055 | 0.101 | 0.183 | -0.076 | 0.024 | 0.058 |
| Lamellar Thickness 66 | -0.023 | 0.485 | 0.594 | -0.067 | 0.047 | 0.103 |
| Lamellar Thickness 70 | -0.052 | 0.120 | 0.206 | -0.153 | 0.000 | 0.000 |
| Lamellar Thickness 74 | -0.118 | 0.000 | 0.002 | -0.147 | 0.000 | 0.000 |
| Lamellar Thickness 78 | -0.140 | 0.000 | 0.000 | -0.122 | 0.000 | 0.002 |
| Lamellar Thickness 82 | -0.076 | 0.024 | 0.058 | -0.118 | 0.000 | 0.002 |

| | | | | | | |
|---|---|---|---|---|---|---|
| Lamellar Thickness 86 | -0.080 | 0.017 | 0.042 | -0.058 | 0.084 | 0.158 |
| Lamellar Thickness 90 | -0.062 | 0.066 | 0.134 | -0.156 | 0.000 | 0.000 |
| Lamellar Thickness 94 | -0.055 | 0.105 | 0.188 | -0.095 | 0.005 | 0.015 |
| Lamellar Thickness 98 | -0.044 | 0.190 | 0.307 | -0.030 | 0.371 | 0.488 |
| Lamellar Thickness 102 | -0.030 | 0.369 | 0.488 | -0.041 | 0.229 | 0.355 |
| Lamellar Thickness 106 | -0.070 | 0.036 | 0.083 | -0.005 | 0.874 | 0.902 |
| Lamellar Thickness 110 | -0.070 | 0.038 | 0.087 | 0.035 | 0.294 | 0.421 |
| Lamellar Thickness 109 | -0.089 | 0.008 | 0.023 | -0.038 | 0.262 | 0.389 |
| Lamellar Thickness 105 | -0.079 | 0.019 | 0.046 | 0.035 | 0.298 | 0.425 |
| Lamellar Thickness 101 | -0.019 | 0.574 | 0.672 | 0.023 | 0.504 | 0.611 |
| Lamellar Thickness 97 | 0.007 | 0.825 | 0.870 | 0.011 | 0.753 | 0.816 |
| Lamellar Thickness 93 | -0.043 | 0.199 | 0.319 | -0.072 | 0.033 | 0.077 |
| Lamellar Thickness 89 | -0.060 | 0.074 | 0.145 | -0.106 | 0.002 | 0.006 |
| Lamellar Thickness 85 | -0.086 | 0.010 | 0.027 | -0.162 | 0.000 | 0.000 |
| Lamellar Thickness 81 | -0.004 | 0.908 | 0.927 | -0.135 | 0.000 | 0.000 |
| Lamellar Thickness 77 | -0.132 | 0.000 | 0.000 | -0.119 | 0.000 | 0.002 |
| Lamellar Thickness 73 | -0.105 | 0.002 | 0.007 | -0.095 | 0.005 | 0.015 |
| Lamellar Thickness 69 | -0.089 | 0.008 | 0.023 | -0.154 | 0.000 | 0.000 |
| Lamellar Thickness 65 | -0.034 | 0.313 | 0.438 | -0.130 | 0.000 | 0.001 |
| Lamellar Thickness 61 | -0.053 | 0.112 | 0.197 | -0.061 | 0.071 | 0.141 |
| Lamellar Thickness 57 | -0.022 | 0.513 | 0.614 | -0.100 | 0.003 | 0.010 |
| Lamellar Thickness 53 | -0.022 | 0.518 | 0.616 | -0.057 | 0.094 | 0.175 |
| Lamellar Thickness 49 | -0.027 | 0.416 | 0.530 | 0.035 | 0.300 | 0.425 |
| Lamellar Thickness 45 | 0.039 | 0.250 | 0.378 | 0.085 | 0.012 | 0.032 |
| Lamellar Thickness 41 | 0.017 | 0.621 | 0.712 | 0.088 | 0.009 | 0.024 |
| Lamellar Thickness 37 | -0.014 | 0.687 | 0.770 | 0.045 | 0.185 | 0.301 |
| Lamellar Thickness 33 | 0.052 | 0.123 | 0.210 | -0.018 | 0.598 | 0.694 |
| Lamellar Thickness 29 | -0.029 | 0.381 | 0.499 | -0.081 | 0.016 | 0.041 |
| Lamellar Thickness 25 | 0.010 | 0.770 | 0.828 | -0.061 | 0.072 | 0.143 |

| | | | | | | |
|---|---|---|---|---|---|---|
| Lamellar Thickness 21 | -0.003 | 0.929 | 0.945 | -0.043 | 0.204 | 0.325 |
| Lamellar Thickness 17 | -0.056 | 0.093 | 0.174 | 0.003 | 0.931 | 0.945 |
| Lamellar Thickness 13 | -0.054 | 0.108 | 0.191 | -0.002 | 0.954 | 0.964 |
| Lamellar Thickness 9 | -0.062 | 0.063 | 0.131 | -0.096 | 0.005 | 0.014 |
| Lamellar Thickness 5 | -0.080 | 0.018 | 0.043 | -0.101 | 0.003 | 0.009 |
| Lamellar Thickness 1 | -0.056 | 0.095 | 0.175 | -0.167 | 0.000 | 0.000 |
| Lamellar Thickness 4 | -0.075 | 0.026 | 0.062 | -0.109 | 0.001 | 0.005 |
| Lamellar Thickness 8 | -0.062 | 0.063 | 0.131 | 0.008 | 0.815 | 0.863 |
| Lamellar Thickness 12 | -0.103 | 0.002 | 0.007 | -0.084 | 0.013 | 0.033 |
| Lamellar Thickness 16 | -0.063 | 0.061 | 0.128 | -0.061 | 0.071 | 0.141 |
| Lamellar Thickness 20 | -0.030 | 0.368 | 0.488 | -0.035 | 0.305 | 0.431 |
| Lamellar Thickness 24 | 0.002 | 0.961 | 0.968 | -0.059 | 0.081 | 0.155 |
| Lamellar Thickness 28 | -0.011 | 0.747 | 0.812 | -0.026 | 0.441 | 0.552 |
| Lamellar Thickness 32 | 0.020 | 0.547 | 0.646 | 0.061 | 0.069 | 0.138 |
| Lamellar Thickness 36 | 0.054 | 0.106 | 0.190 | 0.026 | 0.435 | 0.548 |
| Lamellar Thickness 40 | 0.107 | 0.002 | 0.006 | 0.018 | 0.604 | 0.696 |
| Lamellar Thickness 44 | 0.080 | 0.017 | 0.042 | 0.031 | 0.359 | 0.485 |
| Lamellar Thickness 48 | 0.036 | 0.284 | 0.411 | 0.007 | 0.830 | 0.873 |
| Lamellar Thickness 52 | 0.048 | 0.151 | 0.254 | -0.023 | 0.505 | 0.611 |
| Lamellar Thickness 56 | 0.047 | 0.161 | 0.268 | -0.014 | 0.678 | 0.763 |
| Lamellar Thickness 60 | 0.024 | 0.482 | 0.593 | -0.037 | 0.273 | 0.401 |
| Lamellar Thickness 64 | -0.006 | 0.860 | 0.894 | -0.034 | 0.311 | 0.436 |
| Lamellar Thickness 68 | -0.001 | 0.986 | 0.986 | -0.064 | 0.058 | 0.123 |
| Lamellar Thickness 72 | -0.047 | 0.163 | 0.270 | -0.041 | 0.229 | 0.355 |
| Lamellar Thickness 76 | -0.039 | 0.242 | 0.369 | -0.062 | 0.068 | 0.138 |
| Lamellar Thickness 80 | -0.059 | 0.078 | 0.152 | -0.125 | 0.000 | 0.001 |
| Lamellar Thickness 84 | -0.095 | 0.005 | 0.015 | -0.087 | 0.010 | 0.027 |
| Lamellar Thickness 88 | -0.045 | 0.182 | 0.298 | -0.055 | 0.101 | 0.183 |
| Lamellar Thickness 92 | 0.001 | 0.986 | 0.986 | -0.054 | 0.107 | 0.190 |

| | | | | | | |
|---|---|---|---|---|---|---|
| Lamellar Thickness 96 | -0.034 | 0.317 | 0.441 | -0.064 | 0.057 | 0.122 |
| Lamellar Thickness 100 | -0.104 | 0.002 | 0.007 | -0.088 | 0.009 | 0.024 |
| Lamellar Thickness 104 | -0.083 | 0.014 | 0.035 | -0.058 | 0.084 | 0.158 |
| Lamellar Thickness 108 | -0.067 | 0.047 | 0.103 | -0.039 | 0.254 | 0.381 |
| Lamellar Thickness 112 | -0.040 | 0.234 | 0.361 | 0.032 | 0.342 | 0.470 |
| Lamellar Thickness 111 | -0.110 | 0.001 | 0.005 | -0.041 | 0.221 | 0.348 |
| Lamellar Thickness 107 | -0.069 | 0.039 | 0.088 | 0.015 | 0.656 | 0.744 |
| Lamellar Thickness 103 | -0.028 | 0.399 | 0.520 | -0.005 | 0.890 | 0.916 |
| Lamellar Thickness 99 | -0.036 | 0.289 | 0.416 | 0.046 | 0.170 | 0.279 |
| Lamellar Thickness 95 | -0.018 | 0.600 | 0.694 | 0.007 | 0.834 | 0.874 |
| Lamellar Thickness 91 | 0.056 | 0.097 | 0.178 | 0.010 | 0.763 | 0.823 |
| Lamellar Thickness 87 | 0.016 | 0.633 | 0.720 | 0.022 | 0.508 | 0.612 |
| Lamellar Thickness 83 | -0.030 | 0.368 | 0.488 | -0.022 | 0.511 | 0.613 |
| Lamellar Thickness 79 | 0.013 | 0.701 | 0.776 | 0.019 | 0.576 | 0.672 |
| Lamellar Thickness 75 | 0.008 | 0.804 | 0.854 | 0.015 | 0.661 | 0.746 |
| Lamellar Thickness 71 | 0.068 | 0.042 | 0.093 | 0.013 | 0.692 | 0.772 |
| Lamellar Thickness 67 | 0.107 | 0.001 | 0.005 | 0.036 | 0.284 | 0.411 |
| Lamellar Thickness 63 | 0.095 | 0.005 | 0.015 | 0.086 | 0.011 | 0.030 |
| Lamellar Thickness 59 | 0.086 | 0.010 | 0.028 | 0.067 | 0.048 | 0.104 |
| Lamellar Thickness 55 | 0.039 | 0.241 | 0.369 | -0.006 | 0.865 | 0.896 |
| Lamellar Thickness 51 | -0.013 | 0.707 | 0.780 | -0.013 | 0.698 | 0.776 |
| Lamellar Thickness 47 | -0.028 | 0.408 | 0.526 | -0.034 | 0.319 | 0.442 |
| Lamellar Thickness 43 | -0.116 | 0.001 | 0.002 | -0.053 | 0.114 | 0.200 |
| Lamellar Thickness 39 | -0.128 | 0.000 | 0.001 | -0.138 | 0.000 | 0.000 |
| Lamellar Thickness 35 | -0.090 | 0.008 | 0.022 | -0.093 | 0.006 | 0.017 |
| Lamellar Thickness 31 | -0.041 | 0.225 | 0.353 | -0.076 | 0.024 | 0.058 |
| Lamellar Thickness 27 | -0.049 | 0.144 | 0.244 | -0.038 | 0.263 | 0.389 |
| Lamellar Thickness 23 | -0.038 | 0.264 | 0.389 | -0.026 | 0.447 | 0.557 |
| Lamellar Thickness 19 | -0.004 | 0.905 | 0.927 | -0.032 | 0.348 | 0.475 |

| | | | | | | |
|---|---|---|---|---|---|---|
| Lamellar Thickness 15 | 0.027 | 0.422 | 0.534 | -0.059 | 0.079 | 0.154 |
| Lamellar Thickness 11 | 0.028 | 0.412 | 0.530 | -0.039 | 0.254 | 0.381 |
| Lamellar Thickness 7 | -0.012 | 0.726 | 0.798 | 0.020 | 0.551 | 0.647 |
| Lamellar Thickness 3 | -0.033 | 0.325 | 0.448 | -0.102 | 0.002 | 0.008 |

### 25. Supplementary Table S13. Group-wise comparison of normative z-scores between MCI and test HC

| | Left | | | Right | | |
|---|---|---|---|---|---|---|
| | effect size | P | corrected P | effect size | P | corrected P |
| Lamellar Width 1 | -0.113 | 0.006 | 0.014 | -0.158 | 0.000 | 0.001 |
| Lamellar Width 2 | -0.052 | 0.204 | 0.271 | -0.137 | 0.001 | 0.003 |
| Lamellar Width 3 | -0.077 | 0.063 | 0.095 | -0.141 | 0.001 | 0.003 |
| Lamellar Width 4 | 0.032 | 0.442 | 0.520 | -0.138 | 0.001 | 0.003 |
| Lamellar Width 5 | -0.040 | 0.333 | 0.403 | -0.120 | 0.004 | 0.009 |
| Lamellar Width 6 | -0.043 | 0.300 | 0.366 | -0.116 | 0.005 | 0.012 |
| Lamellar Width 7 | -0.226 | 0.000 | 0.000 | -0.201 | 0.000 | 0.000 |
| Lamellar Width 8 | -0.292 | 0.000 | 0.000 | -0.219 | 0.000 | 0.000 |
| Lamellar Width 9 | -0.312 | 0.000 | 0.000 | -0.228 | 0.000 | 0.000 |
| Lamellar Width 10 | -0.253 | 0.000 | 0.000 | -0.192 | 0.000 | 0.000 |
| Lamellar Width 11 | -0.300 | 0.000 | 0.000 | -0.205 | 0.000 | 0.000 |
| Lamellar Width 12 | -0.259 | 0.000 | 0.000 | -0.219 | 0.000 | 0.000 |
| Lamellar Width 13 | -0.273 | 0.000 | 0.000 | -0.233 | 0.000 | 0.000 |
| Lamellar Width 14 | -0.291 | 0.000 | 0.000 | -0.244 | 0.000 | 0.000 |
| Lamellar Width 15 | -0.310 | 0.000 | 0.000 | -0.180 | 0.000 | 0.000 |
| Lamellar Width 16 | -0.259 | 0.000 | 0.000 | -0.192 | 0.000 | 0.000 |
| Lamellar Width 17 | -0.211 | 0.000 | 0.000 | -0.147 | 0.000 | 0.002 |
| Lamellar Width 18 | -0.130 | 0.002 | 0.005 | -0.126 | 0.002 | 0.006 |
| Lamellar Width 19 | -0.091 | 0.027 | 0.049 | -0.048 | 0.252 | 0.322 |

| | | | | | | |
|---|---|---|---|---|---|---|
| Lamellar Width 20 | -0.140 | 0.001 | 0.002 | -0.137 | 0.001 | 0.003 |
| Lamellar Width 21 | -0.168 | 0.000 | 0.000 | -0.047 | 0.258 | 0.328 |
| Lamellar Width 22 | -0.137 | 0.001 | 0.003 | -0.040 | 0.338 | 0.407 |
| Lamellar Width 23 | -0.118 | 0.004 | 0.010 | -0.032 | 0.444 | 0.520 |
| Lamellar Width 24 | -0.142 | 0.001 | 0.002 | -0.123 | 0.003 | 0.008 |
| Lamellar Width 25 | -0.224 | 0.000 | 0.000 | -0.221 | 0.000 | 0.000 |
| Lamellar Width 26 | -0.208 | 0.000 | 0.000 | -0.240 | 0.000 | 0.000 |
| Lamellar Width 27 | -0.140 | 0.001 | 0.002 | -0.174 | 0.000 | 0.000 |
| Lamellar Width 28 | -0.122 | 0.003 | 0.008 | -0.094 | 0.024 | 0.043 |
| Lamellar Width 29 | -0.101 | 0.014 | 0.028 | -0.058 | 0.163 | 0.224 |
| Lamellar Width 30 | -0.067 | 0.105 | 0.153 | -0.096 | 0.020 | 0.038 |
| Lamellar Width 31 | -0.049 | 0.232 | 0.301 | 0.032 | 0.448 | 0.522 |
| Long-axis Length | -0.277 | 0.000 | 0.000 | -0.162 | 0.000 | 0.000 |
| Lamellar Thickness 2 | -0.050 | 0.222 | 0.290 | -0.090 | 0.030 | 0.053 |
| Lamellar Thickness 6 | -0.051 | 0.216 | 0.283 | -0.080 | 0.053 | 0.082 |
| Lamellar Thickness 10 | -0.036 | 0.384 | 0.457 | -0.014 | 0.744 | 0.776 |
| Lamellar Thickness 14 | -0.012 | 0.762 | 0.787 | -0.028 | 0.498 | 0.565 |
| Lamellar Thickness 18 | -0.101 | 0.014 | 0.028 | -0.117 | 0.005 | 0.011 |
| Lamellar Thickness 22 | -0.166 | 0.000 | 0.000 | -0.183 | 0.000 | 0.000 |
| Lamellar Thickness 26 | -0.078 | 0.058 | 0.088 | -0.085 | 0.040 | 0.065 |
| Lamellar Thickness 30 | -0.068 | 0.100 | 0.146 | -0.015 | 0.712 | 0.754 |
| Lamellar Thickness 34 | -0.043 | 0.300 | 0.366 | -0.045 | 0.276 | 0.344 |
| Lamellar Thickness 38 | -0.061 | 0.137 | 0.192 | -0.045 | 0.282 | 0.348 |
| Lamellar Thickness 42 | -0.147 | 0.000 | 0.001 | -0.208 | 0.000 | 0.000 |
| Lamellar Thickness 46 | -0.288 | 0.000 | 0.000 | -0.230 | 0.000 | 0.000 |
| Lamellar Thickness 50 | -0.331 | 0.000 | 0.000 | -0.288 | 0.000 | 0.000 |
| Lamellar Thickness 54 | -0.312 | 0.000 | 0.000 | -0.303 | 0.000 | 0.000 |
| Lamellar Thickness 58 | -0.327 | 0.000 | 0.000 | -0.320 | 0.000 | 0.000 |
| Lamellar Thickness 62 | -0.216 | 0.000 | 0.000 | -0.234 | 0.000 | 0.000 |

| | | | | | | |
|---|---|---|---|---|---|---|
| Lamellar Thickness 66 | -0.096 | 0.020 | 0.037 | -0.219 | 0.000 | 0.000 |
| Lamellar Thickness 70 | -0.036 | 0.380 | 0.454 | -0.108 | 0.009 | 0.020 |
| Lamellar Thickness 74 | 0.002 | 0.958 | 0.965 | -0.065 | 0.115 | 0.163 |
| Lamellar Thickness 78 | 0.046 | 0.260 | 0.328 | 0.025 | 0.548 | 0.609 |
| Lamellar Thickness 82 | 0.061 | 0.140 | 0.194 | 0.024 | 0.563 | 0.621 |
| Lamellar Thickness 86 | 0.032 | 0.438 | 0.517 | -0.056 | 0.179 | 0.243 |
| Lamellar Thickness 90 | 0.019 | 0.647 | 0.699 | -0.105 | 0.011 | 0.023 |
| Lamellar Thickness 94 | 0.018 | 0.656 | 0.705 | -0.081 | 0.050 | 0.078 |
| Lamellar Thickness 98 | -0.001 | 0.978 | 0.982 | -0.072 | 0.084 | 0.125 |
| Lamellar Thickness 102 | 0.095 | 0.022 | 0.040 | 0.025 | 0.555 | 0.615 |
| Lamellar Thickness 106 | 0.103 | 0.013 | 0.026 | 0.080 | 0.053 | 0.082 |
| Lamellar Thickness 110 | 0.066 | 0.108 | 0.156 | 0.045 | 0.277 | 0.344 |
| Lamellar Thickness 109 | 0.014 | 0.733 | 0.770 | -0.062 | 0.138 | 0.193 |
| Lamellar Thickness 105 | 0.048 | 0.244 | 0.313 | -0.019 | 0.648 | 0.699 |
| Lamellar Thickness 101 | 0.073 | 0.077 | 0.115 | -0.017 | 0.683 | 0.729 |
| Lamellar Thickness 97 | 0.009 | 0.829 | 0.850 | -0.088 | 0.035 | 0.059 |
| Lamellar Thickness 93 | 0.046 | 0.261 | 0.328 | -0.111 | 0.008 | 0.016 |
| Lamellar Thickness 89 | 0.027 | 0.520 | 0.583 | -0.085 | 0.040 | 0.064 |
| Lamellar Thickness 85 | 0.027 | 0.518 | 0.582 | -0.087 | 0.037 | 0.061 |
| Lamellar Thickness 81 | -0.005 | 0.900 | 0.912 | -0.090 | 0.030 | 0.053 |
| Lamellar Thickness 77 | 0.051 | 0.214 | 0.281 | -0.065 | 0.115 | 0.163 |
| Lamellar Thickness 73 | -0.031 | 0.458 | 0.529 | -0.060 | 0.146 | 0.202 |
| Lamellar Thickness 69 | -0.057 | 0.167 | 0.228 | -0.091 | 0.028 | 0.049 |
| Lamellar Thickness 65 | -0.111 | 0.007 | 0.015 | -0.110 | 0.008 | 0.017 |
| Lamellar Thickness 61 | -0.116 | 0.005 | 0.011 | -0.096 | 0.021 | 0.039 |
| Lamellar Thickness 57 | -0.132 | 0.001 | 0.004 | -0.084 | 0.044 | 0.070 |
| Lamellar Thickness 53 | -0.146 | 0.000 | 0.002 | -0.121 | 0.004 | 0.009 |
| Lamellar Thickness 49 | -0.179 | 0.000 | 0.000 | -0.164 | 0.000 | 0.000 |
| Lamellar Thickness 45 | -0.164 | 0.000 | 0.000 | -0.140 | 0.001 | 0.002 |

| | | | | | | |
|---|---|---|---|---|---|---|
| Lamellar Thickness 41 | -0.209 | 0.000 | 0.000 | -0.147 | 0.000 | 0.002 |
| Lamellar Thickness 37 | -0.217 | 0.000 | 0.000 | -0.041 | 0.328 | 0.398 |
| Lamellar Thickness 33 | -0.221 | 0.000 | 0.000 | -0.066 | 0.114 | 0.163 |
| Lamellar Thickness 29 | -0.152 | 0.000 | 0.001 | -0.074 | 0.076 | 0.114 |
| Lamellar Thickness 25 | -0.115 | 0.005 | 0.012 | -0.094 | 0.024 | 0.043 |
| Lamellar Thickness 21 | -0.120 | 0.004 | 0.009 | -0.146 | 0.000 | 0.002 |
| Lamellar Thickness 17 | -0.100 | 0.016 | 0.030 | -0.118 | 0.004 | 0.010 |
| Lamellar Thickness 13 | -0.087 | 0.035 | 0.059 | -0.100 | 0.016 | 0.030 |
| Lamellar Thickness 9 | -0.107 | 0.009 | 0.020 | -0.168 | 0.000 | 0.000 |
| Lamellar Thickness 5 | -0.135 | 0.001 | 0.003 | -0.107 | 0.010 | 0.020 |
| Lamellar Thickness 1 | -0.202 | 0.000 | 0.000 | -0.145 | 0.000 | 0.002 |
| Lamellar Thickness 4 | -0.126 | 0.002 | 0.006 | -0.087 | 0.035 | 0.059 |
| Lamellar Thickness 8 | -0.053 | 0.202 | 0.270 | -0.107 | 0.010 | 0.021 |
| Lamellar Thickness 12 | -0.068 | 0.099 | 0.145 | -0.082 | 0.047 | 0.074 |
| Lamellar Thickness 16 | -0.152 | 0.000 | 0.001 | -0.177 | 0.000 | 0.000 |
| Lamellar Thickness 20 | -0.218 | 0.000 | 0.000 | -0.169 | 0.000 | 0.000 |
| Lamellar Thickness 24 | -0.292 | 0.000 | 0.000 | -0.266 | 0.000 | 0.000 |
| Lamellar Thickness 28 | -0.256 | 0.000 | 0.000 | -0.150 | 0.000 | 0.001 |
| Lamellar Thickness 32 | -0.132 | 0.001 | 0.004 | -0.120 | 0.004 | 0.009 |
| Lamellar Thickness 36 | -0.089 | 0.030 | 0.053 | -0.090 | 0.031 | 0.054 |
| Lamellar Thickness 40 | -0.097 | 0.019 | 0.036 | -0.027 | 0.508 | 0.574 |
| Lamellar Thickness 44 | -0.077 | 0.062 | 0.094 | -0.106 | 0.011 | 0.022 |
| Lamellar Thickness 48 | -0.052 | 0.209 | 0.276 | -0.130 | 0.002 | 0.005 |
| Lamellar Thickness 52 | -0.058 | 0.158 | 0.218 | -0.122 | 0.003 | 0.008 |
| Lamellar Thickness 56 | -0.138 | 0.001 | 0.003 | -0.116 | 0.005 | 0.012 |
| Lamellar Thickness 60 | -0.123 | 0.003 | 0.007 | -0.097 | 0.020 | 0.037 |
| Lamellar Thickness 64 | -0.068 | 0.097 | 0.143 | -0.049 | 0.242 | 0.312 |
| Lamellar Thickness 68 | 0.013 | 0.746 | 0.776 | -0.009 | 0.835 | 0.853 |
| Lamellar Thickness 72 | -0.018 | 0.658 | 0.705 | -0.013 | 0.750 | 0.777 |

| | | | | | | |
|---|---|---|---|---|---|---|
| Lamellar Thickness 76 | -0.036 | 0.389 | 0.461 | -0.021 | 0.614 | 0.670 |
| Lamellar Thickness 80 | 0.016 | 0.692 | 0.735 | -0.053 | 0.198 | 0.265 |
| Lamellar Thickness 84 | 0.130 | 0.002 | 0.005 | 0.123 | 0.003 | 0.008 |
| Lamellar Thickness 88 | -0.026 | 0.536 | 0.598 | -0.014 | 0.744 | 0.776 |
| Lamellar Thickness 92 | 0.029 | 0.480 | 0.551 | -0.022 | 0.603 | 0.660 |
| Lamellar Thickness 96 | -0.131 | 0.002 | 0.004 | -0.194 | 0.000 | 0.000 |
| Lamellar Thickness 100 | -0.161 | 0.000 | 0.000 | -0.164 | 0.000 | 0.000 |
| Lamellar Thickness 104 | -0.125 | 0.002 | 0.006 | -0.137 | 0.001 | 0.003 |
| Lamellar Thickness 108 | 0.005 | 0.904 | 0.914 | -0.089 | 0.031 | 0.054 |
| Lamellar Thickness 112 | 0.029 | 0.485 | 0.554 | 0.022 | 0.601 | 0.660 |
| Lamellar Thickness 111 | 0.020 | 0.634 | 0.689 | -0.103 | 0.013 | 0.026 |
| Lamellar Thickness 107 | 0.014 | 0.729 | 0.769 | -0.087 | 0.036 | 0.061 |
| Lamellar Thickness 103 | -0.054 | 0.193 | 0.261 | -0.105 | 0.011 | 0.023 |
| Lamellar Thickness 99 | -0.006 | 0.893 | 0.909 | -0.163 | 0.000 | 0.000 |
| Lamellar Thickness 95 | -0.090 | 0.029 | 0.052 | -0.121 | 0.004 | 0.009 |
| Lamellar Thickness 91 | -0.085 | 0.039 | 0.064 | -0.117 | 0.005 | 0.011 |
| Lamellar Thickness 87 | -0.010 | 0.803 | 0.826 | -0.038 | 0.355 | 0.426 |
| Lamellar Thickness 83 | 0.000 | 0.990 | 0.990 | -0.082 | 0.049 | 0.076 |
| Lamellar Thickness 79 | 0.128 | 0.002 | 0.005 | 0.078 | 0.061 | 0.094 |
| Lamellar Thickness 75 | 0.133 | 0.001 | 0.004 | 0.128 | 0.002 | 0.006 |
| Lamellar Thickness 71 | 0.134 | 0.001 | 0.003 | 0.115 | 0.006 | 0.012 |
| Lamellar Thickness 67 | 0.031 | 0.453 | 0.526 | 0.046 | 0.266 | 0.333 |
| Lamellar Thickness 63 | -0.099 | 0.017 | 0.032 | -0.031 | 0.462 | 0.532 |
| Lamellar Thickness 59 | -0.126 | 0.002 | 0.006 | -0.085 | 0.041 | 0.066 |
| Lamellar Thickness 55 | -0.084 | 0.042 | 0.067 | -0.102 | 0.014 | 0.027 |
| Lamellar Thickness 51 | -0.082 | 0.047 | 0.074 | -0.088 | 0.035 | 0.059 |
| Lamellar Thickness 47 | -0.123 | 0.003 | 0.008 | -0.047 | 0.260 | 0.328 |
| Lamellar Thickness 43 | -0.086 | 0.037 | 0.061 | -0.029 | 0.489 | 0.557 |
| Lamellar Thickness 39 | -0.066 | 0.109 | 0.158 | -0.095 | 0.022 | 0.040 |

| | | | | | | |
|---|---|---|---|---|---|---|
| Lamellar Thickness 35 | -0.092 | 0.026 | 0.047 | -0.153 | 0.000 | 0.001 |
| Lamellar Thickness 31 | -0.144 | 0.000 | 0.002 | -0.171 | 0.000 | 0.000 |
| Lamellar Thickness 27 | -0.181 | 0.000 | 0.000 | -0.156 | 0.000 | 0.001 |
| Lamellar Thickness 23 | -0.229 | 0.000 | 0.000 | -0.115 | 0.006 | 0.013 |
| Lamellar Thickness 19 | -0.200 | 0.000 | 0.000 | -0.143 | 0.001 | 0.002 |
| Lamellar Thickness 15 | -0.161 | 0.000 | 0.000 | -0.125 | 0.003 | 0.007 |
| Lamellar Thickness 11 | -0.136 | 0.001 | 0.003 | -0.045 | 0.280 | 0.346 |
| Lamellar Thickness 7 | -0.063 | 0.128 | 0.180 | -0.053 | 0.198 | 0.265 |
| Lamellar Thickness 3 | -0.162 | 0.000 | 0.000 | -0.129 | 0.002 | 0.005 |

### 26. Supplementary Table S14. Group-wise comparison of normative z-scores between DM and test HC

| | Left | | | Right | | |
|---|---|---|---|---|---|---|
| | effect size | P | corrected P | effect size | P | corrected P |
| Lamellar Width 1 | -0.129 | 0.000 | 0.000 | -0.163 | 0.000 | 0.000 |
| Lamellar Width 2 | -0.046 | 0.109 | 0.125 | -0.096 | 0.001 | 0.001 |
| Lamellar Width 3 | -0.047 | 0.106 | 0.121 | -0.117 | 0.000 | 0.000 |
| Lamellar Width 4 | -0.102 | 0.000 | 0.001 | -0.081 | 0.005 | 0.007 |
| Lamellar Width 5 | -0.157 | 0.000 | 0.000 | -0.189 | 0.000 | 0.000 |
| Lamellar Width 6 | -0.113 | 0.000 | 0.000 | -0.185 | 0.000 | 0.000 |
| Lamellar Width 7 | -0.286 | 0.000 | 0.000 | -0.298 | 0.000 | 0.000 |
| Lamellar Width 8 | -0.395 | 0.000 | 0.000 | -0.326 | 0.000 | 0.000 |
| Lamellar Width 9 | -0.426 | 0.000 | 0.000 | -0.346 | 0.000 | 0.000 |
| Lamellar Width 10 | -0.365 | 0.000 | 0.000 | -0.259 | 0.000 | 0.000 |
| Lamellar Width 11 | -0.379 | 0.000 | 0.000 | -0.246 | 0.000 | 0.000 |
| Lamellar Width 12 | -0.313 | 0.000 | 0.000 | -0.233 | 0.000 | 0.000 |
| Lamellar Width 13 | -0.328 | 0.000 | 0.000 | -0.286 | 0.000 | 0.000 |

| | | | | | | |
|---|---|---|---|---|---|---|
| Lamellar Width 14 | -0.305 | 0.000 | 0.000 | -0.325 | 0.000 | 0.000 |
| Lamellar Width 15 | -0.299 | 0.000 | 0.000 | -0.306 | 0.000 | 0.000 |
| Lamellar Width 16 | -0.288 | 0.000 | 0.000 | -0.209 | 0.000 | 0.000 |
| Lamellar Width 17 | -0.207 | 0.000 | 0.000 | -0.162 | 0.000 | 0.000 |
| Lamellar Width 18 | -0.142 | 0.000 | 0.000 | -0.128 | 0.000 | 0.000 |
| Lamellar Width 19 | -0.086 | 0.003 | 0.004 | -0.125 | 0.000 | 0.000 |
| Lamellar Width 20 | -0.145 | 0.000 | 0.000 | -0.151 | 0.000 | 0.000 |
| Lamellar Width 21 | -0.187 | 0.000 | 0.000 | -0.145 | 0.000 | 0.000 |
| Lamellar Width 22 | -0.070 | 0.016 | 0.021 | -0.027 | 0.353 | 0.381 |
| Lamellar Width 23 | -0.153 | 0.000 | 0.000 | 0.018 | 0.534 | 0.547 |
| Lamellar Width 24 | -0.193 | 0.000 | 0.000 | -0.058 | 0.044 | 0.054 |
| Lamellar Width 25 | -0.261 | 0.000 | 0.000 | -0.182 | 0.000 | 0.000 |
| Lamellar Width 26 | -0.206 | 0.000 | 0.000 | -0.262 | 0.000 | 0.000 |
| Lamellar Width 27 | -0.146 | 0.000 | 0.000 | -0.163 | 0.000 | 0.000 |
| Lamellar Width 28 | -0.150 | 0.000 | 0.000 | -0.132 | 0.000 | 0.000 |
| Lamellar Width 29 | -0.072 | 0.013 | 0.017 | -0.103 | 0.000 | 0.001 |
| Lamellar Width 30 | -0.071 | 0.014 | 0.018 | -0.089 | 0.002 | 0.003 |
| Lamellar Width 31 | -0.025 | 0.381 | 0.408 | -0.033 | 0.246 | 0.269 |
| Long-axis Length | -0.343 | 0.000 | 0.000 | -0.317 | 0.000 | 0.000 |
| Lamellar Thickness 2 | 0.007 | 0.796 | 0.801 | -0.104 | 0.000 | 0.000 |
| Lamellar Thickness 6 | -0.052 | 0.073 | 0.087 | 0.027 | 0.352 | 0.381 |
| Lamellar Thickness 10 | -0.086 | 0.003 | 0.004 | -0.059 | 0.041 | 0.051 |
| Lamellar Thickness 14 | -0.136 | 0.000 | 0.000 | -0.075 | 0.009 | 0.012 |
| Lamellar Thickness 18 | -0.198 | 0.000 | 0.000 | -0.209 | 0.000 | 0.000 |
| Lamellar Thickness 22 | -0.241 | 0.000 | 0.000 | -0.321 | 0.000 | 0.000 |
| Lamellar Thickness 26 | -0.128 | 0.000 | 0.000 | -0.218 | 0.000 | 0.000 |
| Lamellar Thickness 30 | -0.143 | 0.000 | 0.000 | -0.187 | 0.000 | 0.000 |
| Lamellar Thickness 34 | -0.118 | 0.000 | 0.000 | -0.210 | 0.000 | 0.000 |
| Lamellar Thickness 38 | -0.185 | 0.000 | 0.000 | -0.258 | 0.000 | 0.000 |
| Lamellar Thickness 42 | -0.223 | 0.000 | 0.000 | -0.319 | 0.000 | 0.000 |

| | | | | | | |
|---|---|---|---|---|---|---|
| Lamellar Thickness 46 | -0.371 | 0.000 | 0.000 | -0.315 | 0.000 | 0.000 |
| Lamellar Thickness 50 | -0.448 | 0.000 | 0.000 | -0.336 | 0.000 | 0.000 |
| Lamellar Thickness 54 | -0.458 | 0.000 | 0.000 | -0.362 | 0.000 | 0.000 |
| Lamellar Thickness 58 | -0.436 | 0.000 | 0.000 | -0.369 | 0.000 | 0.000 |
| Lamellar Thickness 62 | -0.349 | 0.000 | 0.000 | -0.316 | 0.000 | 0.000 |
| Lamellar Thickness 66 | -0.244 | 0.000 | 0.000 | -0.260 | 0.000 | 0.000 |
| Lamellar Thickness 70 | -0.048 | 0.094 | 0.110 | -0.173 | 0.000 | 0.000 |
| Lamellar Thickness 74 | -0.023 | 0.429 | 0.447 | -0.114 | 0.000 | 0.000 |
| Lamellar Thickness 78 | 0.022 | 0.450 | 0.464 | -0.045 | 0.114 | 0.130 |
| Lamellar Thickness 82 | 0.053 | 0.067 | 0.080 | -0.059 | 0.039 | 0.049 |
| Lamellar Thickness 86 | 0.052 | 0.074 | 0.088 | -0.067 | 0.019 | 0.024 |
| Lamellar Thickness 90 | -0.035 | 0.225 | 0.249 | -0.133 | 0.000 | 0.000 |
| Lamellar Thickness 94 | -0.073 | 0.011 | 0.015 | -0.200 | 0.000 | 0.000 |
| Lamellar Thickness 98 | -0.024 | 0.399 | 0.426 | -0.135 | 0.000 | 0.000 |
| Lamellar Thickness 102 | 0.080 | 0.006 | 0.008 | -0.032 | 0.260 | 0.284 |
| Lamellar Thickness 106 | 0.116 | 0.000 | 0.000 | 0.048 | 0.097 | 0.113 |
| Lamellar Thickness 110 | 0.107 | 0.000 | 0.000 | 0.028 | 0.335 | 0.364 |
| Lamellar Thickness 109 | -0.005 | 0.870 | 0.870 | -0.038 | 0.193 | 0.215 |
| Lamellar Thickness 105 | 0.050 | 0.081 | 0.096 | 0.044 | 0.124 | 0.140 |
| Lamellar Thickness 101 | 0.022 | 0.442 | 0.458 | 0.012 | 0.683 | 0.695 |
| Lamellar Thickness 97 | -0.023 | 0.417 | 0.440 | -0.023 | 0.427 | 0.447 |
| Lamellar Thickness 93 | -0.023 | 0.424 | 0.446 | -0.118 | 0.000 | 0.000 |
| Lamellar Thickness 89 | -0.009 | 0.763 | 0.771 | -0.143 | 0.000 | 0.000 |
| Lamellar Thickness 85 | -0.071 | 0.013 | 0.018 | -0.134 | 0.000 | 0.000 |
| Lamellar Thickness 81 | -0.054 | 0.062 | 0.075 | -0.075 | 0.009 | 0.012 |
| Lamellar Thickness 77 | -0.051 | 0.079 | 0.093 | -0.130 | 0.000 | 0.000 |
| Lamellar Thickness 73 | -0.148 | 0.000 | 0.000 | -0.115 | 0.000 | 0.000 |
| Lamellar Thickness 69 | -0.185 | 0.000 | 0.000 | -0.223 | 0.000 | 0.000 |
| Lamellar Thickness 65 | -0.201 | 0.000 | 0.000 | -0.236 | 0.000 | 0.000 |
| Lamellar Thickness 61 | -0.211 | 0.000 | 0.000 | -0.185 | 0.000 | 0.000 |

| | | | | | | |
|---|---|---|---|---|---|---|
| Lamellar Thickness 57 | -0.169 | 0.000 | 0.000 | -0.175 | 0.000 | 0.000 |
| Lamellar Thickness 53 | -0.145 | 0.000 | 0.000 | -0.177 | 0.000 | 0.000 |
| Lamellar Thickness 49 | -0.213 | 0.000 | 0.000 | -0.183 | 0.000 | 0.000 |
| Lamellar Thickness 45 | -0.181 | 0.000 | 0.000 | -0.224 | 0.000 | 0.000 |
| Lamellar Thickness 41 | -0.177 | 0.000 | 0.000 | -0.145 | 0.000 | 0.000 |
| Lamellar Thickness 37 | -0.253 | 0.000 | 0.000 | -0.049 | 0.086 | 0.101 |
| Lamellar Thickness 33 | -0.245 | 0.000 | 0.000 | -0.057 | 0.046 | 0.056 |
| Lamellar Thickness 29 | -0.238 | 0.000 | 0.000 | -0.087 | 0.003 | 0.004 |
| Lamellar Thickness 25 | -0.200 | 0.000 | 0.000 | -0.175 | 0.000 | 0.000 |
| Lamellar Thickness 21 | -0.214 | 0.000 | 0.000 | -0.121 | 0.000 | 0.000 |
| Lamellar Thickness 17 | -0.159 | 0.000 | 0.000 | -0.095 | 0.001 | 0.001 |
| Lamellar Thickness 13 | -0.115 | 0.000 | 0.000 | -0.180 | 0.000 | 0.000 |
| Lamellar Thickness 9 | -0.205 | 0.000 | 0.000 | -0.277 | 0.000 | 0.000 |
| Lamellar Thickness 5 | -0.232 | 0.000 | 0.000 | -0.264 | 0.000 | 0.000 |
| Lamellar Thickness 1 | -0.287 | 0.000 | 0.000 | -0.329 | 0.000 | 0.000 |
| Lamellar Thickness 4 | -0.065 | 0.025 | 0.032 | -0.153 | 0.000 | 0.000 |
| Lamellar Thickness 8 | -0.180 | 0.000 | 0.000 | -0.101 | 0.000 | 0.001 |
| Lamellar Thickness 12 | -0.099 | 0.001 | 0.001 | -0.110 | 0.000 | 0.000 |
| Lamellar Thickness 16 | -0.231 | 0.000 | 0.000 | -0.229 | 0.000 | 0.000 |
| Lamellar Thickness 20 | -0.382 | 0.000 | 0.000 | -0.294 | 0.000 | 0.000 |
| Lamellar Thickness 24 | -0.422 | 0.000 | 0.000 | -0.409 | 0.000 | 0.000 |
| Lamellar Thickness 28 | -0.371 | 0.000 | 0.000 | -0.344 | 0.000 | 0.000 |
| Lamellar Thickness 32 | -0.264 | 0.000 | 0.000 | -0.317 | 0.000 | 0.000 |
| Lamellar Thickness 36 | -0.143 | 0.000 | 0.000 | -0.238 | 0.000 | 0.000 |
| Lamellar Thickness 40 | -0.137 | 0.000 | 0.000 | -0.193 | 0.000 | 0.000 |
| Lamellar Thickness 44 | -0.163 | 0.000 | 0.000 | -0.311 | 0.000 | 0.000 |
| Lamellar Thickness 48 | -0.149 | 0.000 | 0.000 | -0.261 | 0.000 | 0.000 |
| Lamellar Thickness 52 | -0.178 | 0.000 | 0.000 | -0.235 | 0.000 | 0.000 |
| Lamellar Thickness 56 | -0.225 | 0.000 | 0.000 | -0.284 | 0.000 | 0.000 |
| Lamellar Thickness 60 | -0.202 | 0.000 | 0.000 | -0.264 | 0.000 | 0.000 |

| | | | | | | |
|---|---|---|---|---|---|---|
| Lamellar Thickness 64 | -0.171 | 0.000 | 0.000 | -0.182 | 0.000 | 0.000 |
| Lamellar Thickness 68 | -0.087 | 0.003 | 0.004 | -0.052 | 0.069 | 0.082 |
| Lamellar Thickness 72 | -0.034 | 0.236 | 0.259 | -0.092 | 0.001 | 0.002 |
| Lamellar Thickness 76 | -0.026 | 0.374 | 0.402 | -0.079 | 0.006 | 0.008 |
| Lamellar Thickness 80 | 0.014 | 0.621 | 0.635 | -0.103 | 0.000 | 0.001 |
| Lamellar Thickness 84 | 0.118 | 0.000 | 0.000 | 0.074 | 0.010 | 0.013 |
| Lamellar Thickness 88 | -0.056 | 0.052 | 0.063 | -0.082 | 0.005 | 0.006 |
| Lamellar Thickness 92 | 0.103 | 0.000 | 0.001 | -0.024 | 0.402 | 0.427 |
| Lamellar Thickness 96 | -0.178 | 0.000 | 0.000 | -0.283 | 0.000 | 0.000 |
| Lamellar Thickness 100 | -0.176 | 0.000 | 0.000 | -0.181 | 0.000 | 0.000 |
| Lamellar Thickness 104 | -0.208 | 0.000 | 0.000 | -0.204 | 0.000 | 0.000 |
| Lamellar Thickness 108 | -0.011 | 0.699 | 0.709 | -0.094 | 0.001 | 0.002 |
| Lamellar Thickness 112 | 0.073 | 0.012 | 0.015 | -0.053 | 0.064 | 0.078 |
| Lamellar Thickness 111 | -0.047 | 0.102 | 0.118 | -0.042 | 0.147 | 0.166 |
| Lamellar Thickness 107 | -0.037 | 0.206 | 0.229 | -0.040 | 0.160 | 0.180 |
| Lamellar Thickness 103 | -0.068 | 0.018 | 0.023 | -0.080 | 0.006 | 0.008 |
| Lamellar Thickness 99 | -0.070 | 0.015 | 0.020 | -0.069 | 0.016 | 0.021 |
| Lamellar Thickness 95 | 0.006 | 0.844 | 0.847 | -0.121 | 0.000 | 0.000 |
| Lamellar Thickness 91 | -0.036 | 0.217 | 0.241 | -0.203 | 0.000 | 0.000 |
| Lamellar Thickness 87 | 0.038 | 0.185 | 0.208 | -0.047 | 0.099 | 0.115 |
| Lamellar Thickness 83 | 0.023 | 0.432 | 0.449 | -0.103 | 0.000 | 0.000 |
| Lamellar Thickness 79 | 0.254 | 0.000 | 0.000 | 0.129 | 0.000 | 0.000 |
| Lamellar Thickness 75 | 0.231 | 0.000 | 0.000 | 0.082 | 0.004 | 0.006 |
| Lamellar Thickness 71 | 0.236 | 0.000 | 0.000 | 0.113 | 0.000 | 0.000 |
| Lamellar Thickness 67 | 0.094 | 0.001 | 0.002 | 0.023 | 0.417 | 0.440 |
| Lamellar Thickness 63 | -0.067 | 0.020 | 0.026 | -0.057 | 0.046 | 0.056 |
| Lamellar Thickness 59 | -0.066 | 0.023 | 0.029 | -0.115 | 0.000 | 0.000 |
| Lamellar Thickness 55 | -0.021 | 0.476 | 0.490 | -0.118 | 0.000 | 0.000 |
| Lamellar Thickness 51 | -0.058 | 0.044 | 0.054 | -0.126 | 0.000 | 0.000 |
| Lamellar Thickness 47 | -0.106 | 0.000 | 0.000 | -0.107 | 0.000 | 0.000 |

| | | | | | | |
|---|---|---|---|---|---|---|
| Lamellar Thickness 43 | -0.098 | 0.001 | 0.001 | -0.115 | 0.000 | 0.000 |
| Lamellar Thickness 39 | -0.147 | 0.000 | 0.000 | -0.248 | 0.000 | 0.000 |
| Lamellar Thickness 35 | -0.260 | 0.000 | 0.000 | -0.261 | 0.000 | 0.000 |
| Lamellar Thickness 31 | -0.202 | 0.000 | 0.000 | -0.304 | 0.000 | 0.000 |
| Lamellar Thickness 27 | -0.271 | 0.000 | 0.000 | -0.320 | 0.000 | 0.000 |
| Lamellar Thickness 23 | -0.309 | 0.000 | 0.000 | -0.323 | 0.000 | 0.000 |
| Lamellar Thickness 19 | -0.302 | 0.000 | 0.000 | -0.347 | 0.000 | 0.000 |
| Lamellar Thickness 15 | -0.286 | 0.000 | 0.000 | -0.340 | 0.000 | 0.000 |
| Lamellar Thickness 11 | -0.280 | 0.000 | 0.000 | -0.251 | 0.000 | 0.000 |
| Lamellar Thickness 7 | -0.134 | 0.000 | 0.000 | -0.204 | 0.000 | 0.000 |
| Lamellar Thickness 3 | -0.228 | 0.000 | 0.000 | -0.315 | 0.000 | 0.000 |

### 27. Supplementary Table S15. Group-wise composite zscore comparison

| | | Left | | | Right | | |
|---|---|---|---|---|---|---|---|
| group | feature | effect size | P | corrected P | effect size | P | corrected P |
| ASD | Lamellar Thickness.Head | 0.068 | 0.004 | 0.004 | 0.011 | 0.635 | 0.318 |
| | Lamellar Thickness.Body | 0.108 | 0.000 | 0.000 | 0.060 | 0.010 | 0.014 |
| | Lamellar Thickness.Tail | 0.118 | 0.000 | 0.000 | 0.023 | 0.306 | 0.184 |
| | Lamellar Width.Head | 0.072 | 0.002 | 0.003 | 0.050 | 0.039 | 0.029 |
| | Lamellar Width.Body | 0.090 | 0.000 | 0.000 | 0.065 | 0.005 | 0.014 |
| | Lamellar Width.Tail | 0.059 | 0.013 | 0.013 | 0.056 | 0.016 | 0.016 |
| ADHD | Lamellar Thickness.Head | 0.078 | 0.000 | 0.000 | 0.028 | 0.138 | 0.152 |
| | Lamellar Thickness.Body | 0.125 | 0.000 | 0.000 | 0.020 | 0.291 | 0.194 |
| | Lamellar Thickness.Tail | 0.074 | 0.000 | 0.000 | 0.024 | 0.199 | 0.159 |
| | Lamellar Width.Head | 0.000 | 0.997 | 0.166 | 0.055 | 0.003 | 0.013 |
| | Lamellar Width.Body | 0.047 | 0.012 | 0.002 | 0.027 | 0.152 | 0.152 |
| | Lamellar Width.Tail | 0.056 | 0.003 | 0.001 | 0.047 | 0.012 | 0.024 |

| | | | | | | | |
|---|---|---|---|---|---|---|---|
| ANX | Lamellar Thickness.Head | 0.036 | 0.294 | 0.367 | 0.032 | 0.337 | 0.337 |
| | Lamellar Thickness.Body | 0.111 | 0.001 | 0.003 | 0.043 | 0.192 | 0.337 |
| | Lamellar Thickness.Tail | 0.018 | 0.581 | 0.503 | -0.034 | 0.289 | 0.337 |
| | Lamellar Width.Head | -0.018 | 0.603 | 0.503 | 0.040 | 0.240 | 0.337 |
| | Lamellar Width.Body | 0.075 | 0.032 | 0.081 | 0.043 | 0.212 | 0.337 |
| | Lamellar Width.Tail | 0.068 | 0.051 | 0.085 | 0.035 | 0.307 | 0.337 |
| MDD | Lamellar Thickness.Head | -0.006 | 0.831 | 0.922 | 0.003 | 0.905 | 0.905 |
| | Lamellar Thickness.Body | 0.058 | 0.045 | 0.271 | 0.052 | 0.068 | 0.411 |
| | Lamellar Thickness.Tail | -0.048 | 0.104 | 0.313 | -0.007 | 0.803 | 0.905 |
| | Lamellar Width.Head | 0.037 | 0.208 | 0.345 | 0.018 | 0.550 | 0.905 |
| | Lamellar Width.Body | 0.037 | 0.230 | 0.345 | 0.022 | 0.464 | 0.905 |
| | Lamellar Width.Tail | -0.003 | 0.922 | 0.922 | 0.011 | 0.712 | 0.905 |
| SCZ | Lamellar Thickness.Head | 0.155 | 0.000 | 0.000 | 0.113 | 0.001 | 0.000 |
| | Lamellar Thickness.Body | 0.059 | 0.074 | 0.030 | 0.107 | 0.002 | 0.000 |
| | Lamellar Thickness.Tail | -0.020 | 0.552 | 0.184 | -0.005 | 0.880 | 0.147 |
| | Lamellar Width.Head | 0.074 | 0.029 | 0.015 | 0.148 | 0.000 | 0.000 |
| | Lamellar Width.Body | 0.152 | 0.000 | 0.000 | 0.140 | 0.000 | 0.000 |
| | Lamellar Width.Tail | 0.148 | 0.000 | 0.000 | 0.168 | 0.000 | 0.000 |
| MCI | Lamellar Thickness.Head | 0.152 | 0.000 | 0.001 | 0.092 | 0.042 | 0.025 |
| | Lamellar Thickness.Body | 0.194 | 0.000 | 0.000 | 0.227 | 0.000 | 0.000 |
| | Lamellar Thickness.Tail | 0.158 | 0.000 | 0.001 | 0.092 | 0.040 | 0.025 |
| | Lamellar Width.Head | 0.102 | 0.014 | 0.014 | 0.110 | 0.010 | 0.010 |
| | Lamellar Width.Body | 0.169 | 0.000 | 0.000 | 0.185 | 0.000 | 0.000 |
| | Lamellar Width.Tail | 0.117 | 0.004 | 0.005 | 0.081 | 0.059 | 0.029 |
| DM | Lamellar Thickness.Head | 0.181 | 0.000 | 0.000 | 0.156 | 0.000 | 0.000 |
| | Lamellar Thickness.Body | 0.362 | 0.000 | 0.000 | 0.378 | 0.000 | 0.000 |
| | Lamellar Thickness.Tail | 0.219 | 0.000 | 0.000 | 0.180 | 0.000 | 0.000 |
| | Lamellar Width.Head | 0.179 | 0.000 | 0.000 | 0.179 | 0.000 | 0.000 |
| | Lamellar Width.Body | 0.247 | 0.000 | 0.000 | 0.272 | 0.000 | 0.000 |
| | Lamellar Width.Tail | 0.234 | 0.000 | 0.000 | 0.221 | 0.000 | 0.000 |

#### 28. Supplementary Table S16. Cognitive function associations

| | Memory | | | Language | | | | Effective function | | | |
|---|---|---|---|---|---|---|---|---|---|---|---|
| features | Effect size | P | Corrected P | Effect size | P | Corrected P | Effect size | P | Corrected P | Effect size | P |
| Lamellar width 1(Left) | 0.018 | 0.766 | 0.210 | 0.061 | 0.314 | 1.008 | 0.065 | 0.038 | 0.533 | 0.624 | 0.258 |
| Lamellar width 1(Right) | 0.121 | 0.036 | 0.012 | 0.131 | 0.028 | 2.213 | 0.025 | 0.165 | 0.006 | 2.763 | 0.012 |
| Lamellar width 2(Left) | -0.083 | 0.155 | 0.048 | -0.046 | 0.443 | -0.768 | 0.086 | -0.068 | 0.263 | -1.122 | 0.146 |
| Lamellar width 2(Right) | 0.007 | 0.906 | 0.234 | 0.014 | 0.813 | 0.237 | 0.407 | 0.056 | 0.356 | 0.925 | 0.178 |
| Lamellar width 3(Left) | -0.109 | 0.061 | 0.020 | -0.060 | 0.314 | -1.010 | 0.065 | -0.092 | 0.129 | -1.521 | 0.088 |
| Lamellar width 3(Right) | -0.014 | 0.805 | 0.215 | -0.002 | 0.967 | -0.041 | 0.468 | 0.060 | 0.331 | 0.973 | 0.171 |
| Lamellar width 4(Left) | 0.012 | 0.840 | 0.217 | 0.094 | 0.117 | 1.575 | 0.026 | 0.087 | 0.157 | 1.421 | 0.102 |
| Lamellar width 4(Right) | 0.034 | 0.560 | 0.160 | 0.021 | 0.723 | 0.354 | 0.374 | 0.054 | 0.372 | 0.895 | 0.180 |
| Lamellar width 5(Left) | 0.016 | 0.786 | 0.210 | 0.045 | 0.446 | 0.763 | 0.086 | 0.053 | 0.384 | 0.873 | 0.192 |
| Lamellar width 5(Right) | 0.029 | 0.612 | 0.169 | 0.061 | 0.308 | 1.021 | 0.165 | 0.099 | 0.100 | 1.649 | 0.065 |
| Lamellar width 6(Left) | 0.184 | 0.002 | 0.001 | 0.140 | 0.019 | 2.350 | 0.005 | 0.198 | 0.001 | 3.303 | 0.002 |
| Lamellar width 6(Right) | 0.226 | 0.000 | 0.000 | 0.092 | 0.126 | 1.533 | 0.085 | 0.177 | 0.003 | 2.951 | 0.010 |
| Lamellar width 7(Left) | 0.227 | 0.000 | 0.000 | 0.108 | 0.074 | 1.796 | 0.017 | 0.157 | 0.010 | 2.599 | 0.009 |
| Lamellar width 7(Right) | 0.237 | 0.000 | 0.000 | 0.182 | 0.002 | 3.098 | 0.004 | 0.209 | 0.000 | 3.525 | 0.002 |
| Lamellar width 8(Left) | 0.329 | 0.000 | 0.000 | 0.217 | 0.000 | 3.693 | 0.000 | 0.249 | 0.000 | 4.215 | 0.000 |
| Lamellar width 8(Right) | 0.300 | 0.000 | 0.000 | 0.228 | 0.000 | 3.909 | 0.000 | 0.294 | 0.000 | 5.054 | 0.000 |
| Lamellar width 9(Left) | 0.265 | 0.000 | 0.000 | 0.123 | 0.039 | 2.078 | 0.009 | 0.177 | 0.003 | 2.956 | 0.004 |
| Lamellar width 9(Right) | 0.226 | 0.000 | 0.000 | 0.143 | 0.016 | 2.432 | 0.016 | 0.201 | 0.001 | 3.390 | 0.003 |
| Lamellar width 10(Left) | 0.288 | 0.000 | 0.000 | 0.136 | 0.023 | 2.295 | 0.006 | 0.186 | 0.002 | 3.113 | 0.003 |
| Lamellar width 10(Right) | 0.216 | 0.000 | 0.000 | 0.105 | 0.078 | 1.771 | 0.058 | 0.152 | 0.012 | 2.531 | 0.013 |
| Lamellar width 11(Left) | 0.290 | 0.000 | 0.000 | 0.208 | 0.000 | 3.536 | 0.000 | 0.234 | 0.000 | 3.927 | 0.000 |
| Lamellar width 11(Right) | 0.220 | 0.000 | 0.000 | 0.070 | 0.243 | 1.170 | 0.135 | 0.110 | 0.068 | 1.834 | 0.053 |
| Lamellar width 12(Left) | 0.337 | 0.000 | 0.000 | 0.204 | 0.001 | 3.470 | 0.000 | 0.185 | 0.002 | 3.078 | 0.003 |
| Lamellar width 12(Right) | 0.256 | 0.000 | 0.000 | 0.154 | 0.009 | 2.627 | 0.011 | 0.153 | 0.011 | 2.554 | 0.013 |
| Lamellar width 13(Left) | 0.323 | 0.000 | 0.000 | 0.229 | 0.000 | 3.915 | 0.000 | 0.209 | 0.001 | 3.491 | 0.001 |
| Lamellar width 13(Right) | 0.250 | 0.000 | 0.000 | 0.087 | 0.142 | 1.473 | 0.089 | 0.093 | 0.123 | 1.547 | 0.077 |

| | | | | | | | | | | | |
|---|---|---|---|---|---|---|---|---|---|---|---|
| Lamellar width 14(Left) | 0.294 | 0.000 | 0.000 | 0.170 | 0.004 | 2.883 | 0.002 | 0.128 | 0.035 | 2.125 | 0.030 |
| Lamellar width 14(Right) | 0.230 | 0.000 | 0.000 | 0.080 | 0.180 | 1.346 | 0.104 | 0.074 | 0.221 | 1.227 | 0.128 |
| Lamellar width 15(Left) | 0.290 | 0.000 | 0.000 | 0.155 | 0.009 | 2.630 | 0.003 | 0.116 | 0.054 | 1.935 | 0.041 |
| Lamellar width 15(Right) | 0.210 | 0.000 | 0.000 | 0.098 | 0.099 | 1.655 | 0.071 | 0.062 | 0.308 | 1.021 | 0.165 |
| Lamellar width 16(Left) | 0.278 | 0.000 | 0.000 | 0.175 | 0.003 | 2.980 | 0.001 | 0.082 | 0.177 | 1.353 | 0.111 |
| Lamellar width 16(Right) | 0.200 | 0.000 | 0.000 | 0.113 | 0.058 | 1.904 | 0.046 | 0.062 | 0.304 | 1.029 | 0.165 |
| Lamellar width 17(Left) | 0.257 | 0.000 | 0.000 | 0.147 | 0.013 | 2.490 | 0.004 | 0.054 | 0.374 | 0.890 | 0.192 |
| Lamellar width 17(Right) | 0.175 | 0.002 | 0.001 | 0.161 | 0.006 | 2.748 | 0.009 | 0.099 | 0.100 | 1.649 | 0.065 |
| Lamellar width 18(Left) | 0.254 | 0.000 | 0.000 | 0.154 | 0.010 | 2.598 | 0.003 | 0.058 | 0.346 | 0.945 | 0.185 |
| Lamellar width 18(Right) | 0.169 | 0.003 | 0.001 | 0.186 | 0.002 | 3.179 | 0.003 | 0.143 | 0.017 | 2.392 | 0.017 |
| Lamellar width 19(Left) | 0.272 | 0.000 | 0.000 | 0.220 | 0.000 | 3.774 | 0.000 | 0.126 | 0.038 | 2.081 | 0.032 |
| Lamellar width 19(Right) | 0.133 | 0.021 | 0.008 | 0.137 | 0.021 | 2.327 | 0.019 | 0.173 | 0.004 | 2.907 | 0.010 |
| Lamellar width 20(Left) | 0.357 | 0.000 | 0.000 | 0.306 | 0.000 | 5.389 | 0.000 | 0.210 | 0.000 | 3.535 | 0.001 |
| Lamellar width 20(Right) | 0.252 | 0.000 | 0.000 | 0.243 | 0.000 | 4.208 | 0.000 | 0.259 | 0.000 | 4.443 | 0.000 |
| Lamellar width 21(Left) | 0.262 | 0.000 | 0.000 | 0.253 | 0.000 | 4.382 | 0.000 | 0.165 | 0.006 | 2.755 | 0.006 |
| Lamellar width 21(Right) | 0.110 | 0.057 | 0.019 | 0.144 | 0.016 | 2.436 | 0.016 | 0.151 | 0.012 | 2.524 | 0.013 |
| Lamellar width 22(Left) | 0.215 | 0.000 | 0.000 | 0.222 | 0.000 | 3.837 | 0.000 | 0.173 | 0.004 | 2.901 | 0.004 |
| Lamellar width 22(Right) | 0.068 | 0.238 | 0.070 | 0.123 | 0.038 | 2.087 | 0.032 | 0.109 | 0.070 | 1.818 | 0.053 |
| Lamellar width 23(Left) | 0.208 | 0.000 | 0.000 | 0.223 | 0.000 | 3.838 | 0.000 | 0.186 | 0.002 | 3.117 | 0.003 |
| Lamellar width 23(Right) | 0.083 | 0.150 | 0.046 | 0.090 | 0.130 | 1.519 | 0.085 | 0.082 | 0.176 | 1.357 | 0.106 |
| Lamellar width 24(Left) | 0.285 | 0.000 | 0.000 | 0.263 | 0.000 | 4.598 | 0.000 | 0.242 | 0.000 | 4.118 | 0.000 |
| Lamellar width 24(Right) | 0.179 | 0.002 | 0.001 | 0.157 | 0.008 | 2.663 | 0.010 | 0.115 | 0.057 | 1.913 | 0.047 |
| Lamellar width 25(Left) | 0.285 | 0.000 | 0.000 | 0.318 | 0.000 | 5.658 | 0.000 | 0.255 | 0.000 | 4.368 | 0.000 |
| Lamellar width 25(Right) | 0.202 | 0.000 | 0.000 | 0.250 | 0.000 | 4.332 | 0.000 | 0.136 | 0.024 | 2.266 | 0.023 |
| Lamellar width 26(Left) | 0.208 | 0.000 | 0.000 | 0.263 | 0.000 | 4.588 | 0.000 | 0.186 | 0.002 | 3.122 | 0.003 |
| Lamellar width 26(Right) | 0.198 | 0.001 | 0.000 | 0.252 | 0.000 | 4.375 | 0.000 | 0.159 | 0.009 | 2.646 | 0.013 |
| Lamellar width 27(Left) | 0.143 | 0.013 | 0.004 | 0.198 | 0.001 | 3.394 | 0.000 | 0.123 | 0.042 | 2.044 | 0.033 |
| Lamellar width 27(Right) | 0.190 | 0.001 | 0.000 | 0.236 | 0.000 | 4.092 | 0.000 | 0.150 | 0.012 | 2.515 | 0.013 |
| Lamellar width 28(Left) | 0.244 | 0.000 | 0.000 | 0.278 | 0.000 | 4.866 | 0.000 | 0.216 | 0.000 | 3.649 | 0.001 |
| Lamellar width 28(Right) | 0.173 | 0.003 | 0.001 | 0.227 | 0.000 | 3.917 | 0.000 | 0.130 | 0.031 | 2.169 | 0.027 |
| Lamellar width 29(Left) | 0.090 | 0.118 | 0.038 | 0.174 | 0.003 | 2.959 | 0.001 | 0.093 | 0.125 | 1.540 | 0.088 |
| Lamellar width 29(Right) | 0.153 | 0.008 | 0.003 | 0.202 | 0.001 | 3.474 | 0.001 | 0.156 | 0.010 | 2.607 | 0.013 |

| | | | | | | | | | | | |
|---|---|---|---|---|---|---|---|---|---|---|---|
| Lamellar width 30(Left) | 0.079 | 0.175 | 0.052 | 0.158 | 0.008 | 2.677 | 0.003 | 0.079 | 0.193 | 1.304 | 0.116 |
| Lamellar width 30(Right) | 0.145 | 0.012 | 0.005 | 0.179 | 0.002 | 3.063 | 0.004 | 0.164 | 0.006 | 2.757 | 0.012 |
| Lamellar width 31(Left) | 0.054 | 0.357 | 0.102 | 0.133 | 0.026 | 2.245 | 0.006 | 0.068 | 0.262 | 1.125 | 0.146 |
| Lamellar width 31(Right) | 0.085 | 0.141 | 0.045 | 0.084 | 0.157 | 1.418 | 0.094 | 0.104 | 0.084 | 1.736 | 0.060 |
| combined_label_Length(Left) | 0.369 | 0.000 | 0.000 | 0.294 | 0.000 | 5.182 | 0.000 | 0.271 | 0.000 | 4.649 | 0.000 |
| combined_label_Length(Right) | 0.303 | 0.000 | 0.000 | 0.203 | 0.001 | 3.494 | 0.001 | 0.264 | 0.000 | 4.527 | 0.000 |
| Lamellar thickness 2(Left) | -0.053 | 0.367 | 0.511 | 0.023 | 0.707 | 0.376 | 0.797 | -0.058 | 0.344 | -0.949 | 0.635 |
| Lamellar thickness 2(Right) | -0.026 | 0.660 | 0.517 | -0.024 | 0.702 | -0.383 | 0.624 | 0.048 | 0.446 | 0.764 | 0.575 |
| Lamellar thickness 6(Left) | 0.053 | 0.362 | 0.511 | -0.002 | 0.974 | -0.033 | 0.930 | 0.016 | 0.787 | 0.271 | 0.867 |
| Lamellar thickness 6(Right) | 0.015 | 0.799 | 0.584 | 0.071 | 0.238 | 1.182 | 0.306 | 0.040 | 0.515 | 0.652 | 0.635 |
| Lamellar thickness 10(Left) | 0.080 | 0.168 | 0.348 | 0.058 | 0.331 | 0.975 | 0.585 | 0.068 | 0.264 | 1.120 | 0.586 |
| Lamellar thickness 10(Right) | 0.058 | 0.319 | 0.286 | 0.067 | 0.264 | 1.120 | 0.322 | 0.061 | 0.318 | 1.001 | 0.463 |
| Lamellar thickness 14(Left) | -0.004 | 0.950 | 0.835 | 0.050 | 0.398 | 0.846 | 0.603 | 0.026 | 0.665 | 0.434 | 0.833 |
| Lamellar thickness 14(Right) | 0.059 | 0.313 | 0.286 | 0.006 | 0.920 | 0.100 | 0.750 | 0.049 | 0.423 | 0.803 | 0.552 |
| Lamellar thickness 18(Left) | 0.110 | 0.056 | 0.201 | 0.100 | 0.093 | 1.685 | 0.275 | 0.028 | 0.648 | 0.457 | 0.833 |
| Lamellar thickness 18(Right) | 0.081 | 0.163 | 0.187 | 0.087 | 0.146 | 1.459 | 0.221 | 0.053 | 0.378 | 0.883 | 0.511 |
| Lamellar thickness 22(Left) | 0.102 | 0.079 | 0.246 | 0.102 | 0.087 | 1.717 | 0.272 | 0.029 | 0.636 | 0.474 | 0.833 |
| Lamellar thickness 22(Right) | 0.208 | 0.000 | 0.003 | 0.126 | 0.034 | 2.134 | 0.074 | 0.128 | 0.034 | 2.135 | 0.133 |
| Lamellar thickness 26(Left) | 0.149 | 0.011 | 0.064 | 0.087 | 0.151 | 1.441 | 0.390 | 0.072 | 0.246 | 1.164 | 0.586 |
| Lamellar thickness 26(Right) | 0.208 | 0.000 | 0.003 | 0.175 | 0.003 | 2.992 | 0.019 | 0.101 | 0.094 | 1.681 | 0.199 |
| Lamellar thickness 30(Left) | 0.174 | 0.003 | 0.023 | 0.121 | 0.046 | 2.008 | 0.201 | 0.084 | 0.174 | 1.362 | 0.563 |
| Lamellar thickness 30(Right) | 0.228 | 0.000 | 0.002 | 0.167 | 0.005 | 2.843 | 0.023 | 0.121 | 0.046 | 2.005 | 0.165 |
| Lamellar thickness 34(Left) | 0.173 | 0.003 | 0.023 | 0.101 | 0.093 | 1.684 | 0.275 | 0.062 | 0.312 | 1.013 | 0.631 |
| Lamellar thickness 34(Right) | 0.203 | 0.000 | 0.003 | 0.127 | 0.032 | 2.151 | 0.073 | 0.108 | 0.073 | 1.799 | 0.185 |
| Lamellar thickness 38(Left) | 0.170 | 0.004 | 0.028 | 0.102 | 0.097 | 1.668 | 0.277 | 0.048 | 0.440 | 0.773 | 0.717 |
| Lamellar thickness 38(Right) | 0.197 | 0.001 | 0.004 | 0.149 | 0.012 | 2.534 | 0.043 | 0.064 | 0.293 | 1.053 | 0.450 |
| Lamellar thickness 42(Left) | 0.211 | 0.000 | 0.005 | 0.119 | 0.052 | 1.948 | 0.203 | 0.073 | 0.243 | 1.170 | 0.586 |
| Lamellar thickness 42(Right) | 0.204 | 0.000 | 0.003 | 0.158 | 0.008 | 2.683 | 0.030 | 0.082 | 0.176 | 1.356 | 0.305 |
| Lamellar thickness 46(Left) | 0.230 | 0.000 | 0.001 | 0.169 | 0.005 | 2.810 | 0.070 | 0.071 | 0.248 | 1.158 | 0.586 |
| Lamellar thickness 46(Right) | 0.201 | 0.000 | 0.004 | 0.174 | 0.003 | 2.975 | 0.019 | 0.071 | 0.242 | 1.172 | 0.396 |
| Lamellar thickness 50(Left) | 0.237 | 0.000 | 0.001 | 0.176 | 0.003 | 2.969 | 0.057 | 0.046 | 0.450 | 0.757 | 0.717 |
| Lamellar thickness 50(Right) | 0.206 | 0.000 | 0.003 | 0.215 | 0.000 | 3.702 | 0.005 | 0.130 | 0.031 | 2.167 | 0.133 |

| | | | | | | | | | | | |
|---|---|---|---|---|---|---|---|---|---|---|---|
| Lamellar thickness 54(Left) | 0.268 | 0.000 | 0.000 | 0.215 | 0.000 | 3.632 | 0.009 | 0.086 | 0.162 | 1.401 | 0.563 |
| Lamellar thickness 54(Right) | 0.227 | 0.000 | 0.002 | 0.233 | 0.000 | 4.033 | 0.003 | 0.118 | 0.051 | 1.960 | 0.172 |
| Lamellar thickness 58(Left) | 0.313 | 0.000 | 0.000 | 0.252 | 0.000 | 4.285 | 0.003 | 0.146 | 0.018 | 2.389 | 0.223 |
| Lamellar thickness 58(Right) | 0.180 | 0.002 | 0.007 | 0.216 | 0.000 | 3.720 | 0.005 | 0.102 | 0.090 | 1.700 | 0.199 |
| Lamellar thickness 62(Left) | 0.260 | 0.000 | 0.000 | 0.216 | 0.000 | 3.656 | 0.009 | 0.112 | 0.068 | 1.831 | 0.413 |
| Lamellar thickness 62(Right) | 0.160 | 0.006 | 0.013 | 0.171 | 0.004 | 2.916 | 0.021 | 0.026 | 0.664 | 0.434 | 0.745 |
| Lamellar thickness 66(Left) | 0.192 | 0.001 | 0.010 | 0.216 | 0.000 | 3.650 | 0.009 | 0.082 | 0.181 | 1.342 | 0.563 |
| Lamellar thickness 66(Right) | 0.141 | 0.015 | 0.032 | 0.143 | 0.017 | 2.409 | 0.052 | 0.000 | 0.998 | 0.002 | 0.998 |
| Lamellar thickness 70(Left) | 0.172 | 0.003 | 0.023 | 0.171 | 0.004 | 2.921 | 0.057 | 0.098 | 0.105 | 1.628 | 0.497 |
| Lamellar thickness 70(Right) | 0.110 | 0.056 | 0.084 | 0.098 | 0.101 | 1.647 | 0.167 | 0.039 | 0.524 | 0.638 | 0.639 |
| Lamellar thickness 74(Left) | 0.054 | 0.355 | 0.511 | 0.115 | 0.052 | 1.949 | 0.203 | 0.099 | 0.100 | 1.651 | 0.497 |
| Lamellar thickness 74(Right) | 0.064 | 0.275 | 0.261 | 0.079 | 0.189 | 1.318 | 0.268 | 0.068 | 0.266 | 1.115 | 0.422 |
| Lamellar thickness 78(Left) | 0.008 | 0.896 | 0.827 | 0.058 | 0.331 | 0.974 | 0.585 | 0.082 | 0.173 | 1.367 | 0.563 |
| Lamellar thickness 78(Right) | 0.072 | 0.212 | 0.224 | 0.064 | 0.281 | 1.079 | 0.335 | 0.027 | 0.660 | 0.440 | 0.745 |
| Lamellar thickness 82(Left) | -0.010 | 0.856 | 0.822 | -0.004 | 0.944 | -0.071 | 0.918 | 0.031 | 0.609 | 0.512 | 0.833 |
| Lamellar thickness 82(Right) | 0.074 | 0.200 | 0.214 | 0.065 | 0.273 | 1.097 | 0.330 | 0.042 | 0.493 | 0.687 | 0.621 |
| Lamellar thickness 86(Left) | 0.022 | 0.702 | 0.748 | 0.041 | 0.492 | 0.688 | 0.666 | 0.049 | 0.418 | 0.811 | 0.712 |
| Lamellar thickness 86(Right) | 0.010 | 0.860 | 0.611 | 0.041 | 0.490 | 0.691 | 0.485 | 0.022 | 0.716 | 0.364 | 0.772 |
| Lamellar thickness 90(Left) | 0.024 | 0.680 | 0.733 | 0.030 | 0.617 | 0.501 | 0.743 | 0.023 | 0.704 | 0.380 | 0.859 |
| Lamellar thickness 90(Right) | -0.063 | 0.280 | 0.261 | 0.003 | 0.965 | 0.044 | 0.779 | -0.031 | 0.604 | -0.519 | 0.717 |
| Lamellar thickness 94(Left) | 0.039 | 0.507 | 0.631 | 0.093 | 0.124 | 1.545 | 0.345 | 0.045 | 0.465 | 0.732 | 0.717 |
| Lamellar thickness 94(Right) | -0.024 | 0.676 | 0.519 | 0.069 | 0.246 | 1.162 | 0.306 | 0.052 | 0.395 | 0.853 | 0.521 |
| Lamellar thickness 98(Left) | 0.080 | 0.172 | 0.348 | 0.116 | 0.054 | 1.938 | 0.203 | 0.095 | 0.117 | 1.573 | 0.531 |
| Lamellar thickness 98(Right) | -0.030 | 0.602 | 0.481 | 0.040 | 0.501 | 0.674 | 0.490 | 0.040 | 0.513 | 0.655 | 0.635 |
| Lamellar thickness 106(Left) | -0.009 | 0.878 | 0.826 | 0.054 | 0.371 | 0.897 | 0.603 | 0.041 | 0.503 | 0.671 | 0.749 |
| Lamellar thickness 106(Right) | 0.081 | 0.165 | 0.187 | 0.105 | 0.079 | 1.761 | 0.140 | 0.119 | 0.049 | 1.974 | 0.171 |
| Lamellar thickness 110(Left) | 0.041 | 0.482 | 0.617 | 0.014 | 0.821 | 0.227 | 0.870 | 0.044 | 0.467 | 0.729 | 0.717 |
| Lamellar thickness 110(Right) | 0.039 | 0.505 | 0.427 | 0.034 | 0.565 | 0.576 | 0.535 | 0.066 | 0.276 | 1.092 | 0.431 |
| Lamellar thickness 109(Left) | 0.028 | 0.624 | 0.680 | 0.004 | 0.940 | 0.075 | 0.918 | 0.078 | 0.196 | 1.295 | 0.564 |
| Lamellar thickness 109(Right) | 0.058 | 0.316 | 0.286 | 0.009 | 0.875 | 0.157 | 0.726 | 0.071 | 0.243 | 1.170 | 0.396 |
| Lamellar thickness 105(Left) | 0.069 | 0.235 | 0.433 | 0.003 | 0.958 | 0.052 | 0.923 | 0.063 | 0.302 | 1.035 | 0.631 |
| Lamellar thickness 105(Right) | 0.048 | 0.408 | 0.357 | 0.000 | 0.996 | 0.005 | 0.792 | 0.087 | 0.150 | 1.445 | 0.270 |

| | | | | | | | | | | | |
|---|---|---|---|---|---|---|---|---|---|---|---|
| Lamellar thickness 101(Left) | -0.003 | 0.962 | 0.835 | -0.029 | 0.624 | -0.490 | 0.744 | 0.002 | 0.971 | 0.037 | 0.970 |
| Lamellar thickness 101(Right) | 0.070 | 0.225 | 0.231 | 0.050 | 0.401 | 0.842 | 0.415 | 0.104 | 0.086 | 1.725 | 0.199 |
| Lamellar thickness 97(Left) | -0.003 | 0.965 | 0.835 | -0.036 | 0.552 | -0.595 | 0.705 | 0.023 | 0.702 | 0.383 | 0.859 |
| Lamellar thickness 97(Right) | 0.109 | 0.058 | 0.085 | 0.085 | 0.153 | 1.432 | 0.229 | 0.135 | 0.025 | 2.253 | 0.131 |
| Lamellar thickness 93(Left) | 0.011 | 0.855 | 0.822 | -0.025 | 0.672 | -0.424 | 0.783 | -0.001 | 0.987 | -0.016 | 0.970 |
| Lamellar thickness 93(Right) | 0.095 | 0.101 | 0.132 | 0.076 | 0.203 | 1.277 | 0.282 | 0.120 | 0.046 | 2.004 | 0.165 |
| Lamellar thickness 89(Left) | 0.047 | 0.420 | 0.552 | 0.016 | 0.793 | 0.262 | 0.852 | 0.033 | 0.586 | 0.545 | 0.819 |
| Lamellar thickness 89(Right) | 0.062 | 0.281 | 0.261 | 0.084 | 0.157 | 1.420 | 0.230 | 0.102 | 0.090 | 1.700 | 0.199 |
| Lamellar thickness 85(Left) | 0.058 | 0.318 | 0.511 | 0.021 | 0.729 | 0.346 | 0.814 | 0.056 | 0.353 | 0.930 | 0.642 |
| Lamellar thickness 85(Right) | 0.063 | 0.272 | 0.261 | 0.069 | 0.247 | 1.160 | 0.306 | 0.112 | 0.063 | 1.867 | 0.185 |
| Lamellar thickness 81(Left) | 0.019 | 0.739 | 0.751 | 0.028 | 0.641 | 0.467 | 0.755 | 0.082 | 0.177 | 1.353 | 0.563 |
| Lamellar thickness 81(Right) | 0.071 | 0.223 | 0.231 | 0.104 | 0.081 | 1.751 | 0.140 | 0.122 | 0.044 | 2.024 | 0.165 |
| Lamellar thickness 77(Left) | 0.108 | 0.062 | 0.204 | 0.103 | 0.082 | 1.744 | 0.264 | 0.160 | 0.008 | 2.687 | 0.169 |
| Lamellar thickness 77(Right) | 0.090 | 0.121 | 0.151 | 0.054 | 0.367 | 0.904 | 0.394 | 0.108 | 0.073 | 1.799 | 0.185 |
| Lamellar thickness 73(Left) | 0.090 | 0.118 | 0.277 | 0.082 | 0.169 | 1.378 | 0.412 | 0.151 | 0.012 | 2.530 | 0.218 |
| Lamellar thickness 73(Right) | 0.114 | 0.050 | 0.079 | 0.101 | 0.092 | 1.691 | 0.156 | 0.130 | 0.031 | 2.167 | 0.133 |
| Lamellar thickness 69(Left) | 0.120 | 0.037 | 0.164 | 0.116 | 0.050 | 1.969 | 0.203 | 0.147 | 0.015 | 2.453 | 0.223 |
| Lamellar thickness 69(Right) | 0.172 | 0.003 | 0.009 | 0.148 | 0.013 | 2.495 | 0.046 | 0.165 | 0.006 | 2.756 | 0.098 |
| Lamellar thickness 65(Left) | 0.134 | 0.021 | 0.114 | 0.128 | 0.032 | 2.151 | 0.156 | 0.193 | 0.001 | 3.223 | 0.052 |
| Lamellar thickness 65(Right) | 0.181 | 0.002 | 0.007 | 0.181 | 0.003 | 3.045 | 0.019 | 0.130 | 0.033 | 2.142 | 0.133 |
| Lamellar thickness 61(Left) | 0.096 | 0.100 | 0.258 | 0.152 | 0.011 | 2.569 | 0.095 | 0.162 | 0.008 | 2.683 | 0.169 |
| Lamellar thickness 61(Right) | 0.195 | 0.001 | 0.004 | 0.210 | 0.000 | 3.547 | 0.006 | 0.111 | 0.071 | 1.813 | 0.185 |
| Lamellar thickness 57(Left) | 0.098 | 0.091 | 0.249 | 0.153 | 0.010 | 2.590 | 0.095 | 0.139 | 0.022 | 2.307 | 0.230 |
| Lamellar thickness 57(Right) | 0.128 | 0.030 | 0.055 | 0.169 | 0.005 | 2.818 | 0.023 | 0.091 | 0.142 | 1.471 | 0.264 |
| Lamellar thickness 53(Left) | 0.059 | 0.306 | 0.507 | 0.142 | 0.017 | 2.399 | 0.114 | 0.110 | 0.068 | 1.831 | 0.413 |
| Lamellar thickness 53(Right) | 0.184 | 0.002 | 0.007 | 0.206 | 0.001 | 3.479 | 0.006 | 0.139 | 0.023 | 2.286 | 0.131 |
| Lamellar thickness 49(Left) | 0.012 | 0.832 | 0.815 | 0.112 | 0.062 | 1.872 | 0.213 | 0.100 | 0.101 | 1.646 | 0.497 |
| Lamellar thickness 49(Right) | 0.141 | 0.015 | 0.032 | 0.168 | 0.005 | 2.852 | 0.023 | 0.061 | 0.319 | 0.998 | 0.463 |
| Lamellar thickness 45(Left) | 0.010 | 0.868 | 0.825 | 0.073 | 0.225 | 1.215 | 0.460 | 0.015 | 0.809 | 0.242 | 0.880 |
| Lamellar thickness 45(Right) | 0.190 | 0.001 | 0.004 | 0.190 | 0.001 | 3.262 | 0.012 | 0.101 | 0.094 | 1.682 | 0.199 |
| Lamellar thickness 41(Left) | 0.030 | 0.610 | 0.680 | 0.042 | 0.493 | 0.686 | 0.666 | 0.051 | 0.410 | 0.825 | 0.712 |
| Lamellar thickness 41(Right) | 0.192 | 0.001 | 0.004 | 0.215 | 0.000 | 3.692 | 0.005 | 0.111 | 0.067 | 1.837 | 0.185 |

| | | | | | | | | | | | |
|---|---|---|---|---|---|---|---|---|---|---|---|
| Lamellar thickness 37(Left) | 0.102 | 0.082 | 0.246 | 0.144 | 0.016 | 2.423 | 0.114 | 0.072 | 0.240 | 1.179 | 0.586 |
| Lamellar thickness 37(Right) | 0.194 | 0.001 | 0.004 | 0.276 | 0.000 | 4.818 | 0.000 | 0.142 | 0.018 | 2.372 | 0.128 |
| Lamellar thickness 33(Left) | 0.050 | 0.397 | 0.536 | 0.078 | 0.195 | 1.299 | 0.431 | 0.036 | 0.557 | 0.588 | 0.799 |
| Lamellar thickness 33(Right) | 0.137 | 0.018 | 0.036 | 0.209 | 0.000 | 3.589 | 0.006 | 0.134 | 0.026 | 2.240 | 0.131 |
| Lamellar thickness 29(Left) | 0.077 | 0.187 | 0.359 | 0.073 | 0.224 | 1.219 | 0.460 | 0.078 | 0.202 | 1.279 | 0.564 |
| Lamellar thickness 29(Right) | 0.218 | 0.000 | 0.003 | 0.175 | 0.003 | 2.979 | 0.019 | 0.147 | 0.015 | 2.446 | 0.112 |
| Lamellar thickness 25(Left) | 0.092 | 0.113 | 0.277 | 0.057 | 0.340 | 0.957 | 0.590 | 0.075 | 0.215 | 1.242 | 0.573 |
| Lamellar thickness 25(Right) | 0.194 | 0.001 | 0.004 | 0.128 | 0.031 | 2.168 | 0.072 | 0.101 | 0.095 | 1.677 | 0.199 |
| Lamellar thickness 21(Left) | 0.031 | 0.596 | 0.680 | 0.051 | 0.395 | 0.852 | 0.603 | 0.040 | 0.513 | 0.655 | 0.749 |
| Lamellar thickness 21(Right) | 0.188 | 0.001 | 0.005 | 0.160 | 0.007 | 2.698 | 0.030 | 0.111 | 0.068 | 1.835 | 0.185 |
| Lamellar thickness 17(Left) | 0.004 | 0.949 | 0.835 | 0.051 | 0.392 | 0.858 | 0.603 | 0.026 | 0.664 | 0.435 | 0.833 |
| Lamellar thickness 17(Right) | 0.166 | 0.004 | 0.011 | 0.176 | 0.003 | 2.991 | 0.019 | 0.137 | 0.024 | 2.276 | 0.131 |
| Lamellar thickness 13(Left) | -0.036 | 0.533 | 0.648 | 0.005 | 0.939 | 0.076 | 0.918 | -0.058 | 0.336 | -0.965 | 0.631 |
| Lamellar thickness 13(Right) | 0.165 | 0.004 | 0.011 | 0.165 | 0.006 | 2.795 | 0.023 | 0.156 | 0.010 | 2.593 | 0.098 |
| Lamellar thickness 9(Left) | 0.055 | 0.342 | 0.511 | 0.079 | 0.189 | 1.316 | 0.427 | 0.022 | 0.713 | 0.369 | 0.859 |
| Lamellar thickness 9(Right) | 0.122 | 0.035 | 0.060 | 0.120 | 0.044 | 2.027 | 0.089 | 0.155 | 0.010 | 2.597 | 0.098 |
| Lamellar thickness 5(Left) | 0.019 | 0.740 | 0.751 | 0.088 | 0.145 | 1.463 | 0.383 | 0.084 | 0.171 | 1.372 | 0.563 |
| Lamellar thickness 5(Right) | 0.111 | 0.056 | 0.084 | 0.058 | 0.336 | 0.964 | 0.374 | 0.088 | 0.151 | 1.440 | 0.270 |
| Lamellar thickness 1(Left) | 0.083 | 0.163 | 0.348 | 0.132 | 0.030 | 2.177 | 0.153 | 0.046 | 0.465 | 0.732 | 0.717 |
| Lamellar thickness 1(Right) | 0.164 | 0.005 | 0.012 | 0.141 | 0.019 | 2.351 | 0.057 | 0.174 | 0.004 | 2.871 | 0.098 |
| Lamellar thickness 4(Left) | 0.060 | 0.300 | 0.506 | 0.051 | 0.391 | 0.860 | 0.603 | 0.044 | 0.463 | 0.735 | 0.717 |
| Lamellar thickness 4(Right) | 0.023 | 0.695 | 0.528 | 0.024 | 0.692 | 0.396 | 0.624 | 0.103 | 0.090 | 1.702 | 0.199 |
| Lamellar thickness 8(Left) | 0.196 | 0.001 | 0.008 | 0.131 | 0.029 | 2.189 | 0.153 | 0.211 | 0.000 | 3.531 | 0.027 |
| Lamellar thickness 8(Right) | 0.102 | 0.079 | 0.111 | 0.079 | 0.188 | 1.320 | 0.268 | 0.060 | 0.321 | 0.994 | 0.463 |
| Lamellar thickness 12(Left) | 0.300 | 0.000 | 0.000 | 0.184 | 0.002 | 3.147 | 0.039 | 0.211 | 0.000 | 3.569 | 0.027 |
| Lamellar thickness 12(Right) | 0.150 | 0.010 | 0.024 | 0.108 | 0.074 | 1.797 | 0.132 | 0.101 | 0.098 | 1.659 | 0.202 |
| Lamellar thickness 16(Left) | 0.214 | 0.000 | 0.003 | 0.126 | 0.034 | 2.129 | 0.158 | 0.131 | 0.031 | 2.174 | 0.238 |
| Lamellar thickness 16(Right) | 0.097 | 0.097 | 0.131 | 0.076 | 0.205 | 1.271 | 0.282 | 0.064 | 0.296 | 1.047 | 0.450 |
| Lamellar thickness 20(Left) | 0.147 | 0.010 | 0.063 | 0.113 | 0.057 | 1.910 | 0.209 | 0.122 | 0.044 | 2.026 | 0.298 |
| Lamellar thickness 20(Right) | 0.053 | 0.366 | 0.324 | 0.000 | 0.998 | -0.002 | 0.792 | 0.003 | 0.958 | 0.052 | 0.976 |
| Lamellar thickness 24(Left) | 0.069 | 0.243 | 0.441 | 0.048 | 0.433 | 0.785 | 0.629 | 0.043 | 0.484 | 0.701 | 0.732 |
| Lamellar thickness 24(Right) | 0.019 | 0.746 | 0.558 | -0.018 | 0.768 | -0.296 | 0.650 | 0.026 | 0.671 | 0.425 | 0.745 |

| | | | | | | | | | | | |
|---|---|---|---|---|---|---|---|---|---|---|---|
| Lamellar thickness 28(Left) | 0.131 | 0.023 | 0.117 | 0.032 | 0.588 | 0.543 | 0.733 | 0.069 | 0.256 | 1.138 | 0.586 |
| Lamellar thickness 28(Right) | 0.013 | 0.821 | 0.594 | -0.024 | 0.691 | -0.398 | 0.624 | -0.008 | 0.894 | -0.133 | 0.928 |
| Lamellar thickness 32(Left) | 0.128 | 0.027 | 0.127 | 0.042 | 0.486 | 0.697 | 0.666 | 0.087 | 0.150 | 1.445 | 0.563 |
| Lamellar thickness 32(Right) | 0.036 | 0.535 | 0.446 | -0.032 | 0.597 | -0.530 | 0.559 | 0.025 | 0.681 | 0.412 | 0.748 |
| Lamellar thickness 36(Left) | 0.090 | 0.122 | 0.280 | 0.069 | 0.253 | 1.145 | 0.497 | 0.066 | 0.279 | 1.085 | 0.596 |
| Lamellar thickness 36(Right) | 0.003 | 0.961 | 0.658 | -0.070 | 0.241 | -1.175 | 0.306 | -0.029 | 0.632 | -0.479 | 0.738 |
| Lamellar thickness 40(Left) | 0.049 | 0.407 | 0.543 | 0.052 | 0.391 | 0.859 | 0.603 | 0.028 | 0.649 | 0.455 | 0.833 |
| Lamellar thickness 40(Right) | -0.024 | 0.676 | 0.519 | -0.022 | 0.708 | -0.374 | 0.624 | 0.026 | 0.663 | 0.436 | 0.745 |
| Lamellar thickness 44(Left) | 0.097 | 0.099 | 0.258 | 0.065 | 0.284 | 1.073 | 0.538 | 0.080 | 0.193 | 1.304 | 0.564 |
| Lamellar thickness 44(Right) | -0.040 | 0.490 | 0.418 | -0.074 | 0.212 | -1.250 | 0.288 | -0.015 | 0.803 | -0.249 | 0.849 |
| Lamellar thickness 48(Left) | 0.099 | 0.089 | 0.249 | 0.089 | 0.139 | 1.486 | 0.376 | 0.083 | 0.174 | 1.364 | 0.563 |
| Lamellar thickness 48(Right) | -0.035 | 0.550 | 0.447 | -0.095 | 0.111 | -1.600 | 0.178 | -0.024 | 0.696 | -0.392 | 0.757 |
| Lamellar thickness 52(Left) | 0.052 | 0.373 | 0.512 | 0.041 | 0.496 | 0.681 | 0.666 | 0.030 | 0.623 | 0.492 | 0.833 |
| Lamellar thickness 52(Right) | -0.008 | 0.886 | 0.624 | -0.038 | 0.520 | -0.643 | 0.503 | -0.018 | 0.764 | -0.300 | 0.816 |
| Lamellar thickness 56(Left) | -0.065 | 0.262 | 0.466 | -0.080 | 0.183 | -1.336 | 0.427 | -0.105 | 0.084 | -1.732 | 0.485 |
| Lamellar thickness 56(Right) | 0.097 | 0.094 | 0.130 | 0.060 | 0.316 | 1.005 | 0.361 | 0.110 | 0.068 | 1.831 | 0.185 |
| Lamellar thickness 60(Left) | 0.034 | 0.562 | 0.661 | 0.031 | 0.607 | 0.516 | 0.743 | 0.017 | 0.783 | 0.276 | 0.867 |
| Lamellar thickness 60(Right) | 0.019 | 0.748 | 0.558 | 0.050 | 0.400 | 0.843 | 0.415 | 0.077 | 0.203 | 1.277 | 0.346 |
| Lamellar thickness 64(Left) | -0.008 | 0.887 | 0.826 | -0.011 | 0.856 | -0.181 | 0.873 | 0.029 | 0.639 | 0.469 | 0.833 |
| Lamellar thickness 64(Right) | 0.085 | 0.142 | 0.172 | 0.057 | 0.344 | 0.948 | 0.379 | 0.092 | 0.130 | 1.520 | 0.248 |
| Lamellar thickness 68(Left) | -0.058 | 0.320 | 0.511 | -0.063 | 0.295 | -1.049 | 0.549 | -0.022 | 0.717 | -0.362 | 0.859 |
| Lamellar thickness 68(Right) | 0.081 | 0.164 | 0.187 | -0.009 | 0.881 | -0.150 | 0.726 | 0.053 | 0.386 | 0.868 | 0.517 |
| Lamellar thickness 72(Left) | 0.109 | 0.061 | 0.204 | 0.140 | 0.020 | 2.349 | 0.122 | 0.049 | 0.418 | 0.811 | 0.712 |
| Lamellar thickness 72(Right) | 0.166 | 0.005 | 0.011 | 0.168 | 0.005 | 2.817 | 0.023 | 0.158 | 0.010 | 2.609 | 0.098 |
| Lamellar thickness 76(Left) | -0.091 | 0.117 | 0.277 | -0.023 | 0.696 | -0.391 | 0.797 | -0.017 | 0.784 | -0.274 | 0.867 |
| Lamellar thickness 76(Right) | 0.081 | 0.161 | 0.187 | 0.050 | 0.407 | 0.831 | 0.416 | 0.062 | 0.304 | 1.030 | 0.456 |
| Lamellar thickness 80(Left) | 0.035 | 0.547 | 0.656 | 0.083 | 0.165 | 1.391 | 0.412 | 0.068 | 0.258 | 1.133 | 0.586 |
| Lamellar thickness 80(Right) | 0.119 | 0.040 | 0.066 | 0.136 | 0.022 | 2.307 | 0.060 | 0.129 | 0.032 | 2.153 | 0.133 |
| Lamellar thickness 84(Left) | 0.124 | 0.032 | 0.148 | 0.153 | 0.010 | 2.585 | 0.095 | 0.080 | 0.190 | 1.315 | 0.564 |
| Lamellar thickness 84(Right) | 0.167 | 0.004 | 0.011 | 0.140 | 0.018 | 2.376 | 0.055 | 0.207 | 0.001 | 3.494 | 0.062 |
| Lamellar thickness 88(Left) | 0.038 | 0.513 | 0.631 | 0.054 | 0.373 | 0.893 | 0.603 | 0.075 | 0.221 | 1.227 | 0.573 |
| Lamellar thickness 88(Right) | 0.068 | 0.247 | 0.250 | 0.056 | 0.349 | 0.938 | 0.379 | 0.098 | 0.107 | 1.617 | 0.212 |

| | | | | | | | | | | | |
|---|---|---|---|---|---|---|---|---|---|---|---|
| Lamellar thickness 92(Left) | 0.117 | 0.044 | 0.185 | 0.104 | 0.081 | 1.749 | 0.264 | 0.035 | 0.566 | 0.575 | 0.801 |
| Lamellar thickness 92(Right) | 0.137 | 0.017 | 0.036 | 0.174 | 0.003 | 2.982 | 0.019 | 0.168 | 0.005 | 2.820 | 0.098 |
| Lamellar thickness 96(Left) | 0.079 | 0.174 | 0.348 | 0.079 | 0.187 | 1.322 | 0.427 | 0.083 | 0.174 | 1.365 | 0.563 |
| Lamellar thickness 96(Right) | 0.113 | 0.051 | 0.079 | 0.137 | 0.021 | 2.317 | 0.060 | 0.130 | 0.031 | 2.165 | 0.133 |
| Lamellar thickness 100(Left) | -0.029 | 0.616 | 0.680 | 0.005 | 0.939 | 0.076 | 0.918 | 0.010 | 0.873 | 0.161 | 0.914 |
| Lamellar thickness 100(Right) | 0.090 | 0.124 | 0.152 | 0.095 | 0.114 | 1.585 | 0.179 | 0.108 | 0.075 | 1.789 | 0.185 |
| Lamellar thickness 104(Left) | 0.089 | 0.128 | 0.286 | 0.039 | 0.515 | 0.653 | 0.673 | 0.060 | 0.326 | 0.985 | 0.631 |
| Lamellar thickness 104(Right) | 0.121 | 0.036 | 0.060 | 0.128 | 0.031 | 2.166 | 0.072 | 0.148 | 0.014 | 2.472 | 0.111 |
| Lamellar thickness 108(Left) | 0.072 | 0.227 | 0.426 | 0.023 | 0.705 | 0.379 | 0.797 | 0.056 | 0.363 | 0.911 | 0.649 |
| Lamellar thickness 108(Right) | 0.079 | 0.176 | 0.197 | 0.043 | 0.471 | 0.722 | 0.471 | 0.068 | 0.263 | 1.121 | 0.422 |
| Lamellar thickness 112(Left) | 0.080 | 0.170 | 0.348 | 0.019 | 0.752 | 0.316 | 0.831 | 0.067 | 0.269 | 1.108 | 0.586 |
| Lamellar thickness 112(Right) | 0.123 | 0.034 | 0.060 | 0.107 | 0.073 | 1.800 | 0.132 | 0.140 | 0.021 | 2.322 | 0.131 |
| Lamellar thickness 111(Left) | -0.019 | 0.743 | 0.751 | -0.042 | 0.487 | -0.696 | 0.666 | -0.002 | 0.973 | -0.034 | 0.970 |
| Lamellar thickness 111(Right) | 0.107 | 0.064 | 0.092 | 0.022 | 0.716 | 0.364 | 0.624 | 0.076 | 0.206 | 1.268 | 0.346 |
| Lamellar thickness 107(Left) | -0.007 | 0.909 | 0.831 | 0.011 | 0.855 | 0.183 | 0.873 | -0.019 | 0.754 | -0.314 | 0.867 |
| Lamellar thickness 107(Right) | 0.131 | 0.023 | 0.044 | 0.107 | 0.073 | 1.801 | 0.132 | 0.162 | 0.007 | 2.711 | 0.098 |
| Lamellar thickness 103(Left) | -0.065 | 0.268 | 0.468 | -0.066 | 0.274 | -1.095 | 0.529 | -0.091 | 0.135 | -1.500 | 0.563 |
| Lamellar thickness 103(Right) | 0.095 | 0.101 | 0.132 | 0.072 | 0.229 | 1.205 | 0.306 | 0.137 | 0.024 | 2.272 | 0.131 |
| Lamellar thickness 99(Left) | -0.053 | 0.364 | 0.511 | -0.040 | 0.502 | -0.672 | 0.666 | -0.087 | 0.157 | -1.420 | 0.563 |
| Lamellar thickness 99(Right) | 0.077 | 0.187 | 0.203 | 0.060 | 0.315 | 1.006 | 0.361 | 0.154 | 0.011 | 2.576 | 0.098 |
| Lamellar thickness 95(Left) | -0.028 | 0.623 | 0.680 | -0.034 | 0.574 | -0.563 | 0.724 | -0.059 | 0.328 | -0.980 | 0.631 |
| Lamellar thickness 95(Right) | 0.063 | 0.277 | 0.261 | 0.048 | 0.427 | 0.795 | 0.432 | 0.108 | 0.075 | 1.788 | 0.185 |
| Lamellar thickness 91(Left) | -0.094 | 0.105 | 0.264 | -0.075 | 0.210 | -1.256 | 0.446 | -0.059 | 0.330 | -0.976 | 0.631 |
| Lamellar thickness 91(Right) | 0.077 | 0.185 | 0.203 | 0.091 | 0.127 | 1.531 | 0.196 | 0.105 | 0.085 | 1.727 | 0.199 |
| Lamellar thickness 87(Left) | -0.055 | 0.343 | 0.511 | -0.046 | 0.446 | -0.763 | 0.639 | 0.010 | 0.864 | 0.172 | 0.914 |
| Lamellar thickness 87(Right) | 0.091 | 0.120 | 0.151 | 0.123 | 0.041 | 2.050 | 0.087 | 0.091 | 0.138 | 1.488 | 0.259 |
| Lamellar thickness 83(Left) | -0.056 | 0.330 | 0.511 | -0.052 | 0.380 | -0.880 | 0.603 | -0.020 | 0.742 | -0.329 | 0.867 |
| Lamellar thickness 83(Right) | 0.042 | 0.467 | 0.403 | 0.062 | 0.298 | 1.043 | 0.350 | 0.057 | 0.348 | 0.940 | 0.486 |
| Lamellar thickness 79(Left) | -0.020 | 0.728 | 0.751 | 0.018 | 0.766 | 0.298 | 0.837 | 0.014 | 0.816 | 0.233 | 0.880 |
| Lamellar thickness 79(Right) | 0.012 | 0.836 | 0.600 | 0.020 | 0.737 | 0.336 | 0.630 | 0.001 | 0.982 | 0.022 | 0.991 |
| Lamellar thickness 75(Left) | 0.039 | 0.497 | 0.628 | 0.061 | 0.308 | 1.022 | 0.562 | 0.059 | 0.333 | 0.971 | 0.631 |
| Lamellar thickness 75(Right) | 0.034 | 0.553 | 0.447 | 0.020 | 0.734 | 0.340 | 0.630 | -0.005 | 0.930 | -0.088 | 0.956 |

| | | | | | | | | | | | |
|---|---|---|---|---|---|---|---|---|---|---|---|
| Lamellar thickness 71(Left) | 0.017 | 0.765 | 0.765 | 0.012 | 0.843 | 0.199 | 0.873 | 0.001 | 0.987 | 0.017 | 0.970 |
| Lamellar thickness 71(Right) | 0.003 | 0.952 | 0.658 | 0.038 | 0.531 | 0.628 | 0.508 | 0.031 | 0.607 | 0.515 | 0.717 |
| Lamellar thickness 67(Left) | -0.005 | 0.938 | 0.835 | 0.030 | 0.610 | 0.511 | 0.743 | -0.016 | 0.788 | -0.270 | 0.867 |
| Lamellar thickness 67(Right) | 0.036 | 0.540 | 0.446 | 0.051 | 0.394 | 0.854 | 0.415 | 0.028 | 0.647 | 0.459 | 0.745 |
| Lamellar thickness 63(Left) | 0.033 | 0.564 | 0.661 | 0.068 | 0.251 | 1.150 | 0.497 | 0.029 | 0.633 | 0.478 | 0.833 |
| Lamellar thickness 63(Right) | -0.015 | 0.796 | 0.584 | 0.022 | 0.715 | 0.365 | 0.624 | -0.043 | 0.477 | -0.711 | 0.609 |
| Lamellar thickness 59(Left) | 0.005 | 0.929 | 0.835 | -0.006 | 0.918 | -0.103 | 0.918 | -0.017 | 0.774 | -0.287 | 0.867 |
| Lamellar thickness 59(Right) | -0.027 | 0.639 | 0.506 | 0.025 | 0.675 | 0.420 | 0.624 | -0.036 | 0.551 | -0.598 | 0.664 |
| Lamellar thickness 55(Left) | -0.015 | 0.799 | 0.790 | -0.013 | 0.831 | -0.214 | 0.872 | -0.040 | 0.515 | -0.651 | 0.749 |
| Lamellar thickness 55(Right) | 0.004 | 0.945 | 0.658 | 0.009 | 0.882 | 0.148 | 0.726 | -0.011 | 0.854 | -0.184 | 0.895 |
| Lamellar thickness 51(Left) | 0.029 | 0.614 | 0.680 | 0.016 | 0.796 | 0.259 | 0.852 | -0.019 | 0.755 | -0.312 | 0.867 |
| Lamellar thickness 51(Right) | 0.065 | 0.264 | 0.261 | 0.059 | 0.327 | 0.983 | 0.368 | 0.054 | 0.371 | 0.897 | 0.508 |
| Lamellar thickness 47(Left) | 0.021 | 0.715 | 0.751 | 0.037 | 0.537 | 0.618 | 0.695 | 0.007 | 0.905 | 0.120 | 0.930 |
| Lamellar thickness 47(Right) | 0.091 | 0.116 | 0.149 | 0.070 | 0.242 | 1.173 | 0.306 | 0.057 | 0.350 | 0.936 | 0.486 |
| Lamellar thickness 43(Left) | 0.053 | 0.363 | 0.511 | 0.076 | 0.204 | 1.275 | 0.440 | 0.076 | 0.212 | 1.250 | 0.573 |
| Lamellar thickness 43(Right) | 0.138 | 0.017 | 0.036 | 0.114 | 0.055 | 1.928 | 0.107 | 0.084 | 0.164 | 1.397 | 0.288 |
| Lamellar thickness 39(Left) | 0.102 | 0.080 | 0.246 | 0.130 | 0.029 | 2.196 | 0.153 | 0.123 | 0.042 | 2.039 | 0.298 |
| Lamellar thickness 39(Right) | 0.130 | 0.025 | 0.046 | 0.095 | 0.111 | 1.598 | 0.178 | 0.058 | 0.339 | 0.959 | 0.482 |
| Lamellar thickness 35(Left) | 0.115 | 0.048 | 0.186 | 0.130 | 0.030 | 2.180 | 0.153 | 0.133 | 0.028 | 2.204 | 0.238 |
| Lamellar thickness 35(Right) | 0.172 | 0.003 | 0.009 | 0.145 | 0.014 | 2.467 | 0.048 | 0.114 | 0.058 | 1.905 | 0.183 |
| Lamellar thickness 31(Left) | 0.111 | 0.056 | 0.201 | 0.142 | 0.017 | 2.407 | 0.114 | 0.143 | 0.018 | 2.372 | 0.223 |
| Lamellar thickness 31(Right) | 0.175 | 0.002 | 0.008 | 0.115 | 0.054 | 1.938 | 0.107 | 0.117 | 0.054 | 1.939 | 0.175 |
| Lamellar thickness 27(Left) | 0.115 | 0.048 | 0.186 | 0.150 | 0.012 | 2.526 | 0.099 | 0.137 | 0.024 | 2.264 | 0.230 |
| Lamellar thickness 27(Right) | 0.171 | 0.003 | 0.009 | 0.128 | 0.031 | 2.166 | 0.072 | 0.095 | 0.117 | 1.571 | 0.228 |
| Lamellar thickness 23(Left) | 0.101 | 0.087 | 0.249 | 0.157 | 0.009 | 2.635 | 0.095 | 0.104 | 0.090 | 1.700 | 0.493 |
| Lamellar thickness 23(Right) | 0.186 | 0.002 | 0.007 | 0.138 | 0.025 | 2.253 | 0.067 | 0.102 | 0.103 | 1.638 | 0.207 |
| Lamellar thickness 19(Left) | 0.078 | 0.178 | 0.349 | 0.082 | 0.171 | 1.373 | 0.412 | 0.046 | 0.452 | 0.753 | 0.717 |
| Lamellar thickness 19(Right) | 0.170 | 0.004 | 0.011 | 0.134 | 0.027 | 2.223 | 0.070 | 0.110 | 0.074 | 1.792 | 0.185 |
| Lamellar thickness 15(Left) | 0.046 | 0.436 | 0.565 | 0.054 | 0.378 | 0.883 | 0.603 | -0.003 | 0.956 | -0.055 | 0.970 |
| Lamellar thickness 15(Right) | 0.114 | 0.049 | 0.079 | 0.107 | 0.074 | 1.796 | 0.132 | 0.160 | 0.008 | 2.671 | 0.098 |
| Lamellar thickness 11(Left) | 0.055 | 0.347 | 0.511 | 0.049 | 0.421 | 0.805 | 0.629 | 0.008 | 0.897 | 0.129 | 0.930 |
| Lamellar thickness 11(Right) | 0.064 | 0.270 | 0.261 | 0.121 | 0.041 | 2.052 | 0.087 | 0.150 | 0.013 | 2.505 | 0.110 |

| | | | | | | | | | | | |
|---|---|---|---|---|---|---|---|---|---|---|---|
| Lamellar thickness 7(Left) | 0.066 | 0.276 | 0.473 | 0.049 | 0.427 | 0.795 | 0.629 | 0.012 | 0.843 | 0.198 | 0.901 |
| Lamellar thickness 7(Right) | 0.135 | 0.022 | 0.043 | 0.132 | 0.030 | 2.176 | 0.072 | 0.172 | 0.005 | 2.819 | 0.098 |
| Lamellar thickness 3(Left) | 0.165 | 0.005 | 0.029 | 0.113 | 0.061 | 1.883 | 0.213 | 0.137 | 0.025 | 2.250 | 0.230 |
| Lamellar thickness 3(Right) | 0.248 | 0.000 | 0.002 | 0.146 | 0.016 | 2.429 | 0.052 | 0.194 | 0.002 | 3.191 | 0.088 |

### References

1. Adler, D. H. *et al.* Characterizing the human hippocampus in aging and Alzheimer's disease using a computational atlas derived from ex vivo MRI and histology. *Proc. Natl. Acad. Sci. U.S.A.* **115**, 4252–4257 (2018).
2. Gross, D. W., Misaghi, E., Steve, T. A., Wilman, A. H. & Beaulieu, C. Curved multiplanar reformatting provides improved visualization of hippocampal anatomy. *Hippocampus* **30**, 156–161 (2020).
3. Gao, N., Ye, C., Chen, H., Hao, X. & Ma, T. MRI -based axis-referenced morphometric model corresponding to lamellar organization for assessing hippocampal atrophy in dementia. *Human Brain Mapping* **45**, e26715 (2024).
4. Di Martino, A. *et al.* The autism brain imaging data exchange: towards a large-scale evaluation of the intrinsic brain architecture in autism. *Mol Psychiatry* **19**, 659–667 (2014).
5. Consortium, T. The ADHD-200 Consortium: a model to advance the translational potential of neuroimaging in clinical neuroscience. *Front. Syst. Neurosci.* **6**, (2012).
6. Shafto, M. A. *et al.* The Cambridge Centre for Ageing and Neuroscience (Cam-CAN) study protocol: a cross-sectional, lifespan, multidisciplinary examination of healthy cognitive ageing. *BMC Neurology* **14**, 204 (2014).
7. Taylor, J. R. *et al.* The Cambridge Centre for Ageing and Neuroscience (Cam-CAN) data repository: Structural and functional MRI, MEG, and cognitive data from a cross-sectional adult lifespan sample. *NeuroImage* **144**, 262–269 (2017).

8. Zuo, X.-N. *et al.* An open science resource for establishing reliability and reproducibility in functional connectomics. *Sci Data* **1**, 140049 (2014).
9. Hanlon, F. M. *et al.* Bilateral hippocampal dysfunction in schizophrenia. *NeuroImage* **58**, 1158–1168 (2011).
10. Mayer, A. R. *et al.* Functional imaging of the hemodynamic sensory gating response in schizophrenia: fMRI and Gating in Schizophrenia. *Hum. Brain Mapp* **34**, 2302–2312 (2013).
11. Biswal, B. B. *et al.* Toward discovery science of human brain function. *Proc. Natl. Acad. Sci. U.S.A.* **107**, 4734–4739 (2010).
12. Alexander, L. M. *et al.* An open resource for transdiagnostic research in pediatric mental health and learning disorders. *Sci Data* **4**, 170181 (2017).
13. Tobe, R. H. *et al.* A longitudinal resource for studying connectome development and its psychiatric associations during childhood. *Sci Data* **9**, 300 (2022).
14. Frontiers | The NKI-Rockland Sample: A Model for Accelerating the Pace of Discovery Science in Psychiatry. https://www.frontiersin.org/journals/neuroscience/articles/10.3389/fnins.2012.00152/full.
15. Wei, D. *et al.* Structural and functional brain scans from the cross-sectional Southwest University adult lifespan dataset. *Sci Data* **5**, 180134 (2018).
16. Liu, W. *et al.* Longitudinal test-retest neuroimaging data from healthy young adults in southwest China. *Sci Data* **4**, 170017 (2017).

17. Nooner, K. B. *et al.* The NKI-Rockland Sample: A Model for Accelerating the Pace of Discovery Science in Psychiatry. *Front Neurosci* **6**, 152 (2012).
18. Tanaka, S. C. *et al.* A multi-site, multi-disorder resting-state magnetic resonance image database. *Sci Data* **8**, 227 (2021).
19. Van Essen, D. C. *et al.* The Human Connectome Project: a data acquisition perspective. *Neuroimage* **62**, 2222–2231 (2012).
20. Bookheimer, S. Y. *et al.* The Lifespan Human Connectome Project in Aging: An overview. *NeuroImage* **185**, 335–348 (2019).
21. Somerville, L. H. *et al.* The Lifespan Human Connectome Project in Development: A large-scale study of brain connectivity development in 5–21 year olds. *NeuroImage* **183**, 456–468 (2018).
22. Mueller, S. G. *et al.* The Alzheimer's disease neuroimaging initiative. *Neuroimaging Clinics* **15**, 869–877 (2005).
23. Ellis, K. A. *et al.* The Australian Imaging, Biomarkers and Lifestyle (AIBL) study of aging: methodology and baseline characteristics of 1112 individuals recruited for a longitudinal study of Alzheimer's disease. *Int Psychogeriatr* **21**, 672–687 (2009).
24. Buckner, R. L., Roffman, J. L., Smoller, J. W., & Neuroinformatics Research Group. Brain Genomics Superstruct Project (GSP). 10891991040, 10972774400, 10798878720, 10438246400, 10689955840, 11055155200, 10736506880, 10900736000, 10677258240, 10844508160, 105135, 681486, 380, 2423138, 2423049, 41902, 8960821248, 651720, 289445, 2435962 Harvard Dataverse https://doi.org/10.7910/DVN/25833 (2014).
25. LaMontagne, P. J. *et al.* OASIS-3: Longitudinal Neuroimaging, Clinical, and Cognitive Dataset for Normal Aging and Alzheimer Disease. 2019.12.13.19014902 Preprint at https://doi.org/10.1101/2019.12.13.19014902 (2019).

26. Jernigan, T. L. *et al.* The Pediatric Imaging, Neurocognition, and Genetics (PING) Data Repository. *Neuroimage* **124**, 1149–1154 (2016).

27. Satterthwaite, T. D. *et al.* The Philadelphia Neurodevelopmental Cohort: A publicly available resource for the study of normal and abnormal brain development in youth. *NeuroImage* **124**, 1115–1119 (2016).

28. Marek, K. *et al.* The Parkinson Progression Marker Initiative (PPMI). *Progress in Neurobiology* **95**, 629–635 (2011).

29. Chopra, S. *et al.* The Transdiagnostic Connectome Project: an open dataset for studying brain-behavior relationships in psychiatry. *Sci Data* **12**, 923 (2025).

30. Momenian, M., Zhengwu Ma, Shuyi Wu, Chengcheng Wang, & Jixing Li. Le Petit Prince Hong Kong: Naturalistic fMRI and EEG dataset from older Cantonese speakers. Openneuro https://doi.org/10.18112/OPENNEURO.DS004718.V1.1.2 (2025).

31. Dobrushina, O. R. *et al.* Modulation of Intrinsic Brain Connectivity by Implicit Electroencephalographic Neurofeedback. *Front. Hum. Neurosci.* **14**, (2020).

32. Poldrack, R. A. *et al.* A phenome-wide examination of neural and cognitive function. *Sci Data* **3**, 160110 (2016).

33. Gold, C. E. *Exploring the Resting State Neural Activity of Monolinguals and Late and Early Bilinguals*. (Brigham Young University, 2018).

34. Bilingualism and the brain - OpenNeuro. https://openneuro.org/datasets/ds001796/versions/1.5.0/file-display/CHANGES.

35. Sunavsky, A. & Poppenk, J. Neuroimaging predictors of creativity in healthy adults. *NeuroImage* **206**, 116292 (2020).

36. The Amsterdam Open MRI Collection, a set of multimodal MRI datasets for individual difference analyses | Scientific Data. https://www.nature.com/articles/s41597-021-00870-6.

37. Garza-Villarreal, E. A. *et al.* Clinical and Functional Connectivity Outcomes of 5-Hz Repetitive Transcranial Magnetic Stimulation as an Add-on Treatment in Cocaine Use Disorder: A Double-Blind Randomized Controlled Trial. *Biological Psychiatry: Cognitive Neuroscience and Neuroimaging* **6**, 745–757 (2021).
38. Rasgado-Toledo, J., Shah, A., Ingalhalikar, M. & Garza-Villarreal, E. A. Neurite orientation dispersion and density imaging in cocaine use disorder. *Progress in Neuro-Psychopharmacology and Biological Psychiatry* **113**, 110474 (2022).
39. Pechenkova, E. V. *et al.* Speech disfluencies: Neurophysiological aspect in normal population. Openneuro https://doi.org/10.18112/OPENNEURO.DS003469.V1.0.0 (2021).
40. Cognitive Control Theoretic Mechanisms of Real-time fMRI-Guided Neuromodulation (CTM) - OpenNeuro. https://openneuro.org/datasets/ds003831/versions/1.0.0/file-display/task-modulate2_bold.json.
41. Hsu, C.-T., Clariana, R., Schloss, B. & Li, P. Neurocognitive Signatures of Naturalistic Reading of Scientific Texts: A Fixation-Related fMRI Study. *Sci Rep* **9**, 10678 (2019).
42. Follmer, D. J., Fang, S.-Y., Clariana, R. B., Meyer, B. J. F. & Li, P. What predicts adult readers' understanding of STEM texts? *Read Writ* **31**, 185–214 (2018).
43. Li, P. & Clariana, R. B. Reading comprehension in L1 and L2: An integrative approach. *Journal of Neurolinguistics* **50**, 94–105 (2019).
44. Sinclair, B. *et al.* Heritability of the network architecture of intrinsic brain functional connectivity. *NeuroImage* **121**, 243–252 (2015).

45. Functional correlates of cognitive performance and working memory in temporal lobe epilepsy: Insights from task-based and resting-state fMRI | PLOS One. https://journals.plos.org/plosone/article?id=10.1371/journal.pone.0295142.

46. Bissett, P. G. *et al.* Cognitive tasks, anatomical MRI, and functional MRI data evaluating the construct of self-regulation. *Sci Data* **11**, 809 (2024).

47. Miranda-Angulo, A. L. *et al.* Sympathovagal quotient and resting-state functional connectivity of control networks are related to gut *Ruminococcaceae* abundance in healthy men. *Psychoneuroendocrinology* **164**, 107003 (2024).

48. Food and Brain Study - OpenNeuro. https://openneuro.org/datasets/ds004697/versions/1.0.2.

49. López-Caballero, F., Curtis, M., Coffman, B. A. & Salisbury, D. F. Is source-resolved magnetoencephalographic mismatch negativity a viable biomarker for early psychosis? *Eur J of Neuroscience* **59**, 1889–1906 (2024).

50. Botvinik-Nezer, R. *et al.* Placebo treatment affects brain systems related to affective and cognitive processes, but not nociceptive pain. Preprint at https://doi.org/10.1101/2023.09.21.558825 (2023).

51. Horta, M., Polk, R. & Ebner, N. Single Dose Intranasal Oxytocin Administration: Data from Healthy Younger and Older Adults. Openneuro https://doi.org/10.18112/OPENNEURO.DS004725.V1.0.1 (2023).

52. Tisdall, L. & Mata, R. AgeRisk. Openneuro https://doi.org/10.18112/OPENNEURO.DS004711.V1.0.0 (2023).

53. Rovai, A., Lolli, V., Trotta, N., Goldman, S. & De Tiège, X. Cerebrovascular Reactivity Normative Dataset. Openneuro https://doi.org/10.18112/OPENNEURO.DS004604.V2.0.0 (2024).

54. Goulding, L., Schmidt, A., Hamm, I., & C. Brock Kirwan. fMRI Investigations of Individual Differences on Memory Activation for Faces and Words: The Effects of Handedness and Phenomenal Experience. Openneuro https://doi.org/10.18112/OPENNEURO.DS004589.V1.0.0 (2023).

55. Grössinger, D. *et al.* The role of superstition of cognitive control during neurofeedback training. Preprint at https://doi.org/10.1101/2021.09.14.460252 (2021).

56. Gibson, M. *et al.* The Aphasia Recovery Cohort, an open-source chronic stroke repository. *Sci Data* **11**, 981 (2024).

57. Racey, C. *et al.* An Open Science MRI Database of over 100 Synaesthetic Brains and Accompanying Deep Phenotypic Information. *Sci Data* **10**, 766 (2023).

58. Frank, L. E. & Zeithamova, D. Evaluating methods for measuring background connectivity in slow event-related fMRI designs. Openneuro https://doi.org/10.18112/OPENNEURO.DS004349.V1.0.0 (2022).

59. Soler-Vidal, J. *et al.* Brain correlates of speech perception in schizophrenia patients with and without auditory hallucinations. *PLoS ONE* **17**, e0276975 (2022).

60. Rogers, C. S., Jones, M. S., McConkey, S. & Peelle, J. E. Listening task. Openneuro https://doi.org/10.18112/OPENNEURO.DS004285.V1.0.0 (2022).

61. TODO:, First1 Last1, First2 Last2, & ... Cross-stage neural pattern similarity in the hippocampus predicts false memory derived from post-event inaccurate information. Openneuro https://doi.org/10.18112/OPENNEURO.DS004261.V2.0.0 (2023).

62. Shao, X., Li, A., Chen, C., Loftus, E. F. & Zhu, B. Cross-stage neural pattern similarity in the hippocampus predicts false memory derived from post-event inaccurate information. *Nat Commun* **14**, 2299 (2023).
63. Thornton, M. A., Barrick, E., Liang, N. & Tamir, D. I. Interacting representations of mental states and traits. Openneuro https://doi.org/10.18112/OPENNEURO.DS004217.V1.0.0 (2022).
64. Nugent, A. C. *et al.* The NIMH intramural healthy volunteer dataset: A comprehensive MEG, MRI, and behavioral resource. *Sci Data* **9**, 518 (2022).
65. Schuch, F. *et al.* An open presurgery MRI dataset of people with epilepsy and focal cortical dysplasia type II. Openneuro https://doi.org/10.18112/OPENNEURO.DS004199.V1.0.6 (2025).
66. Nárai, Á. *et al.* Movement-related artefacts (MR-ART) dataset of matched motion-corrupted and clean structural MRI brain scans. *Sci Data* **9**, 630 (2022).
67. Strike, L. T. *et al.* The Queensland Twin Adolescent Brain Project, a longitudinal study of adolescent brain development. *Sci Data* **10**, 195 (2023).
68. Ma, S. *et al.* WholebodySomatotopicMapping. Openneuro https://doi.org/10.18112/OPENNEURO.DS004044.V2.0.3 (2022).
69. Turesky, T. *et al.* BrainMorphometry_DiminishedGrowth_BEANstudy_2021. Openneuro https://doi.org/10.18112/OPENNEURO.DS003877.V1.2.0 (2026).

70. Zareba, M. R. *et al.* Late chronotype is linked to greater cortical thickness in the left fusiform and entorhinal gyri. *Biological Rhythm Research* **53**, 1626–1638 (2022).

71. Kliemann, D. *et al.* Caltech Conte Center - A multimodal data resource for exploring social cognition and decision-making. Openneuro https://doi.org/10.18112/OPENNEURO.DS003798.V1.0.5 (2021).

72. Smith, D. V., Ludwig, R. M., Dennison, J. B., Reeck, C. & Fareri, D. S. An fMRI Dataset on Social Reward Processing and Decision Making in Younger and Older Adults. Preprint at https://doi.org/10.31234/osf.io/k7d56 (2023).

73. Fareri, D. S. *et al.* Age-related differences in ventral striatal and default mode network function during reciprocated trust. *NeuroImage* **256**, 119267 (2022).

74. Peelle, J. E. *et al.* Increased Connectivity among Sensory and Motor Regions during Visual and Audiovisual Speech Perception. *J. Neurosci.* **42**, 435–442 (2022).

75. Keren, H. *et al.* The temporal representation of experience in subjective mood. *eLife* **10**, e62051 (2021).

76. Ito, T. *et al.* Cognitive task information is transferred between brain regions via resting-state network topology. *Nat Commun* **8**, 1027 (2017).

77. Li, J. *et al.* Le Petit Prince multilingual naturalistic fMRI corpus. *Sci Data* **9**, 530 (2022).

78. Crotti, M., Koschutnig, K. & Wriessnegger, S. The impact of handedness on the neural correlates during kinesthetic motor imagery: a FMRI study. Openneuro https://doi.org/10.18112/OPENNEURO.DS003612.V1.0.5 (2025).

79. R. Nathan Spreng *et al.* Neurocognitive aging data release with behavioral, structural, and multi-echo functional MRI measures. Openneuro https://doi.org/10.18112/OPENNEURO.DS003592.V1.0.13 (2022).

80. Liuzzi, L. *et al.* Magnetoencephalographic correlates of mood and reward dynamics in human adolescents. *Cerebral Cortex* **32**, 3318–3330 (2022).

81. Novén, M. *et al.* Language Learning Aptitude dataset. Openneuro https://doi.org/10.18112/OPENNEURO.DS003508.V1.0.0 (2021).

82. Nussenbaum, K. & Hartley, C. A. Developmental change in prefrontal cortex recruitment supports the emergence of value-guided memory. Openneuro https://doi.org/10.18112/OPENNEURO.DS003499.V1.0.1 (2021).

83. Rasgado-Toledo, J. *et al.* A Dataset to Study Pragmatic Language and Its Underlying Cognitive Processes. *Front. Hum. Neurosci.* **15**, 666210 (2021).

84. Arbula, S., Pisanu, E. & Rumiati, R. I. Agreeableness personality trait and social information encoding. Openneuro https://doi.org/10.18112/OPENNEURO.DS003436.V1.0.0 (2020).

85. Cai, L. Y. *et al.* MASiVar: Multisite, Multiscanner, and Multisubject Acquisitions for Studying Variability in Diffusion Weighted Magnetic Resonance Imaging. Openneuro https://doi.org/10.18112/OPENNEURO.DS003416.V2.0.2 (2021).

86. Tomova, L. *et al.* MRI data of 40 adult participants in response to a cue induced craving task following food fasting, social isolation and baseline (within-subject design). Openneuro https://doi.org/10.18112/OPENNEURO.DS003242.V1.0.0 (2020).

87. ..., Koschutnig, K., Weber, B. & Fink, A. Tidying Up White Matter: Neuroplastic Transformations in Sensorimotor Tracts following Slackline Skill Acquisition. Openneuro https://doi.org/10.18112/OPENNEURO.DS003138.V1.0.1 (2023).

88. Banfi, C. *et al.* Reading-related functional activity in children with isolated spelling deficits and dyslexia. *Language, Cognition and Neuroscience* **36**, 543–561 (2021).

89. Snoek, L. *et al.* The Amsterdam Open MRI Collection, a set of multimodal MRI datasets for individual difference analyses. *Sci Data* **8**, 85 (2021).

90. Booth, J. R. *et al.* Brain Development of Deductive Reasoning. Openneuro https://doi.org/10.18112/OPENNEURO.DS002886.V1.1.0 (2021).

91. Lytle, M. N., Prado, J. & Booth, J. R. A neuroimaging dataset of deductive reasoning in school-aged children. *Data in Brief* **33**, 106405 (2020).

92. Sangil Lee & Kable, J. Cognitive Training. Openneuro https://doi.org/10.18112/OPENNEURO.DS002843.V1.0.1 (2021).

93. Aliko, S., Huang, J., Gheorghiu, F., Meliss, S. & Skipper, J. I. A naturalistic neuroimaging database for understanding the brain using ecological stimuli. *Sci Data* **7**, 347 (2020).

94. Zhu, B. *et al.* Multiple interactive memory representations underlie the induction of false memory. Openneuro https://doi.org/10.18112/OPENNEURO.DS002731.V1.0.2 (2021).

95. Rozenkrantz, L. *et al.* Unexplained Repeated Pregnancy Loss is Associated with Altered Perceptual and Brain Responses to Men's Body-Odor. Openneuro https://doi.org/10.18112/OPENNEURO.DS002717.V1.0.1 (2020).

96. Mather, M. *et al.* Isometric exercise facilitates attention to salient events in women via the noradrenergic system. *NeuroImage* **210**, 116560 (2020).

97. Van Der Laan, L. N., Scholz, C., Poldrack, R. A., De Ridder, D. T. D. & Smidts, A. Can we have a second helping? A replication study on the neurobiological mechanisms underlying self-control. Openneuro https://doi.org/10.18112/OPENNEURO.DS002643.V1.1.0 (2022).

98. Booth, J. R. *et al.* Working Memory and Reward in Children with and without Attention Deficit Hyperactivity Disorder (ADHD). Openneuro https://doi.org/10.18112/OPENNEURO.DS002424.V1.2.0 (2021).

99. Lytle, M. N., Hammer, R. & Booth, J. R. A neuroimaging dataset on working memory and reward processing in children with and without ADHD. *Data in Brief* **31**, 105801 (2020).

100. Luna, B. PETfrog. Openneuro https://doi.org/10.18112/OPENNEURO.DS002385.V1.1.0 (2021).

101. Rogers, C. S. *et al.* Age-Related Differences in Auditory Cortex Activity During Spoken Word Recognition. *Neurobiology of Language* **1**, 452–473 (2020).

102. Nastase, S. A. *et al.* Narratives. Openneuro https://doi.org/10.18112/OPENNEURO.DS002345.V1.1.4 (2020).

103. Lytle, M. N., Bitan, T. & Booth, J. R. A neuroimaging dataset on orthographic, phonological and semantic word processing in school-aged children. *Data in Brief* **28**, 105091 (2020).

104. Bigio, J. *et al.* Cross-Sectional Multidomain Lexical Processing. Openneuro https://doi.org/10.18112/OPENNEURO.DS002236.V1.1.1 (2022).

105. Goffin, C., Slipenkyj, M., Vogel, S. E., Merkley, R. & Ansari, D. Development of Symbolic Number Processing. Openneuro https://doi.org/10.18112/OPENNEURO.DS002116.V1.0.0 (2019).

106. Lytle, M. N., McNorgan, C. & Booth, J. R. A longitudinal neuroimaging dataset on multisensory lexical processing in school-aged children. *Sci Data* **6**, 329 (2019).

107. Booth, J. R. *et al.* Longitudinal Brain Correlates of Multisensory Lexical Processing in Children. Openneuro https://doi.org/10.18112/OPENNEURO.DS001894.V1.4.2 (2022).

108. H. Moriah Sokolowski, Hawes, Z., Peters, L. & Ansari, D. Parallel Adaptation of Symbols, Quantities, and Physical Size. Openneuro https://doi.org/10.18112/OPENNEURO.DS001848.V1.0.1 (2019).

109. Goffin, C., Sokolowski, H. M., Slipenkyj, M. & Ansari, D. Does writing handedness affect neural representation of symbolic number? An fMRI adaptation study. *Cortex* **121**, 27–43 (2019).

110. Fynes-Clinton, S., Marstaller, L. & Burianová, H. Differentiation of functional networks during long-term memory retrieval in children and adolescents. *NeuroImage* **191**, 93–103 (2019).

111. Suárez-Pellicioni, M., Lytle, M., Younger, J. W. & Booth, J. R. A longitudinal neuroimaging dataset on arithmetic processing in school children. *Sci Data* **6**, 190040 (2019).

112. Botvinik-Nezer, R. *et al.* fMRI data of mixed gambles from the Neuroimaging Analysis Replication and Prediction Study. *Sci Data* **6**, 106 (2019).

113. Berteletti, I. *et al.* Brain Correlates of Math Development. Openneuro https://doi.org/10.18112/OPENNEURO.DS001486.V1.3.1 (2021).

114. Yeshurun, Y., Nguyen, M. & Hasson, U. Amplification of local changes along the timescale processing hierarchy. *Proc. Natl. Acad. Sci. U.S.A.* **114**, 9475–9480 (2017).

115. Petersen, S., Schlaggar, B. & Power, J. Washington University 120. Openneuro https://doi.org/10.18112/OPENNEURO.DS000243.V1.0.0 (2025).

116. Galiano, A. *et al.* Resting State Perfusion in Healthy Aging. Openneuro https://doi.org/10.18112/OPENNEURO.DS000240.V2.0.0 (2021).

117. Richardson, H., Lisandrelli, G., Riobueno-Naylor, A. & Saxe, R. MRI data of 3-12 year old children and adults during viewing of a short animated film. Openneuro https://doi.org/10.18112/OPENNEURO.DS000228.V1.1.1 (2023).

118. FitzGerald, T. H. B., Haemmerer, D., Friston, K. J., Li, S.-C. & Dolan, R. J. Sequential Inference VBM. Openneuro https://doi.org/10.18112/OPENNEURO.DS000222.V1.0.1 (2019).

119. Tetreault P *et al.* Brain connectivity predicts placebo response across chronic pain clinical trials. Openneuro https://doi.org/10.18112/OPENNEURO.DS000208.V1.0.1 (2022).

120. Velanova, K., Wheeler, M. E. & Luna, B. Maturational Changes in Anterior Cingulate and Frontoparietal Recruitment Support the Development of Error Processing and Inhibitory Control. *Cerebral Cortex* **18**, 2505–2522 (2008).
